# An immune cell lipid atlas reveals the basis of susceptibility to ferroptosis

**DOI:** 10.1101/2023.02.10.528075

**Authors:** Gerard Pernes, Pooranee K Morgan, Kevin Huynh, Corey Giles, Sudip Paul, Adam Alexander T Smith, Natalie A Mellett, Camilla Bertuzzo Veiga, Thomas JC Collins, T Michael De Silva, Man KS Lee, Peter J Meikle, Graeme I Lancaster, Andrew J Murphy

**Author notes:** Address for correspondence: Dr Graeme I Lancaster,; Prof. Andrew J. Murphy. These authors contributed equally (Gerard Pernes, Pooranee K Morgan). Senior Author (Graeme I Lancaster, Andrew J Murphy).

## Abstract

The cellular lipidome is comprised of thousands of unique lipid species. This complexity underpins the many roles of lipids in cellular biology. How lipidome composition varies between cell types and how such differences contribute to cell-specific functionality is poorly understood. Here, using mass spectrometry-based targeted lipidomics, we have characterised the cellular lipid landscape of the human and mouse immune systems (www.cellularlipidatlas.com). We find that myeloid and lymphoid cell lineages have unique lipid compositions, notably in the usage of ester and ether bonds within glycerophospholipids (PLs) and PL acyl chain composition. To determine if immune cell-specific lipid phenotypes promote cell-specific functional properties we focused on differences in poly-unsaturated fatty acid (PUFA)-containing PL, the levels of which are markedly higher in lymphoid cells relative to myeloid cells. We firstly show that differences in PUFA-PL content provides a mechanistic basis for previously described differences in immune cell susceptibility to ferroptosis, a form of cell death driven by iron-dependent lipid peroxidation, and secondly, that the low PUFA-PL content of neutrophils restrains NADPH oxidase-driven ferroptosis. In summary, we show that the lipid landscape is a defining feature of immune cell identity and that cell-specific lipid phenotypes underpin aspects of immune cell physiology.

## Introduction

Lipids are essential cellular components with roles in multiple aspects of cell biology (Harayama and Riezman, 2018). A fundamental property of lipids is their capacity to self-associate and form bilayers. This feature enables the formation of membranes, allowing cells to segregate their internal constituents from the external environment and, in eukaryotes, to form sub-cellular organelles (Holthuis and Menon, 2014). The varied biophysical properties of lipids enable membranes to alter their shape, thickness, and order (Harayama and Riezman, 2018). These properties are essential for fission, fusion, budding, tubulation, and the formation of lipid rafts (Harayama et al., 2014). These membrane features underlie essential cellular processes such as cell division, mitochondrial dynamics, endocytosis, protein export, and cell signalling (Sezgin et al., 2017; Storck et al., 2018; Walpole and Grinstein, 2020; Zhang et al., 2014). Lipids also have a multitude of biochemical roles, including being harbingers of cell death, for example phosphatidylserine (PS) in apoptosis (Morioka et al., 2019) and polyunsaturated fatty acids (PUFAs) in ferroptosis (Jiang et al., 2021). Lipids act as signalling molecules and contribute to post-translational modifications (e.g. myristylation, palmitoylation) that influence protein localisation, activity, and stability (Chen et al., 2018). Finally, lipids are substrates in energy metabolism and heat production (Verkerke and Kajimura, 2021). The ability of lipids to function in such wide-ranging aspects of cellular biology is due to both the chemical complexity of the lipidome, with ∼100,000 theoretical lipid species, and the compositional diversity of the lipidome, which varies between tissues, cell types, organelles, and membrane leaflets (inner *vs* outer bilayer) (Harayama and Riezman, 2018). The importance of maintaining lipid homeostasis for cellular and organismal fitness is underscored by the many diseases that arise as a result of dysregulated lipid metabolism.

While thousands of chemically diverse lipid species are present within cells, the structural differences between many lipid species are relatively small, e.g. the position and number of double bonds, the types of chemical linkage used to attach aliphatic chains to backbones, and acyl chain length. While significant progress has been made in both describing lipid structural diversity and determining functions of particular lipid classes and species, we currently have a limited understanding of how lipid diversity, and the compositional variance in the cellular lipidome it encodes, affects cellular biology in general, but also how it may contribute to cell-specific functionality. In this regard, a limitation of our current knowledge is how lipid composition varies between cell types. The value of such information is highlighted by recent efforts describing the metabolic landscape of 928 cancer cell lines, where lipidome compositional variance between cell lines led to the identification of therapeutically exploitable lipid profiles (Li et al., 2019).

While compositional differences in tissue lipidomes have been described in mice (Surma et al., 2021), parallel lipidomic analysis in human tissues is challenging. Furthermore, cellular heterogeneity within and between tissues limits the interpretation of lipid composition data obtained from whole tissues. The mammalian immune system is comprised of a multitude of highly specialized cells with diverse, and often unique, functional properties, and therefore provides an opportunity to explore how lipid composition may vary between cell types with cell-specific effector functions. Indeed, previous studies have described marked compositional variance in the lipidomes of a limited number of specific immune cell types (Alarcon-Barrera et al., 2020).

We hypothesized that the cellular lipidome would be a defining feature of immune cell identity and that differences in lipid composition between immune cells would contribute to cell-specific functionality. Herein, we have used mass spectrometry-based lipidomics to comprehensively describe the cellular lipidome of the human and mouse immune systems. Our immune cell lipid atlas comprises >500 individual lipid species measured in 16 human and 8 murine mature immune cell types and is freely available to the community (www.cellularlipidatlas.com). Using this resource, we have identified how cell-specific lipidomic phenotypes govern the susceptibility of immune cells to ferroptotic cell death.

## Results

### A lipidomic cell atlas of the human immune system

To test the hypothesis that the cellular lipidome would be a defining feature of the cells of the human immune system we isolated 16 distinct mature immune cell populations, representing all of the major immune cell lineages and their subsets from the peripheral blood of 14 healthy donors. We then characterized their cellular lipidomes by ultra-high-performance liquid chromatography coupled to tandem mass spectrometry (UHPLC ESI-MS/MS) (Fig. 1a). We quantified 524 individual lipid species, 505 of which were significantly different between at least 2 cell types, belonging to multiple distinct lipid classes, including diacyl glycerophospholipids (PLs) (glycerophosphocholine [PC], glycerophosphoethanolamine [PE], glycerophosphoserine [PS], glycerophosphoinositol [PI], glycerophosphoglycerol [PG], glycerophosphate [PA]), sphingolipids (SLs) (sphingomyelin [SM], ceramide, ganglioside, hexosylceramide), lyso-PLs, ether- and vinyl-ether PLs, and cholesterol. In the present study we were specifically interested in membrane lipids, therefore neutral lipids, e.g. triacylglycerol (TG) species, diacylglycerol (DG) species, and cholesterol esters (CE), which are abundant components of lipid droplets, were not analysed.

**Figure 1.**
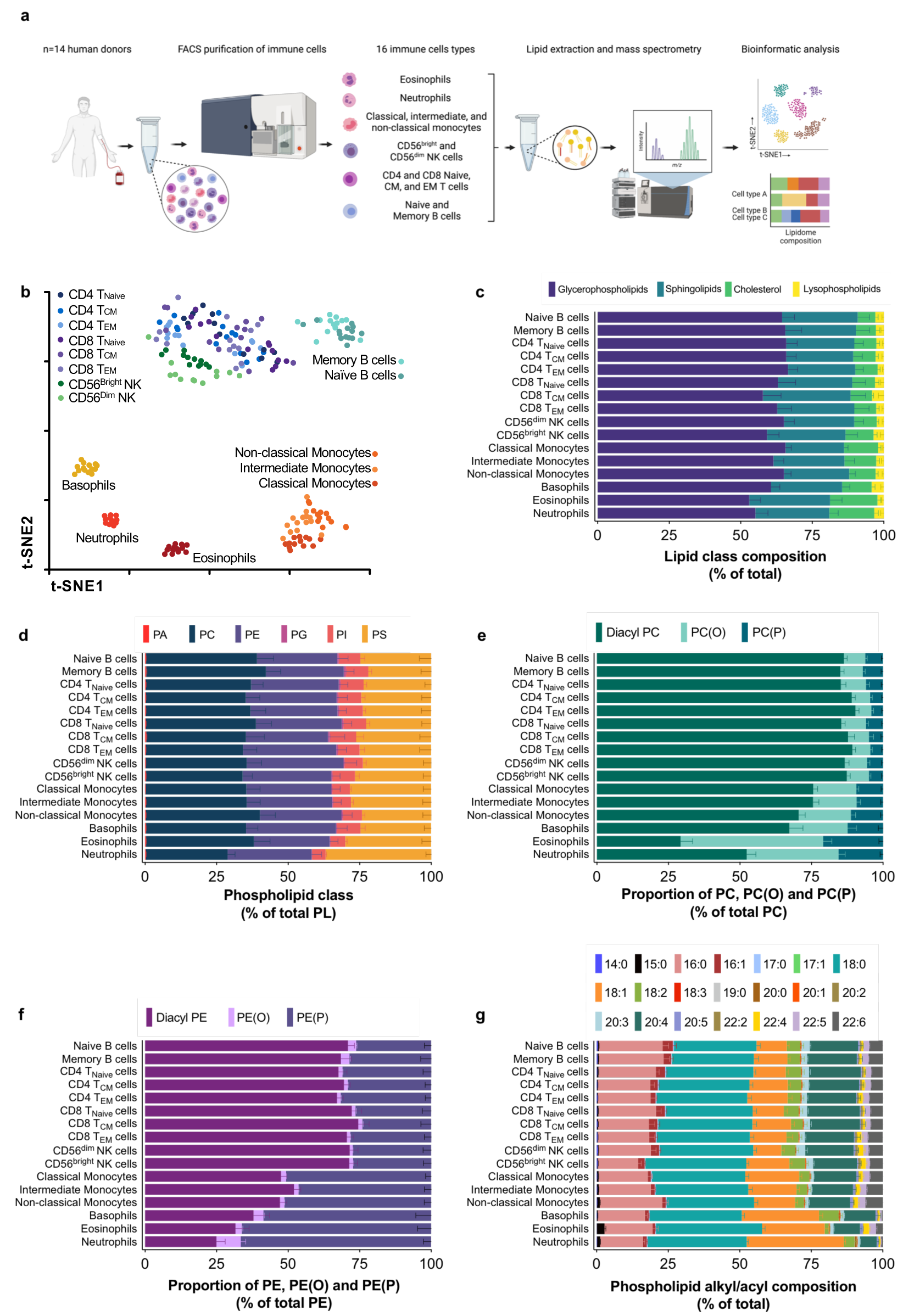
A lipidomic cell atlas of the human immune system. **a,** Workflow schematic. **b,** t-SNE map displaying the global cellular lipidome of the human immune system. Dots are individual donors; n=11-14 individual donors for each cell type. T_EM_, T_Effector Memory_; T_CM_, T_Central Memory_; NK, Natural Killer. **c-g,** Breakdown of lipid features in human immune cells by class (**c**), phospholipid headgroup type (**d**), type of Sn1 chemical linkage in PC (**e**) and PE (**f**), and PL alkyl/acyl composition (**g**). Data are shown as mean + S.D from n=11-14 individual donors. In each figure, statistically significant differences in specific lipid features between cell types was determined by 1-way ANOVA with Tukey’s HSD test after false discovery rate correction (5%; Benjamini-Hochberg). P values for all pairwise comparisons are shown in Supplementary Excel file 1.

t-Distributed stochastic neighbour embedding (t-SNE) was performed to visualize the global lipidome of immune cells (Fig. 1b). t-SNE resolved 6 distinct cellular clusters: (1) neutrophils, (2) eosinophils, (3) basophils, (4) monocytes, (5) B cells, and (6) T and NK cells. Furthermore, within the B cell cluster a distinction was apparent between memory and naïve B cells, while within monocytes, classical and non-classical monocytes clustered distinctly (Fig. 1b). Within T lymphocytes and NK cells, T cell subsets did not form distinct clusters; however, NK cells tended to form a distinct sub-cluster (Fig. 1b). These data demonstrate that the global cellular lipidome is an identifying feature of the cells of the human immune system.

### The cells of the human immune system have distinct profiles of phospholipid composition

To understand the primary drivers of the global lipidome differences within the human immune system, we examined how various chemical features of lipids varied between immune cells. Exploring the major lipid classes in mammalian cell membranes, PLs, SLs, lyso-PLs (PLs containing a single acyl chain) and cholesterol, revealed a generally similar profile across all cells (Fig. 1c; significance values for all human lipid atlas data are shown in Supplementary Excel file 1). However, we noted the proportion of SL was higher in myeloid cells, notably in neutrophils and eosinophils, with a corresponding decrease in the proportion of PL (Fig. 1c).

PLs are the most abundant lipid class present in cell membranes (Holthuis and Menon, 2014) (Fig. 1c) and are defined by their variable head group (e.g. phosphocholine, phosphoethanolamine, phosphoserine, and phosphoinositol) attached at the Sn3 position of the PL glycerol backbone. Overall, differences in the proportions of PL classes between immune cell types were generally unremarkable (Fig. 1d). Of note, neutrophils had an increased proportion of PS, a finding that is consistent with the role of PS as a marker of apoptotic cells, and the very high number of neutrophils that must be efficiently cleared via efferocytosis by bone marrow-resident macrophages (Liew and Kubes, 2019). The proportion of PG was markedly higher in T and B lymphocytes compared with NK cells and myeloid cells (Fig. 1d and Extended Data Fig. 1a). PG is a critical substrate in the synthesis of cardiolipins, a major component of the mitochondrial inner membrane, and may thus be expected to be lower in myeloid cells, particularly granulocytes, where mitochondrial abundance is typically low.

Another source of chemical complexity within PLs is the usage of different chemical linkages (ester *vs* ether/vinyl-ether) to attach fatty acyl chains and fatty alcohols to the sn1 position of the PL glycerol backbone. Therefore, we assessed the proportions of diacyl PLs, alkyl-ether PLs [PL(O)], and alkenyl-ether [PL(P)] (also referred to as plasmalogens) within total PC and PE, the major PL classes within which ether PLs are enriched. While the proportion of total PC and PE within the PL pool was similar between cell types (Fig. 1d), a striking difference was observed in PC and PE composition (Fig. 1e,f). Diacyl-PC which accounts for ∼90% of total PC in lymphoid cells, was reduced to ∼25-70% in myeloid cells (Fig. 1e). Similarly, the proportion of diacyl-PE, which accounted for ∼70% of total PE in lymphoid cells, was reduced to ∼25-50% in myeloid cells (Fig. 1f). Accordingly, the proportions of PC(O), PC(P), PE(O), and PE(P) were significantly higher in myeloid cells than lymphoid cells (Fig. 1e,f). In addition to this lymphoid-myeloid distinction, the proportions of diacyl-PC/PE to ether-PC/PE were significantly different between myeloid cells, being lowest in neutrophils and eosinophils, intermediate in basophils, and highest in monocyte subsets (Fig. 1e,f). Accordingly, the ratio of ether PL to diacyl PL was strikingly altered between immune cell types (Extended Data Fig. 1b,c). These differences in diacyl, alkyl-ether and alkenyl-ether composition within PE and PC are a major driver of the variance observed in the global lipidome between immune cell types.

Another major driver of PL diversity arises from the composition of the acyl/alkyl chains, which typically vary from 16 to 24 carbon atoms in length and contain 0 to 6 double bonds. Such differences are functionally important as they influence the biophysical and biochemical properties of the membrane (Harayama and Riezman, 2018). Saturated 16:0 and 18:0 chains constituted a large proportion (∼50%) of the overall PL pool, which was relatively consistent between immune cell types (Fig. 1g), with some exceptions. Specifically, the proportion of 16:0 was decreased and 18:0 increased in neutrophils and eosinophils relative to other cell types; the opposite effect was observed in naïve and memory B cells (Fig. 1g). The proportions of mono-unsaturated chains, primarily 16:1 and 18:1, were significantly altered between cell types. The proportion of 16:1 was lower in myeloid cells compared with lymphoid cells (Fig. 1g; ∼2% *vs* 5%, respectively), while the proportion of 18:1 within the PL pool was markedly higher in granulocytes (neutrophils [39%], eosinophils [22%], and basophils [27%]) compared with all other cell types (10-15% in lymphoid cells and 14-19% in monocytes) (Fig. 1g). With regards to poly-unsaturated species, the proportions of 20:4 and 22:6 within the overall PL pool was reduced in granulocytes relative to the other cell types (Fig. 1g). To provide a more detailed breakdown of PL composition, we calculated alky/acyl chain composition within the major PL classes (PC, PE, PI, and PS) (Extended Data Fig. 2). These analyses highlight the remarkable diversity in alky/acyl composition between PL classes, demonstrating that the differences in global PL alky/acyl chain composition between immune cell types (Fig. 1g) are attributable to specific compositional differences within particular PL classes (Extended Data Fig. 2). For example, the higher overall proportion of 18:1 in granulocytes is largely driven by an increase within PE and PS (Extended Data Fig. 2d,h). Moreover, within granulocytes, most notably neutrophils, the reduced overall proportion of 20:4 is largely due to differences in PC and PE (Extended Data Fig. 2a,d), while the reduced overall proportion of 22:6 is largely due to differences in PE(P) and PS (Extended Data Fig. 2f,g). Indeed, the proportion of 20:4 within PC and PE displays remarkable variance between immune cell types; e.g. comprising ∼12% (PC) and ∼30% (PE) of the total acyl chain content within lymphoid cells, but only ∼1% (PC) and ∼4% (PE) in neutrophils (Extended Data Fig. 3a,b). A similar degree of variance was observed for the proportion of 22:6 within PE(P) and PS (Extended Data Fig. 3c,d). Collectively, the above analysis reveals that differences in PL composition are a defining feature of the human immune cell lipidome. Specifically, we identify differences in the usage of chemical linkages (i.e. ester, ether, and vinyl-ether) and alky/acyl chain diversity within PLs as major drivers of lipidome variance between immune cell types.

### Sphingolipid composition is largely conserved between the cells of human immune system

SLs are major constituents of cell membranes and play critical roles in cell biology (Hannun and Obeid, 2018). Compositional diversity within SLs occurs as a result of differences in the sphingoid base, N-acyl chain, and the status of the headgroup, e.g. SM, glucosyl-ceramides, and ceramide-1-phosphate. We firstly assessed SL diversity based on the absence (ceramide, dihydroceramide, deoxy-ceramide) or presence (hexosyl [Hex]-ceramide [1, 2, or 3 sugar moieties], ganglioside, and sphingomyelin) of different SL head groups. While several significant differences between cell types were observed, very few lineage or cell subclass effects were observed in SL head group diversity, with two primary exceptions: (i) striking increases and decreases in the proportions of Hex2ceramide and SM, respectively, in neutrophils; and (ii) an increased proportion of ceramide and Hex-ceramide and a decrease in the proportion of SM in both naïve and memory B cells (Extended Data Fig. 4a). Next, we assessed variance in the length of the N-acyl chain within the ceramide class of SLs. Very few lineage or cell subclass effects were observed for N-acyl chain diversity. A notable exception occurred within naïve and particularly memory B cells, which had an increased proportion of the shorter 16:0 and 18:0 N-acyl chains compared with all other cell types (Extended Data Fig. 4b). Finally, we examined chemical diversity within the sphingoid base of ceramides. In most SLs, the palmitate (16:0)-derived d18:1 sphingoid base is the most prevalent. However, other fatty acids can form atypical sphingoid bases, i.e. myristate (14:0) and stearate (18:0), which form the corresponding d16:1 and d20:1 sphingoid bases (Han et al., 2009; Hannun and Obeid, 2018). Finally, an atypical d18:2 sphingoid base can be formed by the desaturation of the 18:1 sphingoid base by fatty acid desaturase 3 (FADS3) (Karsai et al., 2020). As expected, d18:1 was the primary sphingoid base in all cell types (Extended Data Fig. 4c). While little difference between cell types was observed, we noted that CD56^bright^ NK cells were enriched in 20:1 sphingoid bases, while eosinophils had elevated 18:2 sphingoid bases (Extended Data Fig. 4c). Collectively, these data demonstrate that while chemical diversity within SL is not a major driver of global lipidome variance between the cells of the immune system, specific SL phenotypes were observable in neutrophils, B cells, and CD56^bright^ NK cells, which could directly impact their function.

### A lipidomic cell atlas of the mouse immune system

To create a lipidomic atlas of the murine immune system, we used FACS to isolate 8 distinct immune cell types from the blood of C57Bl6/J mice (male, 4-6 weeks of age, fed a standard laboratory mouse diet) and determined cellular lipid composition by mass spectrometry (Fig. 2a). Of the 524 measured lipid species, 400 were significantly different between at least 2 cell types. Visualisation of the global lipidomes of the different murine immune cell types using t-SNE revealed a clear distinction between myeloid and lymphoid cells as well as between the myeloid cell types (Fig. 2b). As in the human lipid atlas, T cell subsets and NK cells were indistinguishable from each other, while B cells formed a somewhat distinct cluster (Fig. 2b). Ly6C^hi^ and Ly6C^lo^ monocytes, considered to be somewhat analogous to human classical and non-classical monocytes, also formed distinct clusters (Fig. 2b).

**Figure 2.**
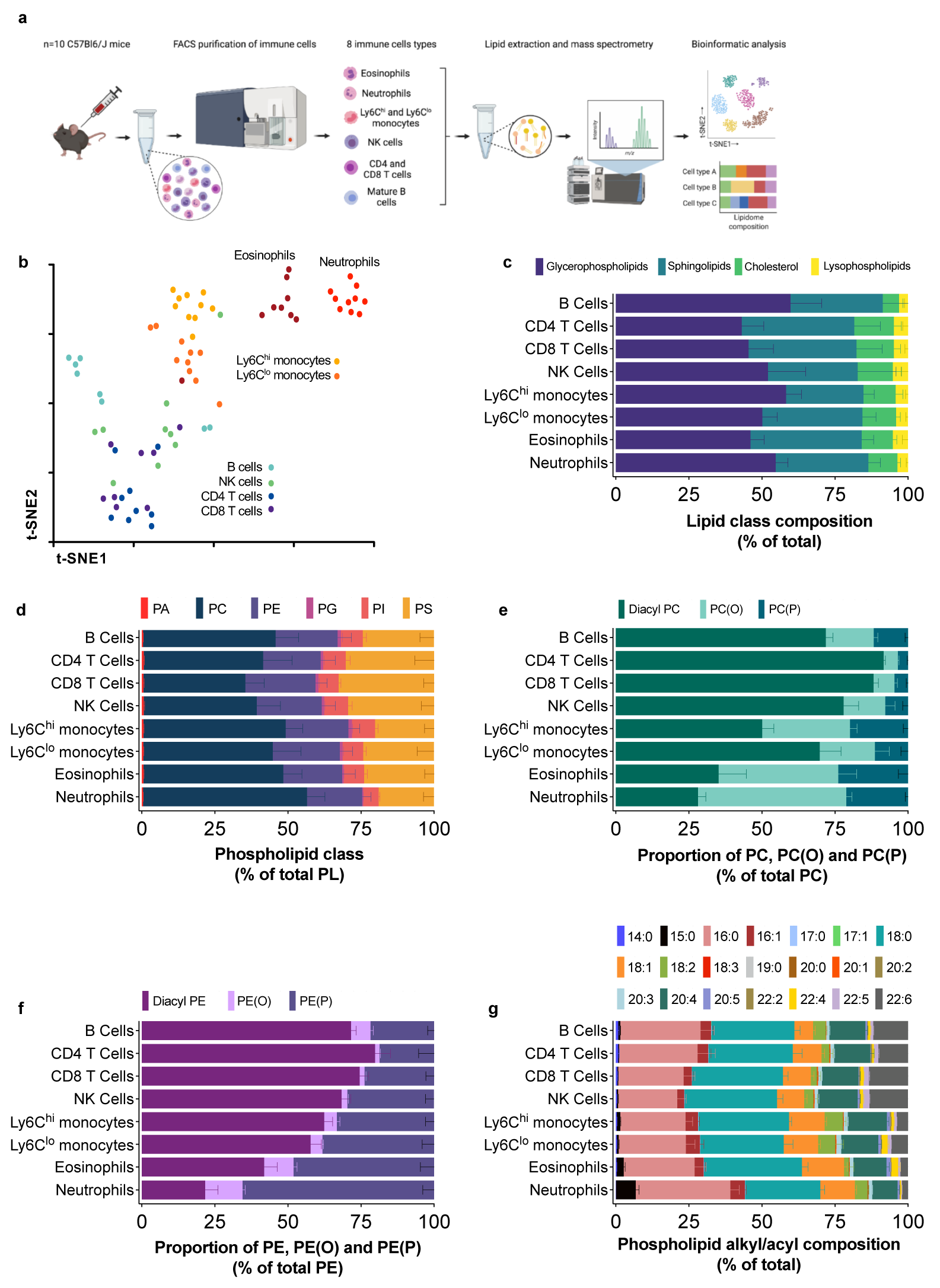
A lipidomic cell atlas of the mouse immune system. **a,** Workflow schematic. **b,** t-SNE map displaying the global cellular lipidome of the mouse immune system. Dots are individual mice; n=8-10 individual mice for each cell type. **c-g,** Breakdown of lipid features in murine immune cells by class (**c**), phospholipid headgroup type (**d**), type of Sn1 chemical linkage in PC (**c**) and PE (**f**), and PL alkyl/acyl composition (**g**). Data are shown as mean + S.D from n=8-10 mice. In each figure, statistically significant differences in specific lipid features between cell types was determined by 1-way ANOVA with Tukey’s HSD test after false discovery rate correction (5%; Benjamini-Hochberg). P values for all pairwise comparisons are shown in Supplementary Excel file 2.

Similar to the human immune cells, there were no marked cell lineage or cell type distinctions in the overall proportions of PL, SL, lyso-PL, and cholesterol or the proportions of the major PL classes (Fig. 2c,d; significance values for all mouse lipid atlas data are shown in Supplementary Excel file 2). However, as in the human data set, the proportion of PG was significantly reduced in neutrophils and eosinophils relative to other cell types (Extended Data Fig. 1d). The proportions of diacyl-PC/PE, ether-PC(O)/PE(O), and vinyl-ether PC(P)/PE(P) within PC and PE classes were markedly altered between lymphoid and myeloid cells as well as between myeloid cells (Fig. 2e,f and Extended Data Fig. 1e,f). These observations parallel those in human immune cells and collectively demonstrate that the utilization of distinct sn1 chemical linkages (ester, ether, vinyl-ether bonds) between cell types (primarily myeloid *vs* lymphoid) is a conserved feature of the human and mouse immune systems.

We next examined the diversity in global PL alky/acyl composition between murine immune cells. While numerous significant differences were present between specific cell types, no clear lineage distinctions were observed (Fig. 2g), with the exception of 22:6, the proportion of which was markedly lower in myeloid relative to lymphoid cells. Analysis of alkyl/acyl composition within PC, PE, PI, and PS classes revealed numerous lineage and cell sub type-specific effects (Extended Data Fig. 2i-p). Firstly, the proportion of 20:4 within PC (Extended Data Fig. 2i, 3e) and PE (Extended Data Fig. 2l, 3f) was markedly reduced in neutrophils and eosinophils, and to a lesser extent in monocytes. The aforementioned decrease in the proportion of 22:6 in myeloid cells was driven by differences within PE(P) (Extended Data Fig. 2n, 3g) and PS (Extended Data Fig. 2p, 3h). Finally, the proportion of 18:1 within PS was significantly higher in neutrophils and eosinophils (Extended Data Fig. 3p). This profile of changes is similar to that observed within the human immune system. Finally, it is noteworthy that while substantial variance is observed in alky/acyl composition between PL classes, human and mouse immune systems show remarkable consistency across the PL classes (Extended Data Fig. 2).

### The cells of the mouse immune system have relatively similar profiles of sphingolipid composition

As in the human immune system, variance in SL chemical diversity between immune cells is limited. Indeed, the only clear lineage distinction was a decrease in the proportion of Hex-ceramide and an increase in the proportion of SM in granulocytic cells relative to all other cell types (Extended Data Fig. 4d). Very few significant differences between cell types were observed in ceramide N-acyl chain composition or sphingoid base composition (Extended Data Fig. 4e,f). Furthermore, two of the most striking SL effects seen in the human immune system – (i) an increased proportion of Hex2-ceramide in neutrophils, and (ii) an increased proportion of the 20:1 sphingoid base in CD56^bright^ NK cells – was not observed in the mouse immune system.

### The high ether lipid content of mature myeloid cells emerges during myelopoiesis

One of the most striking features of mouse and human immune cell lipid composition was the finding that myeloid cells are enriched in ether-PLs. (Fig. 1e,f and 2e,f). To examine the ontogeny of these observations, we assessed how ether lipid content changed during myeloid cell development. To this end, we performed lipidomic analysis on mature myeloid cells (neutrophils, eosinophils, and Ly6C^hi^ monocytes), committed neutrophil precursors (immature neutrophils), uni-potent progenitors (pre-neutrophils and common monocyte progenitors [cMoP]), and bi-potent progenitors (common myeloid progenitors [CMP] and granulocyte-macrophage progenitors [GMP]) and haematapoietic stem and progenitor cells (lineage marker negative, Sca1^+^ cKit^+^ [LSKs]) isolated from bone marrow of 8 to 10-week-old C57BL6/J mice. We show that the proportions of alkyl- and alkenyl-ether PC (Fig. 3a) and PE (Fig. 3b) progressively increase during myeloid cell development. These changes were particularly marked during neutrophil maturation. Another major feature of mature myeloid cells was a reduced content of 20:4 and 22:6 within PC, PE(P), and PS, (Extended Data Fig. 2), relative to lymphoid cells. The proportions of 20:4 within PC and 22:6 within PE(P) decreased during neutrophil development but were unchanged during eosinophil and monocyte development (Fig. 3c,d). This suggests that the reduced proportion of 20:4 and 22:6 in mature myeloid cells relative to lymphoid cells is the result of: (i) a reduction in the proportion of 20:4- and 22:6-containing PLs during neutrophil development; and (ii) an increase in the proportions of 20:4 and 22:6 during lymphoid cell development (e.g. PE(P)-containing 22:6 accounted for 1.7% of the acyl chains within PE(P) in LSKs and 18% in T cells).

**Figure 3.**
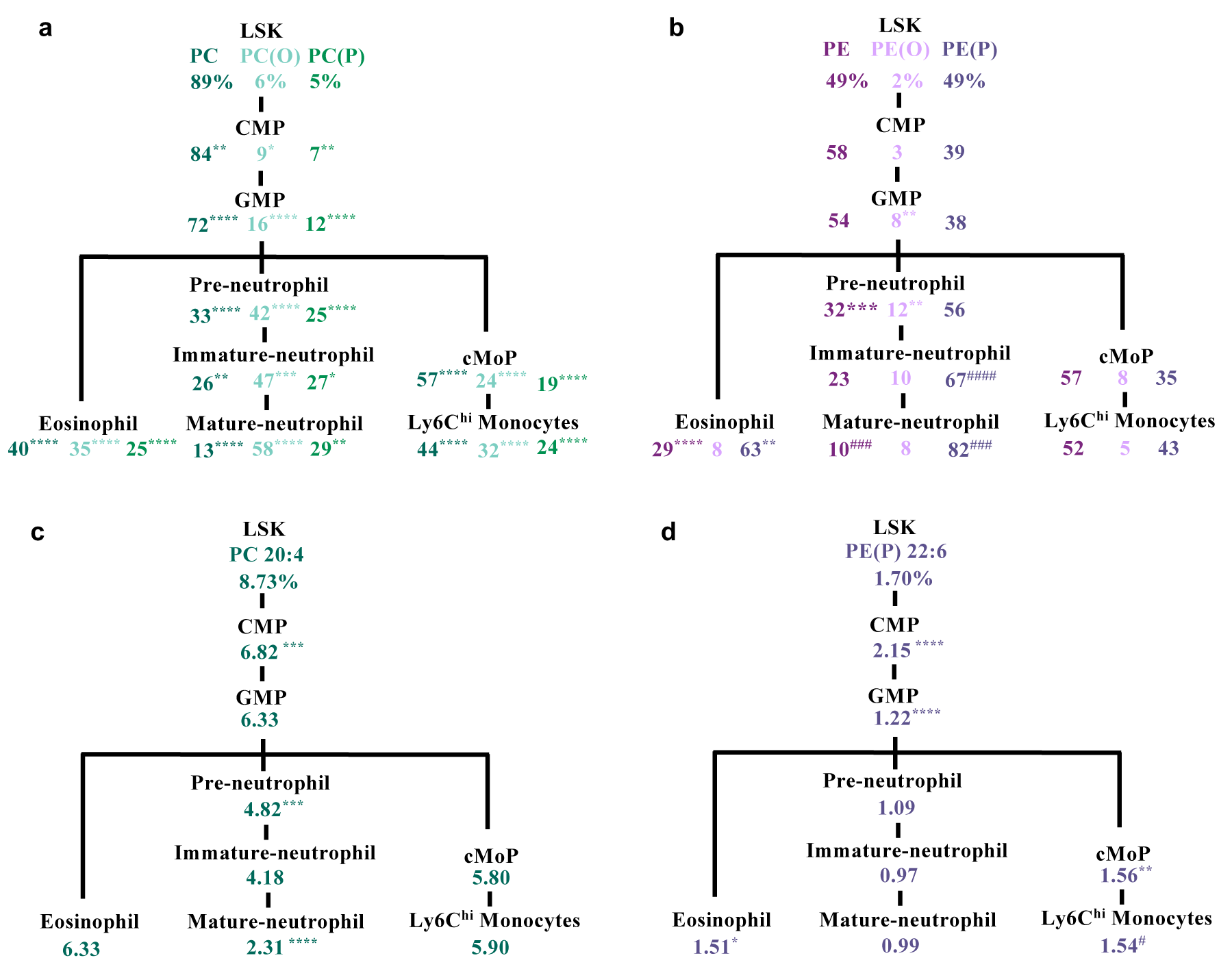
The proportion of diacyl and ether PLs changes during myelopoiesis. **a-d,** Hierarchical representation of the changes in PC composition (**a**), PE composition (**b**), the proportion of 20:4 within PC (**c**), and the proportion of 22:6 within PE(P) during myelopoiesis (**d**). Data shown is the mean of the % of diacyl (PC and PE), ether (PC(O) and PE(O)), and vinyl-ether (PC(P) and PE(P)) within total PC and PE (**a,b**) and the mean of the % of 20:4 and 22:6 within PC and PE(P) (**c,d**). Data are from n=10 individual mice for each cell type. Data was analysed using a 1-way ANOVA with Tukey’s HSD test. *p<0.05, **p<0.01, ***p<0.001, and ****p < 0.0001 between a given cell type and its immediate precursor. ###p<0.001, and ####p < 0.0001 between a given cell type and its immediate precursor but one.

### Lymphoid cells have a higher proportion of highly peroxidation prone PUFA-PLs relative to myeloid cells

One of the most striking features of immune cell lipid composition we identified was marked variance in the proportions of PUFA-containing PLs, particularly those PLs containing highly unsaturated 20:4 and 22:6 PUFAs (Extended Data Fig. 2,3). This observation was particularly interesting as the abundance of PUFA-containing PLs determines the susceptibility to ferroptosis, a form of cell death triggered by iron-dependent lipid peroxidation (Jiang et al., 2021). PE(18:0/20:4) and PE(18:0/22:4) were initially identified as the key executioners of ferroptosis (Kagan et al., 2017). PE(18:0_20:4) displayed considerable variance between immune cells, being highest in T cell subsets and lowest in neutrophils (Extended Data Fig. 5a,c). PE(18:0_22:4) was more variable, although again was lowest in neutrophils (Extended Data Fig. 5b,d). More recent work suggests that overall cellular PUFA abundance dictates susceptibility to ferroptosis (Zou et al., 2020). Therefore, to provide a more holistic view of how differences in PUFA-containing PL abundance may impact immune cell susceptibility to ferroptosis we calculated a cellular peroxidation index (CPI) based on the known relative hydrogen atom transfer propagation rate constants of fatty acids (Xu et al., 2009; Yin et al., 2011) and their proportion within the overall PL pool. In both human and mouse immune cells, lymphoid cells had the highest CPI, followed by monocytes, and granulocytes (Fig. 4a,b). With the exception of PIs, the CPI for the major PL classes was lowest in granulocytes (Extended Data Fig. 5e,f). These observations provide a potential molecular basis for the previously described susceptibility of T cells to ferroptosis, and the resistance of myeloid cells, including neutrophils, monocytes, and macrophages, to ferroptosis (Canli et al., 2017; Drijvers et al., 2021; Jia et al., 2020; Matsushita et al., 2015; Xu et al., 2021a).

### Differences in PUFA-PL composition provide a basis for the differential susceptibility of immune cells to ferroptosis

We hypothesized that immune cells enriched in highly unsaturated PUFA-PLs would be more sensitive to lipid peroxidation and ferroptosis. Accordingly, we examined lipid peroxidation status and ferroptosis susceptibility in murine bone marrow immune cells treated with ML210, a highly-specific inhibitor of glutathione peroxidase (GPX)4, the primary pathway by which cells limit lipid peroxidation to prevent ferroptosis (Jiang et al., 2021). ML210 treatment caused marked lipid peroxidation and cell death in T and B cells, but not Ly6C^hi^ monocytes or neutrophils (Fig. 4c-e). These effects were reversed by co-treatment with radical trapping antioxidants (RTAs) and iron chelation, confirming cells were dying via ferroptosis (Fig. 4f,g and Extended Data Fig. 6a-d). Lipid peroxidation causes PUFA fragmentation, resulting in a decrease in the abundance of PUFA-containing PLs. ML210 treatment decreased the abundance of numerous PUFA-containing PLs in T and B cells (Fig. 4h,i and Extended Data Fig. 7a,b,d,e). This included canonical executioners of ferroptosis PE(18:0_20:4) and PE(18:0_22:4), as well as PUFA-containing ether PLs, PCs, and PSs. These changes were reversed by co-treatment with the ferroptosis inhibitor, Ferrostatin-1 (Fer-1) (Extended Data Fig. 7a,b,d,e). We also observed a decrease in several PUFA-containing PLs in Ly6C^hi^ monocytes (Fig. 4j and Extended Data Fig. 7c,f); however, these changes were not reversed by Fer-1. ML210 treatment did not significantly alter the abundance of any PUFA-containing PLs in neutrophils (Fig. 4k and Extended Data Fig. 7g). To corroborate the ferroptosis and lipid peroxidation results obtained with ML210, we used a second GPX4 inhibitor, RSL3, and erastin, an indirect inhibitor of GPX4 that induces ferroptosis via inhibition of the cystine-glutamate antiporter System X_C-_. Both RSL3 (Extended Data Fig. 8a-g) and erastin (Extended Data Fig. 8h-m) induced cell death and lipid peroxidation in lymphoid cells, but not myeloid cells, which could be reversed by RTAs.

**Figure 4.**
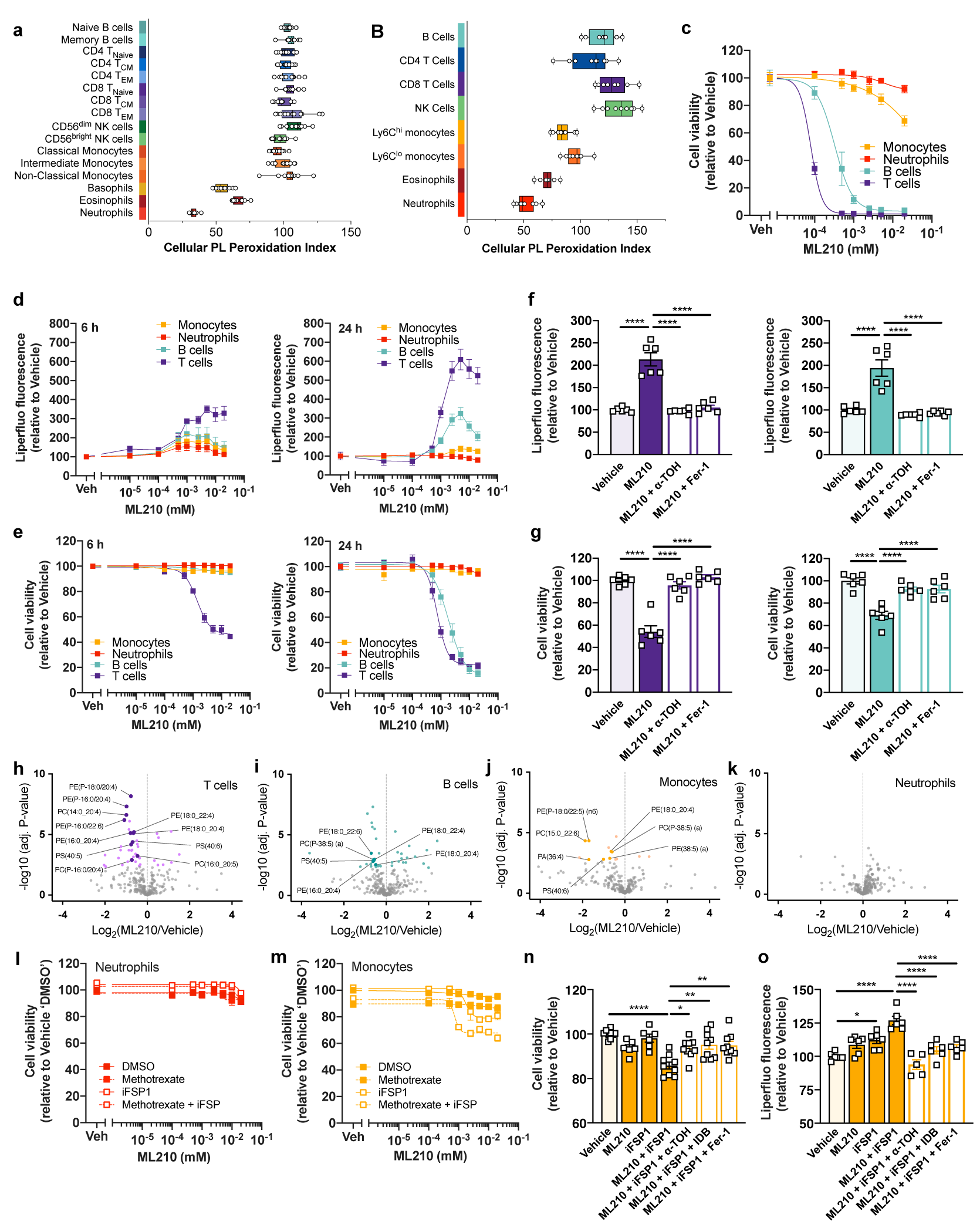
Differences in PUFA-PL composition provide a basis for the differential susceptibility of immune cells to ferroptosis. **a,b,** Cellular phospholipid peroxidation indexes (CPI) of human (**a**; n=11-14 individual donors) and mouse (**b**; n=8-10 individual mice) immune cells. **c**, Cell viability of FACS-sorted neutrophils, monocytes, T and B cells treated with the indicated doses of ML210 for 24 h; n=4 biological replicates. **d,e,** Liperfluo fluorescence (**d**) and cell viability (**e**) in bone marrow neutrophils, monocytes, T, and B cells treated with ML210 at the indicated doses for either 6 h (left panels) or 24 h (right panels); n=6 biological replicates. **f,g,** Liperfluo fluorescence (**f**) and cell viability (**g**) in T and B cells treated with vehicle (DMSO), ML210 (1 µM) or ML210 in combination with either α-tocopherol (α-TOH; 200 µM) or ferrostatin (Fer-1; 1 µM) for 24 h. Data was analysed using a 1-way ANOVA with Tukey’s HSD test. *p<0.05, **p<0.01, ***p<0.001, and ****p < 0.0001. n=6 biological replicates. **h-k,** Volcano plots showing the changes in PL species in FACS-sorted T cells (**h**), B cells (**i**), monocytes (**j**), and neutrophils (**k**) treated with ML210 (1 µM) or vehicle (DMSO) for 24 h. Coloured dots indicate the PL species that were significantly different (1-way ANOVA with Tukey’s HSD test) after false discovery rate correction (Benjamini-Hochberg). n=7-8 biological replicates. **l,m,** Cell viability of bone marrow neutrophils (**l**) and monocytes (**m**) treated with the indicated doses of ML210 alone or in combination with methotrexate (1.5 µM), ferroptosis suppressor protein 1 inhibitor (iFSP1; 3 µM), or methotrexate + iFSP1 for 24 h. n=5-6 biological replicates. **n,o,** Cell viability (**n**) and liperfluo fluorescence (**o**) and in bone marrow monocytes treated with vehicle (DMSO), ML210 (1 µM), iFSP1 (3 µM), or ML210 + iFSP1, in the presence of α-TOH (200 µM), idebenone (10 µM) or Fer-1 (1 µM) for 24 h. n=6-9 biological replicates. Data was analysed using a 1-way ANOVA with Tukey’s HSD test. *p<0.05, **p<0.01, ***p<0.001, and ****p < 0.0001. Data are shown as either a box and whisker plots showing values from individual donors (**a,b**) or mean ± S.E.M (**c-o**).

While it is well established that the abundance of PUFA-containing PLs has a cardinal role in dictating the cellular susceptibility to ferroptosis, cells possess several pathways in addition to GPX4 to prevent excessive lipid peroxidation and ferroptosis. Therefore, the importance of altered lipid composition (i.e. reduced PUFA levels) relative to the potential involvement of other ferroptosis-suppressing pathways in mediating myeloid cell resistance to ferroptosis is unclear. Accordingly, we next determined if neutrophil and Ly6C^hi^ monocyte resistance to lipid peroxidation and ferroptosis following GPX4 inhibition was due to the involvement of the other major ferroptosis suppressing pathways, specifically the ferroptosis-suppressing protein 1 (FSP1), GTP cyclohydrolase 1/dihydrofolate reductase (GCH1/DHFR), and dihydroorotate dehydrogenase (DHODH) pathways (Bersuker et al., 2019; Doll et al., 2019; Mao et al., 2021; Soula et al., 2020). Remarkably, neutrophils remained largely resistant to ferroptosis following co-treatment with the GPX4 inhibitor ML210 and iFSP1 (an FSP1 inhibitor); ML210 and methotrexate (a DHFR inhibitor); ML210 and SPRi3 (an inhibitor of sepiapterin reductase – an enzyme that regulates BH_4_ synthesis); and even triple co-inhibition of GPX4, FSP1, and GCH1/DHFR (Fig. 4l and Extended Data Fig. 6e). Combined inhibition of GPX4 and FSP1 partially sensitised Ly6C^hi^ monocytes to ferroptosis and lipid peroxidation, effects that were reversed by co-treatment with RTAs (Fig. 4m-o). Ferroptosis in T and B cells was not exacerbated following combined inhibition of the GPX4, FSP1, and GCH1/DHFR pathways (Extended Data Fig. 6f,g). To determine if the DHODH pathway was responsible for resistance to lipid peroxidation and ferroptosis, we treated cells with brequinar, a DHODH inhibitor previously shown to induce ferroptosis (Mao et al., 2021). Brequinar treatment alone was sufficient to cause substantial cell death in neutrophils and Ly6C^hi^ monocytes (Extended Data Fig. 9a,b), which was associated with an increase in lipid peroxidation (Extended Data Fig. 9c,d). However, brequinar-induced cell death could not be reversed by treatment with α-TOH, Fer-1, or the iron chelator ciclopirox, demonstrating that neutrophil and Ly6C^hi^ monocyte cell death is unlikely to be ferroptosis (Extended Data Fig. 9e,f). Notably, an inhibitor of necroptosis (Necrostatin-1s; Nec-1s), but not apoptosis (Z-VAD-FMK), reversed brequinar-induced cell death, but not lipid peroxidation, in neutrophils (Extended Data Fig. 9e) and partially prevented brequinar-induced cell death in Ly6C^hi^ monocytes (Extended Data Fig. 9f). Collectively, these data suggest that differences in the abundance of PUFA-containing PLs is a major determinant of immune cell susceptibility to lipid peroxidation and ferroptosis.

### Human neutrophils and T cells are differentially susceptible to ferroptosis

We next examined if the differential susceptibility of murine lymphoid and myeloid cells to ferroptosis was observed in human immune cells. Accordingly, purified T cells and neutrophils isolated from peripheral blood samples of healthy volunteers were treated with ML210. T cells displayed a loss of viability and increased lipid peroxidation following ML210 treatment, which was reversed by Fer-1 (Fig. 5a,b). In contrast, human neutrophils displayed very little change in viability and lipid peroxidation in response to ML210 (Fig. 5c,d). These findings demonstrate that the differential susceptibility of lymphoid and myeloid cells to lipid peroxidation and ferroptosis is a conserved feature of the mouse and human immune systems.

**Figure 5.**
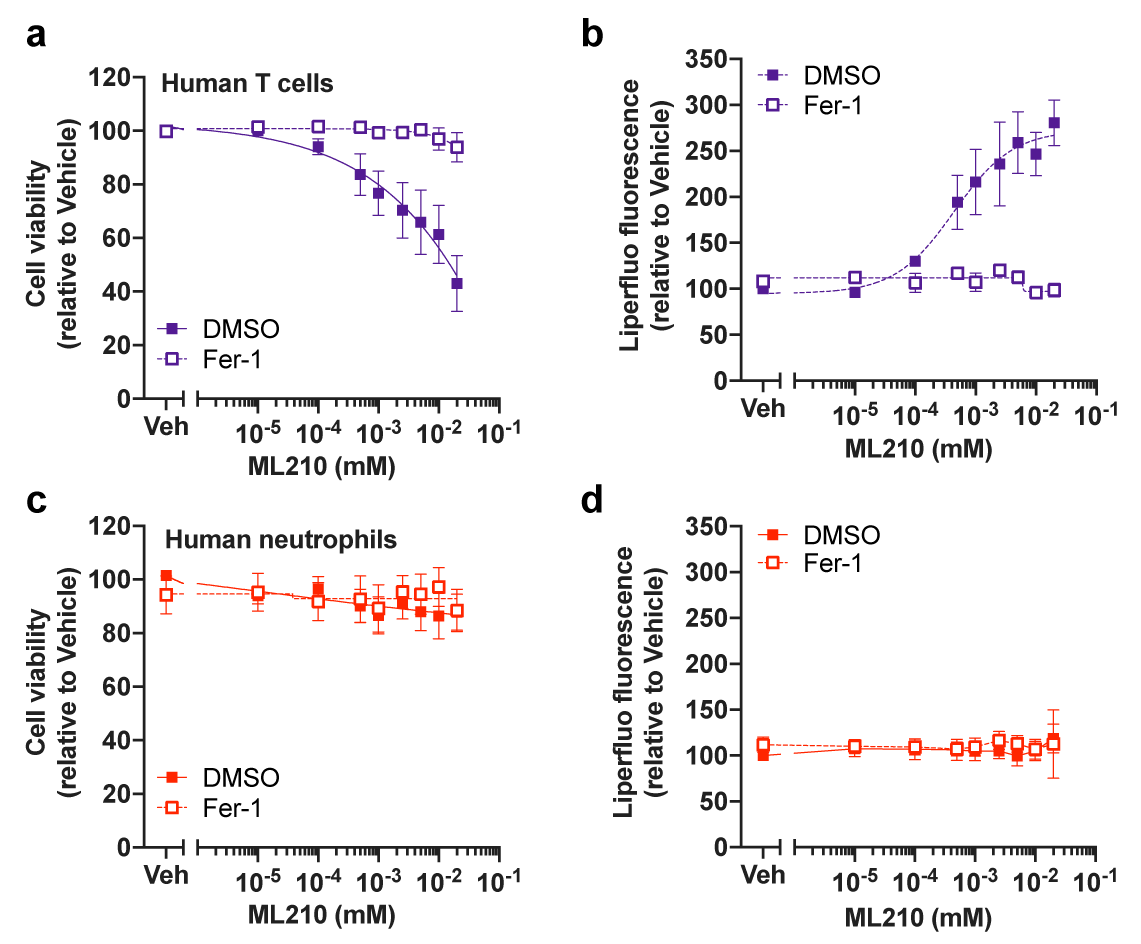
Human neutrophils and T cells are differentially susceptible to ferroptosis. **a-d,** Cell viability and liperfluo fluorescence in T cells (**a,b**) and neutrophils (**c,d**) treated with ML210 at the indicated doses in the presence or absence of Fer-1 (1 µM). Data are from n=5-6 individual donors. All data are shown as mean ± S.E.M.

### Altering cellular lipid composition changes immune cell susceptibility to lipid peroxidation and ferroptosis

Our data suggests that the PUFA content of immune cells governs their susceptibility to ferroptosis. Accordingly, we hypothesized that altering endogenous PUFA levels would change ferroptosis susceptibility. Firstly, we supplemented T cells with either PE(18:0/18:1) or the MUFA oleate (18:1) to reduce the proportion of PUFA-PLs in the membrane. Both oleate and PE(18:0/18:1) shifted the T cell PL composition towards a MUFA enriched profile, concomitantly decreasing the proportion of PLs containing 4 carbon-carbon double bonds, e.g. 20:4-containing PLs (Extended Data Fig. 10a-c). Importantly, both oleate and PE(18:0/18:1) supplementation protected T cells from lipid peroxidation and ferroptosis triggered by GPX4 inhibition (Fig. 6a,b). Next, we supplemented neutrophils with arachidonic acid (20:4) and docosahexaenoic acid (22:6) (AA + DHA) to increase endogenous PUFA levels. Indeed, AA + DHA treatment reduced the proportions of 1 and 2 carbon-carbon double bond-containing PLs, while concomitantly increasing the proportions 5, 8 and 10 carbon-carbon double bond-containing PLs, e.g. 20:4 and 22:6-containing-PLs (Extended Data Fig. 10d,e). Remarkably, in the absence of treatment with ML210, AA + DHA supplementation caused a dose-dependent cell death and significantly increased lipid peroxidation in neutrophils (Fig. 6c,d). These effects were largely reversed by co-treatment of neutrophils with Fer-1 but not inhibitors of apoptosis or necroptosis, indicating *bona fide* ferroptosis (Fig. 6e,f). Collectively, these data demonstrate that differences in PUFA abundance contribute to the differential susceptibility of immune cells to lipid peroxidation and ferroptosis.

**Figure 6.**
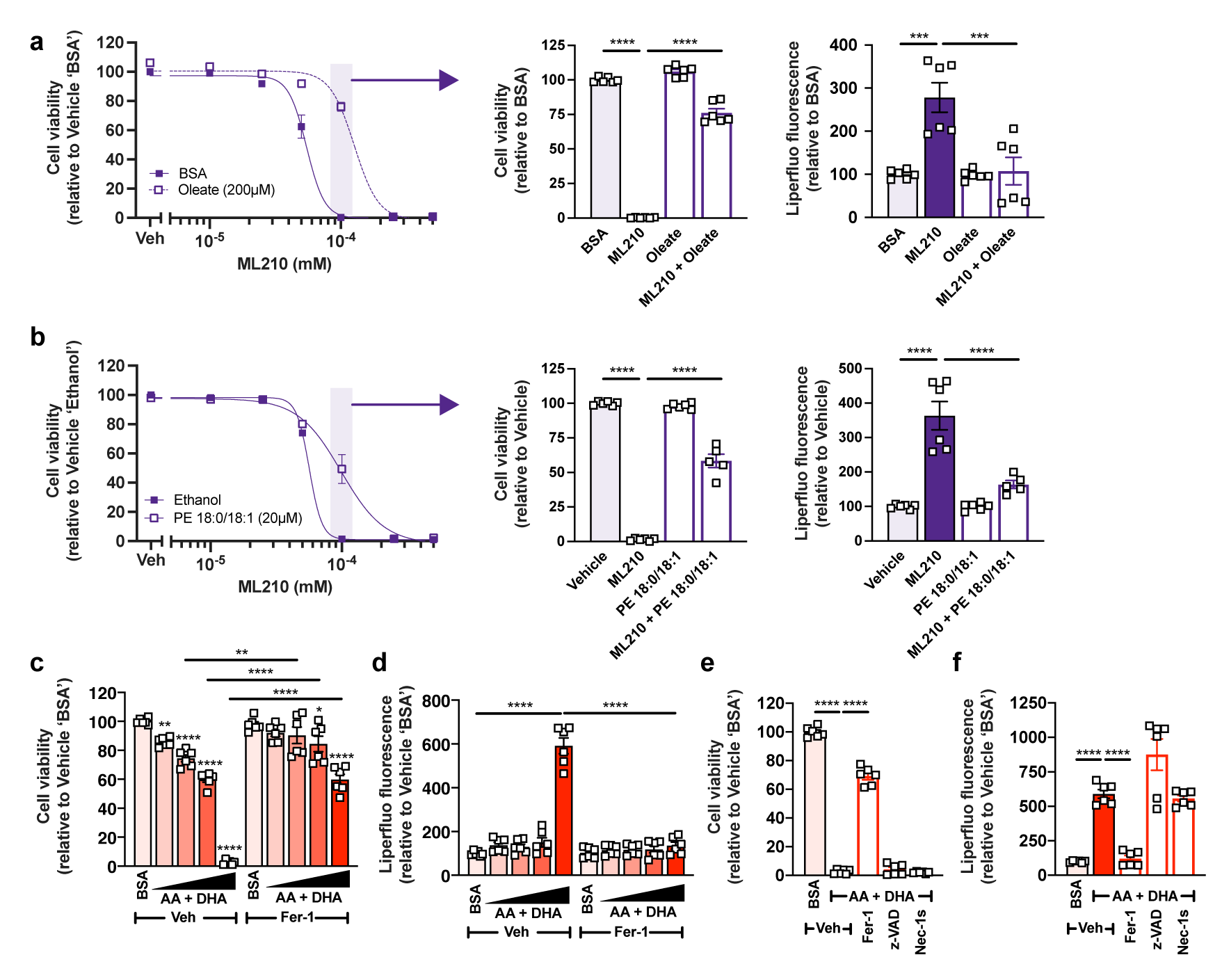
Lipid supplementation in T cells and neutrophils alters sensitivity to lipid peroxidation and ferroptosis. **a**, Cell viability and liperfluo fluorescence purified T cells treated with the indicated doses of ML210 for 24 h following pre-treatment with oleate (200 µM) or vehicle (bovine serum albumin; BSA) for 16 h. n=6 biological replicates. **b,** Cell viability and liperfluo fluorescence in purified T cells treated with the indicated doses of ML210 for 24 h following pre-treatment with PE 18:0/18:1 (20 µM) or vehicle (ethanol) for 16 h. n=5-6 biological replicates. **c,d,** Cell viability and liperfluo fluorescence in purified neutrophils treated with an AA + DHA mixture (0 [BSA], 50, 100, 200, or 400 µM) in the presence of Fer-1 (1 µM) or vehicle (DMSO) for 24 h. n=6 biological replicates. **e,f,** Cell viability and liperfluo fluorescence in purified neutrophils treated with a AA + DHA mixture (400 µM) in the presence of Fer-1 (1 µM) or vehicle (DMSO) for 24 h or following pre-treatment with z-VAD (25 µM) or Nec-1s (10 µM) for 1 hour. n=6 biological replicates. Data was analysed using a 1-way or 2-way ANOVA with Tukey’s HSD test. *p<0.05, **p<0.01, ***p<0.001, and ****p < 0.0001. Data are shown as mean ± S.E.M.

### NADPH oxidase 2 (NOX2)-derived ROS promote lipid peroxidation and ferroptosis in neutrophils

The NADPH oxidase family of enzymes (NOX1-5, DUOX1,2) produce ROS and have previously been implicated in the regulation of ferroptosis in cancer cells (Dixon et al., 2012). NOX2 is the NOX isoform expressed in phagocytic cells and is highly expressed in neutrophils. Accordingly, we hypothesised that NOX2 is the source of the ROS that trigger lipid peroxidation and ferroptosis in neutrophils supplemented with AA + DHA. Indeed, neutrophils from NOX2 KO mice were protected from AA + DHA-induced lipid peroxidation and ferroptosis (Fig. 7a,b). Following activation, NOX2 rapidly produces large quantities of superoxide which plays a critical role in neutrophil-mediated immunity. A notable feature of NOX2 activation (and other NOXs) is that in addition to producing large amounts of ROS, they also consume large amounts of NADPH. NADPH is a crucial co-factor in regenerating reduced GSH, which is required for GPX4 function. Thus, we hypothesized that the activation of NOX2 would create conditions within the cell that promote lipid peroxidation and ferroptosis. Neutrophils treated with phorbol 12-myristate 13-acetate (PMA), a potent NOX2 activator, had increased lipid peroxidation and a moderate reduction (p=0.09) in cell viability (Fig. 7c,d). Notably, inhibition of GPX4 in PMA stimulated neutrophils caused a marked loss of cell viability and further increased lipid peroxidation (Fig. 7c,d). Importantly, these effects were reversed by Fer-1 and NOX2 deletion, confirming that the cell death observed was ferroptosis and the dependence on NOX2 (Fig. 7a,b). Thus, while non-activated neutrophils do not use GPX4 to prevent excessive lipid peroxidation and ferroptosis, activated neutrophils rely on GPX4 to minimise self-inflicted lipid damage caused by heightened ROS production. Finally, we hypothesised that increasing the PUFA-PL content of neutrophils would exacerbate PMA-induced ferroptosis. In this experiment, we supplemented neutrophils with a dose (50µM) of AA + DHA that caused minimal cell death (Fig. 6c-h) but still effectively increased PUFA-PL content (Extended Data Fig. 10d,e) for 16 h before stimulating with PMA. Consistent with the previous experiment, in vehicle treated neutrophils PMA alone induced modest cell death (p=0.12), which was enhanced by concomitant GPX4 inhibition and reversed by Fer-1 (Fig. 7e). Notably, PMA treatment induced a more pronounced loss of cell viability in neutrophils supplemented with AA + DHA (Fig. 7e). Interestingly, ML210 treatment did not further reduce cell viability (Fig. 7e). Collectively, our data demonstrate that a reduced PUFA-PL content is essential for both resting and activated neutrophils to minimise lipid peroxidation and ferroptosis driven by NOX2-dependent ROS production.

**Figure 7.**
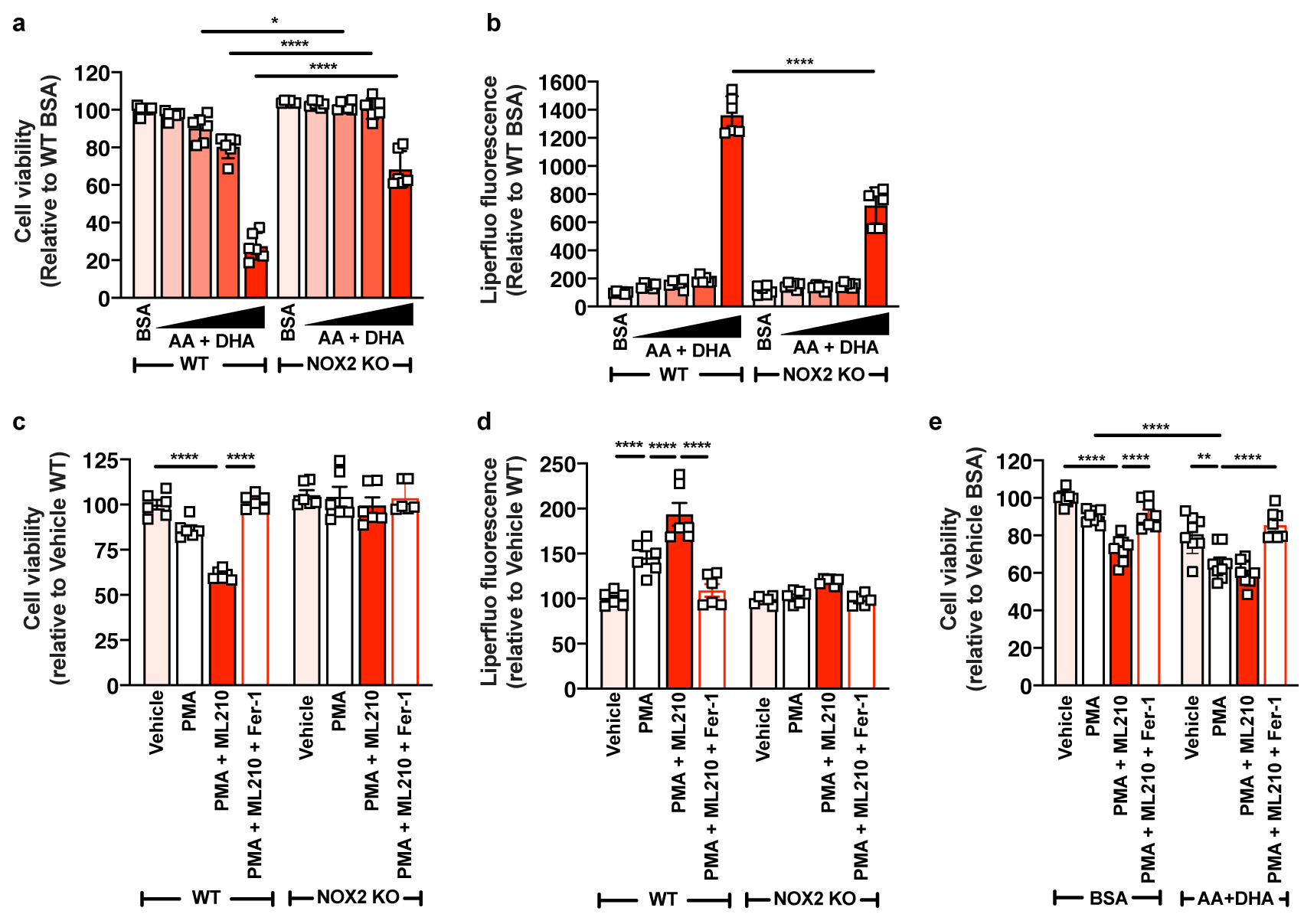
NADPH oxidase 2 (NOX2)-derived ROS promote lipid peroxidation and ferroptosis in neutrophils. **a,b,** Cell viability and liperfluo fluorescence in purified neutrophils from WT and NOX2 KO mice treated with an AA + DHA mixture (0 [BSA], 50, 100, 200, or 400 µM) for 24 h. n=6 biological replicates. **c,d,** Cell viability and liperfluo fluorescence in purified neutrophils from WT and NOX2 KO mice treated as indicated with PMA (50 nM), ML210 (5 µM), and Fer-1 (1 µM) for 6 h. n=6 biological replicates. **e,** Cell viability in purified neutrophils that were supplemented with either AA + DHA (50 µM) or vehicle (bovine serum albumin; BSA) for 16 h before being treated as indicated with PMA (50 nM), ML210 (5 µM), and Fer-1 (1 µM) for 6 h. n=8 biological replicates. **a-e,** Data was analysed using a 2-way ANOVA with Tukey’s HSD test. *p<0.05, **p<0.01, ***p<0.001, and ****p < 0.0001. Data are shown as mean ± S.E.M.

## Discussion

Many features define immune cell identity, from physical parameters such as cell size, nuclear morphology, and granule content, to molecular features such as the transcriptome and protein surface marker expression. Importantly, such differences underlie the unique functional properties of distinct immune cell types. The major motivation for the current work was the idea that the lipid landscape of the cells of the immune system is also a defining feature of immune cell identity and that differences in lipid composition contribute to cell-specific functionality. To address this idea, we created a cellular lipid atlas of the immune system. This resource, which comprises >500 individual lipid species measured in immune cell types from different developmental lineages and with distinct roles in host immunity, in both mice and humans, demonstrates that the lipid composition of immune cells is distinct. By analysing the major chemical features that define lipid identity and function (e.g. PL headgroup, acyl chain diversity, chemical linkages), we have identified unique, immune cell-specific lipid profiles and provided evidence that these differences have important biological consequences.

One of the most striking differences in lipid composition we identified from the lipid atlas was the ∼ 2-3-fold enrichment of ether lipids in myeloid cells, in particular granulocytes, compared with lymphoid cells. Furthermore, the high level of ether lipids in mature myeloid cells is the result of lipidome re-modelling during myelopoiesis. These changes are particularly pronounced during neutrophil development, from ∼90% diacyl PC to ∼90% ether/vinyl-ether PC and from ∼50% diacyl PE to ∼90% ether/vinyl-ether PE in LSKs and mature neutrophils, respectively. While an increase in the proportion of ether lipids in myeloid cells, particularly neutrophils, has been appreciated for several decades (Mueller et al., 1982; Ramesha and Pickett, 1987), their functional importance is not well understood. Ether PLs are distinguished from conventional diacyl PLs by the presence of an ether or vinyl-ether bond used to attach the sn1 alkyl chain, as opposed to an ester linkage in diacyl PLs (Dean and Lodhi, 2018; Paul et al., 2019). This relatively modest structural difference imparts ether PLs with unique biophysical and biochemical roles, including altering membrane properties, in cell signalling, and as anti-oxidants (Dean and Lodhi, 2018; Paul et al., 2019). Previous studies have shown that activated neutrophils and eosinophils produce chlorinating and brominating reactive species that attack the vinyl-ether bond to produce 2-chlorohexadecal and 2-bromohexadecanal, respectively, that can act as phagocyte chemoattractants (Albert et al., 2002; Albert et al., 2003; Thukkani et al., 2002). In neutrophils, ether lipids may influence membrane lipid composition to limit ER stress-induced apoptosis and neutropenia (Lodhi et al., 2015); however, it has also been reported that mice deficient in ether lipids have normal neutrophil numbers (Dorninger et al., 2015). Our data support the idea that ether lipids have critical roles in the development and function of neutrophils, and likely within other granulocytes and myeloid cells, the elucidation of which will be an important future goal.

Having determined that different immune cell types have specific lipid signatures, we wanted to establish that such differences confer cell-specific functional properties. For several reasons, we focused on PUFA-containing PL. Firstly, in both mice and humans, the proportions of 20:4 and 22:6 within PC, PE, and PS were markedly lower in mature myeloid cells, most notably in neutrophils. Due to the presence of multiple abstractable H atoms, PUFAs are highly susceptible to damage by oxygen containing radicals (Yin et al., 2011). This feature of PUFAs is central to their involvement in ferroptosis (Jiang et al., 2021). While ferroptosis has primarily been studied in cancer cells, amongst non-cancer cells, the role of ferroptosis in T cells is quite well-described. While ferroptosis does not appear to contribute to T cell development, T cells are highly susceptible to ferroptosis triggered by GPX4 deletion, leading to reduced numbers of T cells in the periphery and a failure to expand in number in response to infection (Matsushita et al., 2015). Moreover, even when GPX4 is present, T cells appear susceptible to lipid peroxidation and ferroptosis, as has been described for tumour infiltrating CD8^+^ T cells (Ma et al., 2021; Xu et al., 2021b) and follicular helper T cells (Yao et al., 2021), the consequences of which are impaired anti-tumour immunity and impaired responses to immunization, respectively. Why T cells are susceptible to lipid peroxidation, yet other immune cells, in particular myeloid cells, are resistant, was unclear. We provide a parsimonious explanation for this phenomenon, demonstrating that T cells are enriched in PUFA-containing PLs, notably those containing 22:6, which are highly susceptible to lipid peroxidation. Importantly, we show that biasing the composition of the T cell lipidome towards a more MUFA-enriched profile reduced the susceptibility of T cells to lipid peroxidation and ferroptosis. Consistent with these findings, deletion of acetyl-CoA synthetase long chain family member 4 (ACSL4), the enzyme primarily responsible for promoting the incorporation of free PUFAs into PUFA-containing PLs, protects T cells from ferroptosis (Drijvers et al., 2021). In view of their propensity to undergo peroxidation, the high content of 22:6-containing PLs in T cells implies an essential role of these PLs in T cell function. Indeed, while ACSL4 deficient T cells are resistant to ferroptosis, they have impaired anti-tumour CD8^+^ T cell immune responses (Drijvers et al., 2021). The role of PUFA-containing PLs, particularly 22:6, in T cell function will be an important area for future work.

Previous studies have shown that myeloid cells, including neutrophils and monocytes, are resistant to ferroptosis triggered by GPX4 deletion (Canli et al., 2017). Using our lipid atlas, we identified that neutrophils, and to a lesser extent Ly6C^hi^ monocytes, had markedly reduced levels of PUFA-containing PLs, particularly those containing the highly peroxidizable 20:4 and 22:6. Given the cardinal role of PUFA-containing PLs in ferroptosis, these observations provide an explanation for the resistance of myeloid cells to lipid peroxidation and ferroptosis. Indeed, using multiple approaches, we show that the neutrophil lipidome is resistant to lipid peroxidation triggered by GPX4 inhibition. GPX4’s unique ability to reduce PL-hydroperoxides to PL-alcohols place it at the centre of the cell’s defence against lipid peroxidation and ferroptosis (Jiang et al., 2021; Yang et al., 2014). However, several additional pathways, in particular the DHODH (Mao et al., 2021), FSP1 (Bersuker et al., 2019; Doll et al., 2019), and DHFR (Soula et al., 2020) pathways, help to limit ferroptosis. While we observed a modest protective role of the FSP1 pathway in Ly6C^hi^ monocytes, these pathways do not appear to contribute to the resistance of neutrophils to PL peroxidation and ferroptosis. It is noteworthy that compared with neutrophils, Ly6C^hi^ monocytes have higher levels of PUFA-containing PL yet are similarly resistant to ferroptosis. Indeed, a similar observation can be made about B cells, which have a very similar PUFA-containing PL profile to T cells, yet B cells are more resistant to ferroptosis than T cells. These differences may be explained by differences in the profile of PLs that undergo peroxidation in T and B cells; for example, we observed a marked decrease in ether PL only in T cells, and the peroxidation of ether lipids was recently shown to promote ferroptosis (Zou et al., 2020); or potentially to differences in the levels of known regulators of ferroptosis, e.g. glutathione, iron, and the abundance/activity of GPX4, FSP1. Alternatively, yet to be defined, protective mechanisms may exist to limit ferroptosis in B cells and Ly6C^hi^ monocytes.

It was shown previously that neutrophils are resistant to ferroptosis triggered by GPX4 deletion (Canli et al., 2017). Our results support and extend these observations, demonstrating that even under conditions in which several major lipid peroxidation and ferroptosis suppressing pathway are inhibited, neutrophils are almost completely resistant to lipid peroxidation and ferroptotic cell death. Importantly, we also demonstrate that increasing the PUFA content of neutrophils is sufficient to trigger ferroptosis. These findings suggest that biasing their PL composition towards a less PUFA-enriched profile, thereby minimising peroxidative damage to the cell membrane, is a major mechanism by which neutrophils avoid ferroptotic cell death. Indeed, as described above, a major feature of neutrophil development and maturation is a reduction in the proportion of highly peroxidizable 20:4 and 22:6 PUFAs.

The resistance of neutrophils to lipid peroxidation and ferroptosis relative to other immune cell types strongly suggests that avoiding ferroptotic cell death is an important aspect of neutrophil biology. This raises the question – why are neutrophils, and indeed other myeloid cell types, so resistant to ferroptotic cell death? As a consequence of their roles in host physiology and pathology, neutrophils operate within highly oxidative environments, for example sites of infection and tissue damage. Additionally, a critical homeostatic function of neutrophils and monocytes is to patrol the blood vasculature (Burn et al., 2021), an environment that is rich in oxygen and iron and has recently been demonstrated to promote ferroptosis (Ubellacker et al., 2020). Finally, the production of superoxide (O^-^_2_) by NADPH-oxidase is a critical feature of neutrophil-mediated immunity (Nguyen et al., 2017). O^-^_2_ is rapidly converted to H_2_O_2_, which, in the presence of iron, is converted to HO^•^, a major inducer of lipid peroxidation and ferroptosis. In support of this concept we have demonstrated that neutrophils with an increased content of highly unsaturated PUFA-PLs are more susceptible to ferroptosis induced by NOX2 activation. Accordingly, we propose that the neutrophil lipidome, and that of other myeloid cells, is specifically adapted to guard against oxidative damage to membrane lipids and ferroptotic cell death. These adaptations would allow myeloid cells to survive and function effectively in the toxic environments in which they must operate.

In summary, we have developed a lipid atlas of the cells of the human and mouse immune system. Using this this resource, we have shown that cellular lipid composition is an identifying feature of immune cells. Our analysis of the lipid features of the human and mouse immune system has identified numerous, cell-specific lipid phenotypes that should provide fruitful avenues for further research into how differences in lipid composition impact cell-specific functionality. In this regard, we have used the lipid atlas to identify differences in the abundance of PUFA-containing PL as a major mechanism by which immune cells are differentially susceptible to lipid peroxidation and ferroptosis.

## Supporting information

Human statistics

Mouse statistics

## Acknowledgements

We thank the members of the AMREP Flow Cytometry Core Facility for their expert assistance. We thank the members of AMREP Animal Services who provided wonderful care of the mice used in this work. We thank all the human volunteers who provided blood samples to assist with this work. We are grateful to Professor Chris Sobey for providing access to the NOX2 KO mice. We are extremely grateful to the following funding sources: National Health and Medical Research Council of Australia grants GNT1189012 to G.I.L, GNT1194329 to A.J.M, GNT1197190 to K.H., and GNT2009965 to P.J.M., a CSL Centenary Fellowship to A.J.M., and the Victorian Government’s Operational Support Program.

## Methods

### Preparation of cells for the human immune cell lipid atlas

Human immune cells were obtained from buffy coats received from Red Cross Australia (Melbourne, Australia). Buffy coats were diluted 1:5 with PBS containing 5% FBS and 0.5 mM EDTA. Blood was layered onto a discontinuous Histopaque (Sigma Aldrich, NSW, Australia) density gradient with the densities 1.077 g/ml and 1.119 g/ml to isolate PBMCs and granulocytes, respectively, and centrifuged for 30 mins at 300 g with the brakes off. The two fractions were transferred into separate tubes, washed, and centrifuged for 10 mins at 200 g with the brakes on to remove platelets. The supernatant was removed and pellets were lysed with 1x red blood cell (RBC) lysis buffer (Thermo Fisher) for 5 mins, after which lysis was stopped by the addition PBS containing FBS and EDTA. Samples were then centrifuged at 300 g for 5 mins and the white cell pellet obtained.

### Preparation of cells for the mouse immune cell lipid atlas

Mouse immune cells were obtained from the peripheral blood of 4-6-week-old, male C57Bl/6J mice fed a standard laboratory mouse diet (Irradiated rat and mouse diet; Speciality Feeds, Glen Forrest, Australia). C57Bl/6J mice were housed at the ARA Animal Services Facility. with all procedures approved by the institutional animal ethics committee (ARA AEC). Mice were sacrificed via CO_2_ asphyxiation and blood obtained via cardiac puncture. Blood samples were lysed for 15 mins in RBC, after which lysis was stopped with the addition of 1X FACS buffer (HBSS w/o Ca^2+^ and Mg^2+^ containing BSA and 0.5mM EDTA). Samples were then centrifuged at 3000 rpm for 5 mins at 4°C and the white cell pellet obtained.

### Preparation of cells for the myelopoiesis lipid atlas

Mature myeloid cells and their progenitors were obtained from the bone marrow of 8-10-week-old, male C57Bl/6J mice. To obtain bone marrow cells, hind legs were removed and bones were flushed with PBS without Ca^2+^ and Mg^2+^. Aspirates were centrifuged at 3000 rpm for 5 mins at 4°C, cells treated with 1x RBC lysis buffer for 5 minutes with the reaction stopped with an excess of FACS buffer, and bone marrow cell pellets obtained after a final spin (300 g, 5 minutes).

### Antibody staining and fluorescence-activated cell sorting (FACS)

Cells were stained with the antibody cocktails outlined in the tables below staining for cell-specific surface markers and incubated for 30 minutes on ice. Antibodies were used at a 1:400 dilution unless stated otherwise. Staining was stopped with FACS buffer and cells were subsequently washed and filtered through a 35 µm strainer prior to sorting. The antibodies and sorting panels used to purify the specific human immune cell populations were as follows: Panel 1 – Naïve and Memory B cells; Panel 2 – CD4 and CD8 T cells; Panel 3 – NK cells; Panel 4 – Monocytes; Panel 5 – Basophils; Panel 6 –Neutrophils and Eosinophils The antibodies and sorting panels used to purify the specific murine cell populations were as follows: Panel 1 – B cells, NK cells, CD4 and CD8 T cells; Panel 2 – Ly6C^lo^ and Ly6C^hi^ Monocytes, Neutrophils, and Eosinophils.

| **Antibodies and sorting panels used to purify human immune cells** |  |
| --- | --- |
| **Cell type** | **Antibodies used in sorting panels** |
| Naïve B cell | L/D CD19<sup>+</sup> CD27<sup>+</sup> Mitotracker<sup>-</sup> |
| Memory B cell | L/D CD19<sup>+</sup> CD27<sup>+</sup> Mitotracker<sup>+</sup> |
| CD4 Naïve T cell | L/D CD3<sup>+</sup> CD4<sup>+</sup> CCR7<sup>+</sup> CD45RA<sup>+</sup> |
| CD4 Central Memory T cell | L/D CD3<sup>+</sup> CD4<sup>+</sup> CCR7<sup>+</sup> CD45RA<sup>-</sup> |
| CD4 Effector Memory T cell | L/D CD3<sup>+</sup> CD4<sup>+</sup> CCR7<sup>-</sup> CD45RA<sup>-</sup> |
| CD8 Naïve T cell | L/D CD3<sup>+</sup> CD8<sup>+</sup> CCR7<sup>+</sup> CD45RA<sup>+</sup> |
| CD8 Central Memory T cell | L/D CD3<sup>+</sup> CD8<sup>+</sup> CCR7<sup>+</sup> CD45RA<sup>-</sup> |
| CD8 Effector Memory T cell | L/D CD3<sup>+</sup> CD8<sup>+</sup> CCR7<sup>-</sup> CD45RA<sup>-</sup> |
| CD56 Dim NK cell | L/D CD3<sup>-</sup> CD56<sup>+</sup> CD16<sup>-</sup> |
| CD56 Bright NK cell | L/D CD3<sup>-</sup> CD56<sup>+</sup> CD16<sup>+</sup> |
| Classical Monocyte | L/D Lin<sup>-</sup> HLA-DR<sup>+</sup> CD14<sup>+</sup> CD16<sup>-</sup> |
| Intermediate Monocyte | L/D Lin<sup>-</sup> HLA-DR<sup>+</sup> CD14<sup>+</sup> CD16<sup>+</sup> |
| Non-classical Monocyte | L/D Lin<sup>-</sup> HLA-DR<sup>+</sup> CD14<sup>dim</sup> CD16<sup>+</sup> |
| Basophil | L/D HLA-DR<sup>-</sup> CD203c<sup>+</sup> CD123<sup>+</sup> |
| Eosinophil | L/D SSC<sup>high</sup> Siglec8<sup>+</sup> |
| Neutrophil | L/D SSC<sup>high</sup> CD15<sup>+</sup> CD16<sup>+</sup> |

| **Antibodies and sorting panels used to purify murine immune cells** |  |
| --- | --- |
| **Cell type** | **Antibodies used in sorting panels** |
| B cells | L/D CD45<sup>+</sup> CD3<sup>-</sup> CD19<sup>+</sup> |
| NK cells | L/D CD45<sup>+</sup> CD3<sup>-</sup> NK1.1<sup>+</sup> |
| CD4 T cells | L/D CD45<sup>+</sup> CD3<sup>+</sup> CD4<sup>+</sup> |
| CD8 T cells | L/D CD45<sup>+</sup> CD3<sup>+</sup> CD8<sup>+</sup> |
| Ly6C<sup>lo</sup> monocytes | L/D CD45<sup>+</sup> CD115<sup>+</sup> Gr1<sup>-</sup> |
| Ly6C<sup>hi</sup> monocytes | L/D CD45<sup>+</sup> CD115<sup>+</sup> Gr1<sup>+</sup> |
| Neutrophils | L/D CD45<sup>+</sup> CD115<sup>-</sup> Gr1<sup>+</sup> |
| Eosinophils | L/D CD45<sup>+</sup> SSC<sup>high</sup> SiglecF<sup>+</sup> |

The antibodies and sorting panels used to purify the specific mature and progenitor cells were as follows: Panel 1 – LSK, CMP, and GMP; Panel 2 – cMoP and Ly6C^hi^ monocytes; Panel 3 – Pre-Neutrophils, Immature Neutrophils, Mature Neutrophils, and Eosinophils.

| **Antibodies and sorting panels used to purify mature myeloid and progenitor cells** |  |
| --- | --- |
| **Cell type** | **Antibodies used in sorting panels** |
| **LSK** | L/D, Lin<sup>-</sup> (CD3, CD4, CD8, B220, NK1.1, Gr1, CD11b), cKit<sup>+</sup>, Sca1<sup>+</sup> |
| **CMP** | L/D, Lin<sup>-</sup> (CD3, CD4, CD8, B220, NK1.1, Gr1, CD11b) cKit<sup>+</sup> Sca1<sup>-</sup> CD34<sup>+</sup> FcγRIII<sup>-</sup> |
| **GMP** | L/D, Lin<sup>-</sup> (CD3, CD4, CD8, B220, NK1.1, Gr1, CD11b) cKit<sup>+</sup> Sca1<sup>-</sup> CD34<sup>+</sup> FcγRIII<sup>+</sup> |
| **cMoP** | L/D, Lin<sup>-</sup> (CD3, CD4, CD8, B220, NK1.1) CD117<sup>+</sup> CD115<sup>+</sup> CD135<sup>-</sup> Ly6C<sup>+</sup> CD11b<sup>-</sup> |
| **Ly6C<sup>hi</sup> monocytes** | L/D, Lin<sup>-</sup> (CD3, CD4, CD8, B220, NK1.1) CD117<sup>-</sup> CD115<sup>+</sup> CD135<sup>-</sup> Ly6C<sup>high</sup> CD11b<sup>+</sup> |
| **Pre-Neutrophil** | L/D, Lin<sup>-</sup> (CD3, CD4, CD8, B220, NK1.1, CD90.2) cKit<sup>int</sup> CD115<sup>-</sup> SiglecF<sup>-</sup> Gr1<sup>+</sup> CD11b<sup>+</sup> CXCR4<sup>int</sup> |
| **Immature Neutrophil** | L/D, Lin<sup>-</sup> (CD3, CD4, CD8, B220, NK1.1, CD90.2) cKit<sup>int</sup> CD115<sup>-</sup> SiglecF<sup>-</sup> Gr1<sup>+</sup> CD11b<sup>+</sup> CXCR4<sup>-</sup> Ly6G CXCR2<sup>-</sup> |
| **Mature Neutrophil** | L/D, Lin<sup>-</sup> (CD3, CD4, CD8, B220, NK1.1, CD90.2) cKit<sup>int</sup> CD115<sup>-</sup> SiglecF<sup>-</sup> Gr1<sup>+</sup> CD11b<sup>+</sup> CXCR4<sup>-</sup> Ly6G CXCR2<sup>-</sup> |
| **Eosinophil** | L/D, CD45<sup>+</sup> SSC<sup>hi</sup> SiglecF<sup>+</sup> |

FACS was performed at the ARA Flow cytometry core facility. Individual cell populations were sorted using BD FACSAria, BD FACSAria Fusion and BD Influx (BD Biosciences). All gating strategies were first set up based on forward scatter area vs. side scatter area, forward scatter height vs. forward scatter area (doublet exclusion) and side scatter area vs. viability dye (viable cell isolation). A sorted event threshold was set to 250,000 and 60,000 cells for human and murine samples, respectively, and cells were sorted according to the expression of the specific surface markers detailed in the preceding tables. Following isolation, cells were washed with PBS w/o Ca^2+^ and Mg^2+^ and stored at -80°C.

### Lipid extraction

Cell samples were lyophilised using either a Savant SpeedVac (Thermo Scientific) or a CoolSafe freeze dryer (ScanVac) prior to extraction and resuspended in 10 µl MilliQ H_2_O. Lipids were extracted using a modified single phase chloroform/methanol extraction method (Weir et al., 2013). Briefly, 200 µl chloroform-methanol (2:1) was added to each sample along with an internal standard (ITSD) mixture containing stable-isotope labelled or non-physiological lipids. In tandem, blank control samples and plasma QCs were extracted and dispersed evenly throughout the extraction order to ensure optimal assay performance and to monitor variation that may arise from the extraction. Samples were subsequently mixed with a rotary mixer for 10 minutes at 90 rpm, sonicated for 30 mins at room temperature and centrifuged at 13,000 rpm for 10 mins to precipitate proteins from the lipid extracts. Supernatant containing the extracted lipids were transferred to a 96 well plate and evaporated using a Savant SpeedVac. Once dried, extracts were reconstituted in H_2_O-saturated butanol and methanol with 10 mM ammonium formate and moved to glass vials and stored until mass spectrometry analysis.

### Liquid chromatography tandem mass spectrometry (LC-MS/MS)

To facilitate better characterisation of lipid species in immune cells, we adapted our previously reported approach (Huynh et al., 2019), optimising the workflow on pooled immune cell samples rather than plasma. Structural elucidation of glycerophospholipids and sphingomyelins were carried out using a combination of conditions as outlined in Huynh et al. 2019. Lipid extracts were analysed using an Agilent 6490 triple quadrupole (QqQ) mass spectrometer coupled to an Agilent 1290 high performance liquid chromatography (HPLC) system and a ZORBAX eclipse plus C18 column (2.1×100mm 1.8µm, Agilent) with thermostat set to 45°C. The final mass spectrometry analysis on each cell population was performed in positive mode with dynamic scheduled MRM. Solvents consisted of solvent A (50% H_2_O, 30% acetonitrile, 20% isopropanol with 10mM ammonium formate) and solvent B (1% H_2_O, 9% acetonitrile, 90% isopropanol with 10mM ammonium formate) and followed a modified 20-minute gradient as follows:

A wash vial comprising of 1:1 butanol:methanol was used after each sample injection. To further improve chromatic peak shape for many anionic/acidic lipid species (notably PS, PA, PIP, S1P), an additional pre-run passivation step was done with phosphoric acid to minimise interaction between the HPLC unit and these lipids.

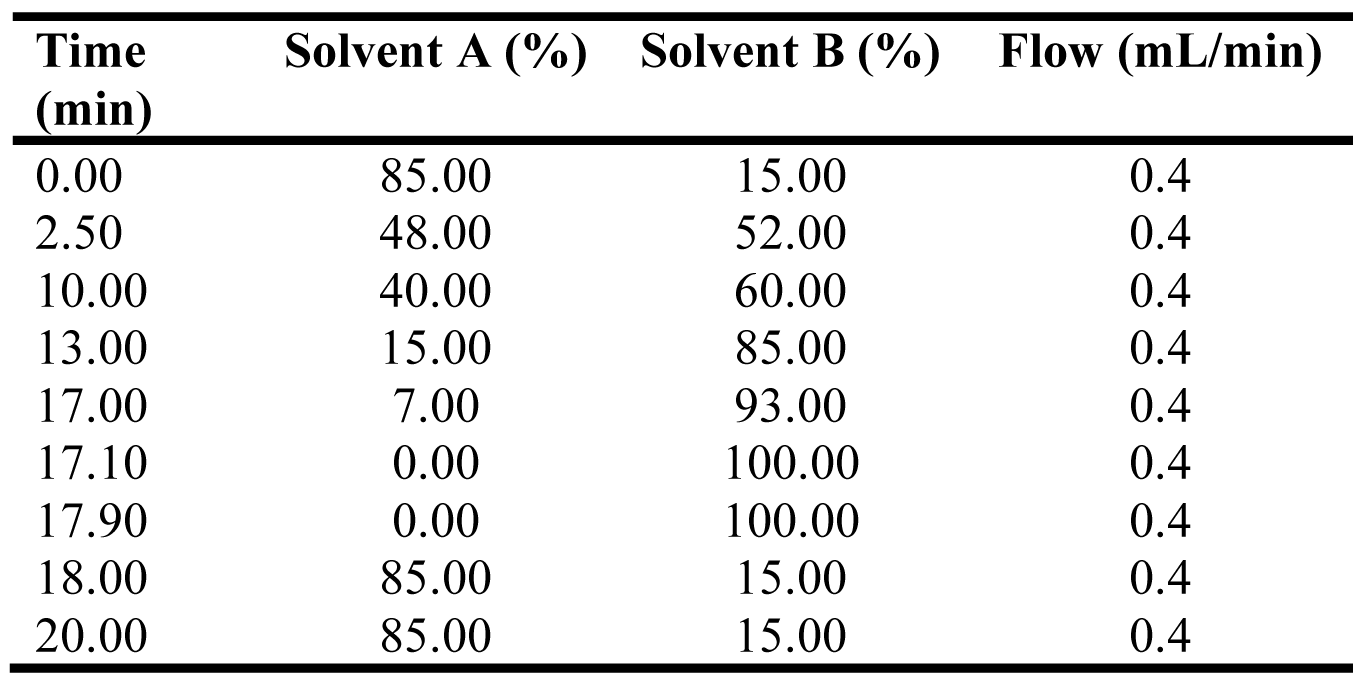

### Lipid nomenclature

The lipid names used follow guidelines set by the LIPIDMAPS consortium and (Fahy et al., 2005; Fahy et al., 2009; Liebisch et al., 2013). Phospholipids with detailed characteristics i.e., acyl chain composition are annotated as [PC(16:0_20:4)] with PC being the lipid class and (16:0_20:4) representing the acyl chains found on the glycerol backbone, irrespective of sn1 or sn2 position. Lipids without specific structural annotations are named based on their sum acyl chain length and degrees of saturation e.g. PC(36:4). Isomeric lipid species separated chromatographically but incompletely annotated were designated (a), (b) etc., with (a) and (b) representing elution order. Owing to technical limitations, we are unable to assign acyl chains to a specific sn1 or sn2 position for the majority of PL species, only that the acyl chain is present at either the sn1 or sn2 position. In the case of ether-PC and PE, the alkyl or alkenyl chains are always located at the sn1 position and the acyl chain always at the sn2 position. In addition, we are unable to determine acyl chain composition for all PLs and the data shown therefore represents those PLs for which we are able to obtain information on their structural composition. Importantly, this represents >90% of total PLs for all cell types, with the exception of neutrophils which was ∼80%, and therefore likely accurately reflects global PL alkyl/acyl chain composition.

### Data normalisation

Analyte areas were obtained from integrating chromatograms that corresponded to a lipid of interest using the Masshunter Quantitative analysis software (Agilent; version B9.00). Lipid concentrations were determined using the following formula:

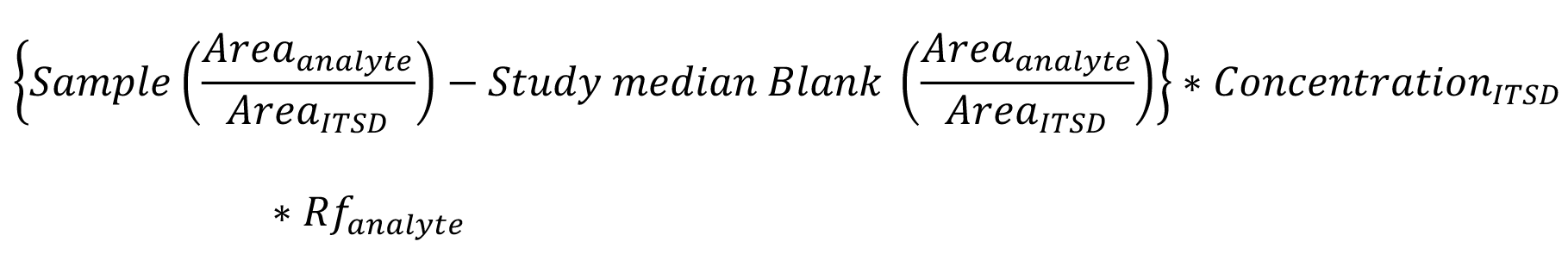

In brief, individual analyte areas were divided by the area of the corresponding internal standards and the median of ITSD containing blank samples was subtracted from each analyte (background subtraction). This value was then multiplied by the ITSD concentration and the individual analyte’s response factor (Rf). Any values that were zeroed after background subtraction as a consequence of being less than the median value of all blank + ITSD samples, were replaced with 1/10^th^ of the minimum value for the corresponding analyte. Data was ultimately normalised to pmol/µmol total lipidome where the background subtracted data for an individual lipid was divided by the sum of the total lipidome and multiplied by a factor of 10^6^ for ease of graphical representation.

### Initial data analysis

To identify any technical outliers within the mouse and human lipid atlas data sets we initially inspected the data sets with a variety of unbiased analytical methods. A heatmap of log-transformed lipid proportions was created for each sample group. Samples were hierarchically clustered using complete linkage on Euclidean distance. Each heatmap was visually inspected for potential outlier samples. Concomitantly, a principal component analysis (PCA) was performed on each sample group, and score plots manually examined to aid outlier identification. Thirdly, the distance to the origin values (distO i.e., the distance of each sample to the origin in PC space), were calculated as a measure of sample extremeness within each PCA, using as many PCs necessary to capture at least 70% of the total variability in the data. Samples with the largest distO values within sample groups were flagged as potential outliers. Next, we examined the lipid data for each sample that was identified as a potential technical outlier. Only samples where a clear technical error in the data could be identified were removed; typically, such samples would contain multiple lipid parameters that were 3-4 standard deviations from both the group mean (i.e. within a given cell type) and from all other cell group means within the atlas. For the human cell atlas, this resulted in the following n number in each group from the initial 14 human donors: [naïve B cell (14); memory B cells (13); CD4 T_naive_ (13); CD4 T_CM_ (14); CD4 T_EM_ (13); CD8 T_naive_ (14); CD8 T_CM_ (12); CD8 T_EM_ (13); CD56^dim^ NK cells (12); CD56^bright^ NK cells (12); classical monocytes (14); intermediate monocytes (14); non-classical monocytes (13); basophils (14); eosinophils (13); neutrophils (11). For the mouse cell atlas, this resulted in the following n number in each group from the initial 10 C57Bl6/J mouse donors: B cells (8); CD4 T cells (9), CD8 T cells (8); NK cells (9); Ly6C^lo^ monocytes (9); Ly6C^hi^ monocytes (10); eosinophils (9); neutrophils (10).

### Data analysis

All downstream analysis was conducted in R version 3.6.2 (AN). PCA was performed using the FactoMineR and factoextra packages using log10 transformed lipid concentrations. Data was centred and scaled prior to PCA. For tSNE analysis datasets were log transformed and unit variance scaled (all lipids have a mean of 0 and a standard deviation of 1). t-SNE was used to embed the lipidomics data on to two dimensions for visualization. One-way ANOVAs followed by post-hoc testing using Tukey’s test were performed using in-built functions in R. ANOVA generated p-values were corrected for multiple comparisons using the Benjamini-Hochberg method for false discovery rate (FDR; 5%) correction. FDR-corrected p values were statistically significant when p < 0.05.

### Ferroptosis studies in mouse immune cells

In all mouse experiments, immune cells were isolated from either wild-type, male C57BI6/J mice or NOX2 knockout mice aged 6-10 weeks old after being humanely killed by CO_2_ asphyxiation. C57Bl6/J mice were bred and housed at the ARA Animal Services Facility (Melbourne, Australia). NOX2 KO mice on a C57Bl6/J background were housed at La Trobe University. All procedures complied with national guidelines for the care and use of laboratory mice and were approved by the appropriate institutional animal ethics committee.

### Cell culture and conditions

Tibia and femurs were isolated from the hind limbs of the mice and bone marrow flushed with RPMI-1640 media containing GLUTAMAX (RPMI; Life Technologies) through a 40 µm filter. Red blood cells were lysed, and white blood cells centrifuged, washed, and resuspended in RPMI supplemented with 5% heat-inactivated foetal bovine serum (FBS; Life Technologies). The total white blood cell count was obtained using an automated haematology analyser (Sysmex XS-1000i). 500,000 bone marrow cells per well were seeded into a 96-well plate (Corning) for incubation with indicated treatments. All cells were maintained in RPMI supplemented with 5% heat-inactivated FBS. Cells were grown in incubators with controlled temperature of 37°C, 5% CO_2_ and 95% humidity.

For FACS-sorting, in addition to the tibias and femurs, spleens were collected. Bone marrow and spleen cells were flushed separately through a 40 µm filter with RPMI to obtain a single cell suspension, RBCs were lysed, and cells were stained on ice with antibodies against CD45, CD11b, CD19, CD3, Ly6C, and Ly6G at 1:400 dilution for 30 mins in the dark on ice. After 30 mins, ∼1 ml of RPMI was added, cells spun (500 g for 5 mins) and the cell pellet resuspended in ∼1ml of RPMI and filtered (40 µm) prior to sorting on a BD FACSAria Fusion flow cytometer at the AMREP flow cytometry core facility (Burnet Institute). Neutrophils, monocytes, and B cells were sorted from the bone marrow and T cells from spleen.

In experiments that use purified cell populations, neutrophils were isolated from bone marrow using EasySep Mouse Neutrophil Enrichment kit (STEMCELL Technologies) and T cells were isolated from the spleen using the MojoSort^TM^ Mouse CD3 T cell Isolation Kit (Biolegend). Experiments where purified T cells were cultured for longer than 24 hours, murine IL-7 (10ng/ml; PeproTech) was supplemented in the medium to prolong survival and reduce basal cell death.

### Ferroptosis experiments

Whole bone marrow cells cultured in RPMI with 5% heat inactivated FBS were treated with ML210 or RSL3 for 6 or 24 h at the concentrations indicated in the specific figures. After treatment, lipid peroxidation and cell viability were assessed using flow cytometry (described below). In experiments using specific inhibitors of ferroptosis, bone marrow cells were treated with either ML210 (1 µM) or RSL3 (2 µM) for 24 h in the absence or presence of Ferrostatin-1 (Fer-1;1 µM), α -tocopherol (α-TOH; 200 µM) or ciclopirox (CPX; 10 µM). In FACS-sorted ferroptosis experiments, sorted neutrophils, monocytes, T cells, and B cells were centrifuged (500 g for 10 mins), resuspended in culture medium and seeded (200k cells/well) into a 96-well plate with varying doses of ML210 or RSL3 for 24 hours, after which cell viability was assessed.

Experiments with combined double or triple inhibition of the GPX4, FSP1 and DHFR pathway were performed in whole bone marrow cells. A dose response of ML210 was performed at the indicated concentrations for 24 h in the presence of either methotrexate (1.5 µM), iFSP1 (3 µM), methotrexate (1.5 µM) and iFSP1 (3 µM), or vehicle control (DMSO). After 24 hours, cells were washed and stained for cell viability. For the inhibition of the DHODH pathway, brequinar sodium (BQR) was diluted to the appropriate concentration (25 µM to 1 mM) in culture medium. Whole bone marrow cells were co-incubated with either ML210 (1 or 10 µM) or vehicle control (DMSO) with different doses of brequinar sodium for 24 hours. In the ferroptosis inhibitor experiments, bone marrow cells were treated with BQR (500 µM) in the presence of the respective ferroptosis inhibitors (Fer-1, 1 µM; α-TOH, 200 µM; or CPX, 10 µM) for 24 hours. For apoptosis and necrostatin inhibitors, cells were pre-treated with 25 µM of Z-VAD(OMe)-FMK (z-VAD) or 10 µM of ()-Necrostatin-2 (Nec-1s) for 1 hour at 37°C, followed with BQR (500 µM) for 24 hours. In the neutrophil activation experiments, purified mouse WT or NOX2 neutrophils were supplemented with PMA (50nM) in the presence of ML210 (5 µM) with or without Fer-1 (1µM) for 6 hours.

### Flow cytometry

Briefly, following treatment in 96-well plates, cells were washed, centrifuged (500 g for 5 mins) and stained in 100 µL of FACS buffer (HBSS with 0.1% BSA w/v, 5 mM EDTA) containing mouse antibodies anti-CD45 (BD Sciences), anti-CD11b (Biolegend), anti-CD19 (Biolegend), anti-CD3 (Biolegend/ Invitrogen), anti-Ly6C (Invitrogen), and anti-Ly6G (Biolegend) at 1:400 dilution for 30 mins in the dark on ice. After 30 mins, cells were washed and transferred to FACS tubes for analysis on an LSRII Fortessa (BD Biosciences) flow cytometer with FACSDiva software. All cells were determined using side scatter area (SSC-A) vs. forward scatter area (FSC-A). Single cells were selected based on SSC-A vs. SSC-height (SSC-H). Leukocytes were selected as CD45^+^, from which myeloid cells (CD11b^+^) and lymphoid cells (CD11b^-^) were gated. Neutrophils were identified from the CD11b^+^ gate as Ly6G^+^Ly6C^-^, while monocytes were Ly6G^-^Ly6C^+^. From the CD11b^-^ gate, T cells were identified as CD3^+^ and B cells as CD19^+^. Upon identification of cell type, cell viability was determined using DAPI and lipid peroxidation assessed by determining the mean fluorescence intensity of Liperfluo and C_11_-BODIPY^481/591^ (both detected in the FITC channel, from separate staining panels). All antibody details can be found in the resource table. Data was analysed using FlowJo.

### Cell viability analysis

The viability of the cultured cells was determined by DAPI (4’,6-Diamidino-2-Phenlindole, Dihydrochloride; BD Pharmingen), a nucleic acid stain that is excluded from viable cells. Immediately prior to flow cytometry analysis, cells were resuspended in FACS buffer containing 0.05 µg/mL DAPI solution and incubated for 5 mins at room temperature in the dark. After incubation, cells were immediately analysed on the flow cytometer. Upon identification of cell type, cell viability was determined using DAPI fluorescence where DAPI^-^ cells were considered viable cells and DAPI^+^ cells were conserved dead cells.

### Lipid peroxidation analysis

Lipid peroxidation was assessed using the fluorescent probes C_11_-BODIPY^581/591^ (4,4-difluoro-5-(4-phenyl-1,3-butadienyl)-4-bora-3a,4a-diaza-s-indacene-3-undecanoic acid; Thermofisher Scientific) and Liperfluo (Dojindo). C_11_-BODIPY^481/591^ is a lipid probe that oxidises in the presence of ROS in cell membranes. Upon oxidation of the polyunsaturated butandineyl component, the dye fluoresces with an emission of ∼510nm (FITC channel). For staining, 2 µM of C_11_-BODIPY^481/591^ was added in the culture medium of cells throughout the incubation period. After treatment, cells were washed and examined by flow cytometry. In the basal state, liperfluo does not fluoresce, but upon reaction with lipid hydroperoxides emits a fluorescence signal with an emission of ∼510nm (FITC channel). Prior to collection of cells, 2 µM of Liperfluo was resuspended into the culture medium for the last 30 min of treatment at 37°C, after which cells were washed with cold FACs buffer, stained for surface markers, and the mean fluorescence intensity of either C_11_-BODIPY^481/591^ or liperfluo determined.

### Lipid supplementation to modulate immune cell lipidomes

Lipid supplementation experiments were performed in purified cell types using enrichment kits. Before the start of the experiment, arachidonic acid (AA 20:4; Sigma-Aldrich), docosahexaenoic acid (DHA 22:6; Sigma-Aldrich) and oleic acid (OA 18:1; Sigma-Aldrich) were solubilised in ethanol to a stock concentration of 1M. Subsequently, the respective fatty acid control (ethanol), 200 µM of OA and 400 µM of DHA and AA were dissolved in filtered (0.2 µM Minisart filter; Sigma-Aldrich) RPMI media containing 5% FBS and 2% w/v bovine serum albumin (BSA; Sigma-Aldrich #A6003). FA mixtures were gently rocked for 1 hour at 37°C to aid conjugation of the fatty acids to BSA. Once prepared, DHA and AA or vehicle control (BSA + ethanol) mixtures were then added to the purified BM neutrophils (250k cells/well) in complete medium with or without Fer-1 (1 µM). After 24 hours of incubation, cell viability and lipid peroxidation levels were assessed using DAPI and liperfluo staining. In the inhibitor experiments, neutrophils were pre-treated with either z-VAD (25 µM) or Nec-1s (10 µM) respectively for 1 hour before the addition of the 400 µM of DHA+AA mixture while Fer-1 (1 µM) was added immediately after the addition of the 400 µM of DHA+AA mixture. For oleic acid treatment in T cells, following the conjugation of OA to BSA, purified spleen CD3^+^ T cells (250k cells/well) were pre-treated with either vehicle control BSA or oleic acid (200 µM) for 16 hours in complete medium supplemented with IL-7 (10 ng/ml) followed by ML210 treatment for 24 hours, after which cell viability and liperfluo fluorescence was assessed. 1-stearoyl-2-oleoyl-sn-glycero-3-phosphoethanolamine (PE18:0/18:1; Avanti) was dissolved in ethanol and sonicated in a water bath for 15 min. Isolated spleen CD3^+^ T cells (250k cells/ well) were treated with either vehicle control (ethanol) or PE 18:0 18:1 (20 µM) in complete medium for 16 hours followed by ML210 treatment for another 24 hours, after which cell viability and liperfluo fluorescence was assessed.

### Lipidomic analysis in ML210 treated immune cells

Sorted cell types (neutrophils, monocytes, T and B cells) were treated with ML210 (1 µM) in the absence or presence of Fer-1 (1 µM). After the experimental period, cells were collected and washed twice in 200 µl of ice-cold PBS (without Ca^2+^ and Mg^2+^) and sonicated using an S-400 sonicator (Misonix) at an amplitude of 13% for 25 seconds. Protein concentration was determined by Quant-iT Protein Assay Kit (Thermo Fisher Scientific, #Q33210) as per manufacturer’s protocol. To prepare samples for lipid extraction, the cell homogenates were stored at -80°C for 2 hours and dried down overnight in a Savant SpeediVac (Thermo Scientific). Sample order was randomised and lipids were extracted using a single-phase butanol/methanol extraction method, as we have described previously (Alshehry et al., 2015). Mass spectrometry analysis was performed as described above. Data were normalised to the total lipidome (sum of all phospholipids, sphingolipids, free cholesterol, and lyso-PLs) and data from individual lipid species are presented as relative to the total lipidome.

### Ferroptosis studies in human immune cells

Peripheral blood samples were collected in lithium heparin coated tubes from healthy male volunteers (21-50 years of age). To lyse red blood cells samples were treated with sterile 1x red blood lysis buffer for 15 mins with frequent vortexes, after which samples were washed in FACS buffer. Neutrophils were isolated using MojoSort™ Whole Blood Human Neutrophil Isolation Kit (Biolegend) and T cells were isolated using MojoSort^TM^ Human CD3 T cell Isolation Kit (Biolegend). Purified human neutrophils and T cells were cultured in Human Plasma-Like Medium (HPLM; ThermoFisher Scientific) with 5% heat inactivated FBS supplemented. Neutrophils and T cells were treated with ML210 for 72 hours in the absence or presence of Fer-1(1 µM), after which lipid peroxidation and cell viability were assessed by Liperfluo fluorescence and DAPI staining (as described above).

### Calculation of the cellular peroxidation index (CPI)

The propensity for a PUFA to be peroxidised is determined by the number of bis-alllyic carbons (CH_2_ centres flanked by double bonds) that are the target of free radical H atom abstraction. We calculated the CPI based on the known relative H atom transfer propagation rate constants of fatty acids in solution, 0.014, 1, 3.2, 4.0, and 5.4 for fatty acids with 1, 2, 4, 5, and 6 double bonds, (Xu et al., 2009; Yin et al., 2011); saturated fatty acids are essentially un-reactive to free radical oxidation at 37°C. Because a linear relationship exists between the H atom transfer propagation rate constants of fatty acids and the number of bis-allylic carbons, we interpolated a relative H atom transfer propagation rate constant for fatty acids with 3 double bonds (2.08) as this was not empirically determined in previous work. To determine the CPI, we firstly calculated the sum of each acyl chain within the total PL acyl chain pool (expressed as a %) multiplied by its corresponding relative H atom transfer propagation rate constant (e.g. 4.56 [the % of 22:6 acyl chains within the total PL acyl chain pool in naïve B cells] x 5.4 [the relative H atom transfer propagation rate constant for fatty acids with 6 double bonds]).

| REAGENT | SOURCE | IDENTIFIER |
| --- | --- | --- |
| **Antibodies** |  |  |
| anti-human CCR7 PB | Biolegend | Cat# 353210 RRID: AB_10918984 |
| anti-human CD123 PE | Biolegend | Cat# 306006 RRID: AB_314580 |
| anti-human CD14 APC/Cy7 | Biolegend | Cat# 367108 RRID: AB_2566710 |
| anti-human CD14 PB | BD Bioscience | Cat# 558121 RRID: AB_397041 |
| anti-human CD15 PE | BD Bioscience | Cat# 562371 RRID: AB_11154049 |
| anti-human CD16 PE/Cy7 | BD Bioscience | Cat# 557744 RRID: AB_396850 |
| anti-human CD19 PE | BD Bioscience | Cat# 555413 RRID: AB_395813 |
| anti-human CD19 PE/Cy5 | Biolegend | Cat# 302210 RRID: AB_314240 |
| anti-human CD2 PE | BD Bioscience | Cat# 555327 RRID: AB_395734 |
| anti-human CD203c FITC | Biolegend | Cat# 324614 RRID: AB_11218991 |
| anti-human CD27 PB | Biolegend | Cat# 356414 RRID: AB_356414 |
| anti-human CD3 FITC | Biolegend | Cat# 300306 RRID: AB_314042 |
| anti-human CD3 PE | Biolegend | Cat# 300308 RRID: AB_314044 |
| anti-human CD4 APC | Biolegend | Cat# 357408 RRID: AB_2565660 |
| anti-human CD45RA PE/Cy7 | Biolegend | Cat# 304126 RRID: AB_10708879 |
| anti-human CD56 PE | BD Bioscience | Cat# 555516 RRID: AB_395906 |
| anti-human CD56 PerCP/Cy5.5 | Biolegend | Cat# 318322 RRID: AB_893389 |
| anti-human CD66b PE/Cy7 | Biolegend | Cat# 305116 RRID: AB_2566605 |
| anti-human CD8 PE | Biolegend | Cat# 300908 RRID: AB_314112 |
| anti-human HLA-DR APC | Biolegend | Cat# 327022 RRID: AB_2734311 |
| anti-human Nkp46 PE | BD Bioscience | Cat# 557991 RRID: AB_396974 |
| anti-human Siglec8 APC | Biolegend | Cat# 347106 RRID: AB_2561402 |
| anti-mouse B220 FITC | eBioScience | Cat# 11-0451-82 RRID: AB_465054 |
| anti-mouse CD115 APC | Biolegend | Cat# 135510 RRID: AB_2085221 |
| anti-mouse CD115 BV605 | Biolegend | Cat# 135517 RRID: AB_2562760 |
| anti-mouse CD11b FITC | Biolegend | Cat# 101206 RRID: AB_312789 |
| anti-mouse CD11b PE | BD Bioscience | Cat# 553311 RRID: AB_396680 |
| anti-mouse CD11b PE/Cy7 | Biolegend | Cat# 101216 RRID: AB_312799 |
| anti-mouse CD135 BV421 | Biolegend | Cat# 135313 RRID: AB_2562338 |
| anti-mouse CD19 APC | Biolegend | Cat# 115512 RRID: AB_313647 |
| anti-mouse CD19 APC | Biolegend | Cat# 115512 RRID: AB_313647 |
| anti-mouse CD3 APC/Cy7 | Biolegend | Cat# 100222 RRID: AB_2242784 |
| anti-mouse CD3 FITC | Biolegend | Cat# 100204 RRID: AB_312661 |
| anti-mouse CD3 PE | Biolegend | Cat# 100206 RRID: AB_312663 |
| anti-mouse CD34 PE | Biolegend | Cat# 128610 RRID: AB_2074601 |
| anti-mouse CD4 FITC | Biolegend | Cat# 100510 RRID: AB_312713 |
| anti-mouse CD4 PE | BD Bioscience | Cat# 550630 RRID: AB_393790 |
| anti-mouse CD45 BUV395 | BD Bioscience | Cat# 564279 RRID: AB_2651134 |
| anti-mouse CD45 PB | Biolegend | Cat# 103126 RRID: AB_493535 |
| anti-mouse CD45 PE | Biolegend | Cat# 103106 RRID: AB_312971 |
| anti-mouse CD8 FITC | eBioScience | Cat# MA1-10303 RRID: AB_11153636 |
| anti-mouse CD8 PerCP | BD Bioscience | Cat# 553036 RRID: AB_394573 |
| anti-mouse CD90.2 FITC | Biolegend | Cat# 140304 RRID: AB_10642812 |
| anti-mouse cKit APC/Cy7 | Biolegend | Cat# 105812 RRID: AB_313221 |
| anti-mouse cKit V450 | BD Bioscience | Cat# 560588 RRID: AB_1645265 |
| anti-mouse CXCR2 PE | Biolegend | Cat# 149304 RRID: AB_2565692 |
| anti-mouse CXCR4 AF647 | Biolegend | Cat# 146504 RRID: AB_2562591 |
| anti-mouse FcγRIII PerCP | BD Bioscience | Cat# 560540 RRID: AB_1645259 |
| anti-mouse Gr1 FITC | BD Bioscience | Cat# 553127 RRID: AB_394643 |
| anti-mouse Gr1 PerCP | BD Bioscience | Cat# 552093 RRID: AB_394334 |
| anti-mouse Ly6C BV785 | Biolegend | Cat# 128041 RRID: AB_2565852 |
| anti-mouse Ly6C PerCP | Biolegend | Cat# 128028 RRID: AB_10897805 |
| anti-mouse Ly6G APC/Cy7 | Biolegend | Cat# 127623 RRID: AB_10645331 |
| anti-mouse NK1.1 FITC | BD Bioscience | Cat# 561082 RRID: AB_394676 |
| anti-mouse Sca1 FITC | Biolegend | Cat# 122506 RRID: AB_756191 |
| anti-mouse Sca1 PB | Biolegend | Cat# 108120 RRID: AB_493273 |
| anti-mouse SiglecF BV605 | BD Bioscience | Cat# 740388 RRID: AB_2740118 |
| anti-mouse SiglecF PE | BD Bioscience | Cat# 562068 RRID: AB_394341 |
| anti-mouse Ter119 FITC | eBioScience | Cat# MA5-17821 RRID: AB_2539205 |
| Liperfluo | Dojindo | Cat# L248; CAS: 1448846-35-2 |
| BODIPY™ 581/591 C11 | ThermoFisher | Cat# D3861 |
| DAPI solution | BD Pharmingen | Cat# 564907 RRID: AB_2869624 |
| Ghost dye (viability) | Tonbo Biosciences | Cat# 13-0870 |
| MitoTracker Green | Invitrogen | Cat# M7514 |
| **Chemicals, fatty acids, and kits** |  |  |
| Erastin | Sigma-Aldrich | Cat# E7781 |
| ML210 | Sigma-Aldrich | Cat# SML0521 |
| Alpha-Tocopherol | Sigma-Aldrich | Cat# T3251 |
| Ferostatin-1 | Sigma-Aldrich | Cat# SML0583 |
| Ciclopirox | Cayman Chemical | Cat# 16021 |
| 1S,3R-RSL3 | Sigma-Aldrich | Cat# SML2234 |
| Brequinar sodium | Tocris | Cat# 6196 |
| Methotrexate | Sigma-Aldrich | Cat# M9929 |
| iFSP1 | Cayman Chemical | Cat# 29483 |
| Z-VAD(OMe)-FMK | Cayman Chemical | Cat# 14463 |
| (-)-Necrostatin-2 | Cayman Chemical | Cat# 11657 |
| SPRi3 | Tocris | Cat# 6597 |
| Idebenone | Sigma-Aldrich | Cat# I5659 |
| Murine IL-7 | PeproTech | Cat# 217-17-50 |
| PE 18:0 18:1 | Avanti | Cat# 850758P |
| Oleic acid | Sigma-Aldrich | Cat# O1008 |
| Arachidonic acid | Sigma-Aldrich | Cat# A3611 |
| Docosahexaenoic acid | Sigma-Aldrich | Cat# D2534 |
| EasySep™ Mouse Neutrophil Enrichment Kit | STEMCELL Technologies | Cat# 19762 |
| MojoSort™ Mouse CD3 T cell Isolation Kit | Biolegend | Cat# 480024 |
| MojoSort™ Human CD3 T cell Isolation Kit | Biolegend | Cat# 480131 |
| MojoSort™ Whole Blood Human Neutrophil Isolation Kit | Biolegend | Cat# 480152 |
| Human Plasma-Like Medium | ThermoFisher | Cat# A4899101 |

