## Extended data for "An immune cell lipid atlas reveals the basis of susceptibility to ferroptosis"

#### Extended Data Figure 1

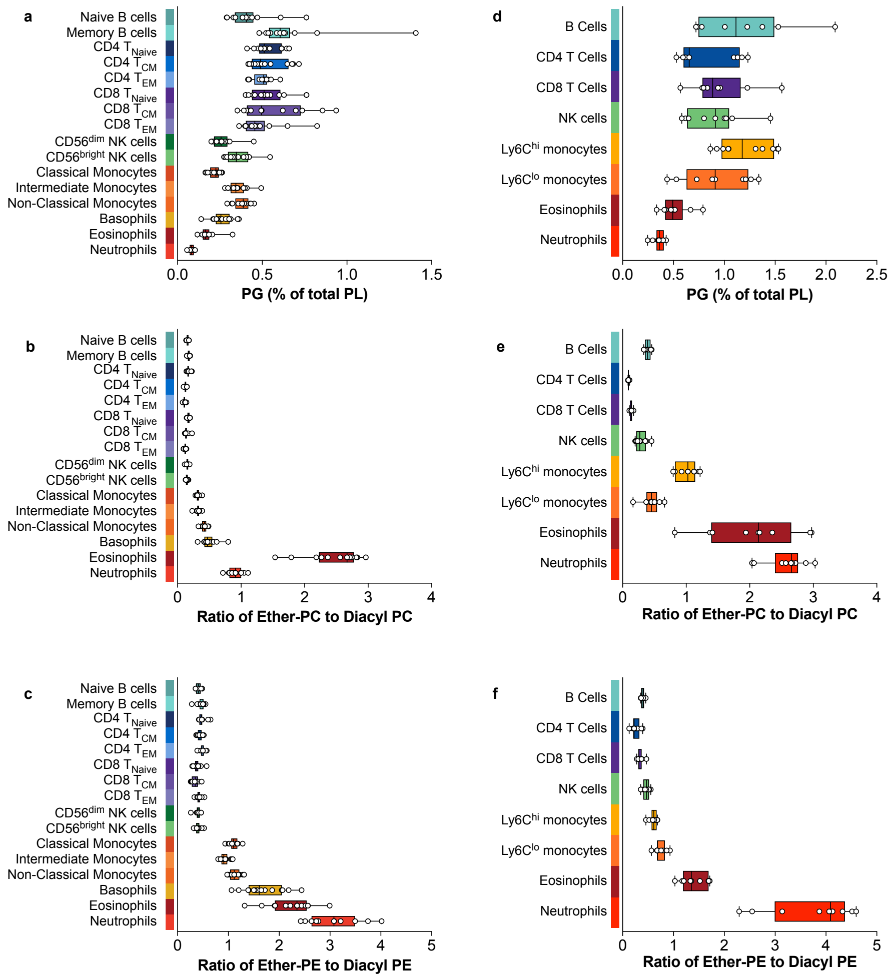


**Extended Data Figure 1. Changes in PG and the ether to diacyl ratio for PC and PE.** **a,d,** PG levels as a % of total PL in human (**a**) and mouse (**d**) immune cells. **b,c,e,f,** Ratio of ether PC to diacyl PC (**b,e**) and ether PE to diacyl PE (**c,f**) in human (**b,c**) and mouse (**e,f**) immune cells. **a-c,** n=11-14 individual donors for each cell type; **d-f,** n=8-10 individual mice for each cell type. Data are shown as a box and whiskers, with the lower and upper limits of the box corresponding to the 25th and 75th percentile, the line within the box being the median, and the whiskers extending to the minimum and maximum values. Biological replicates from individual human donors or mice are shown. P values for PG pairwise comparisons are shown in Supplementary Excel files 1 and 2.

### Extended Data Figure 2

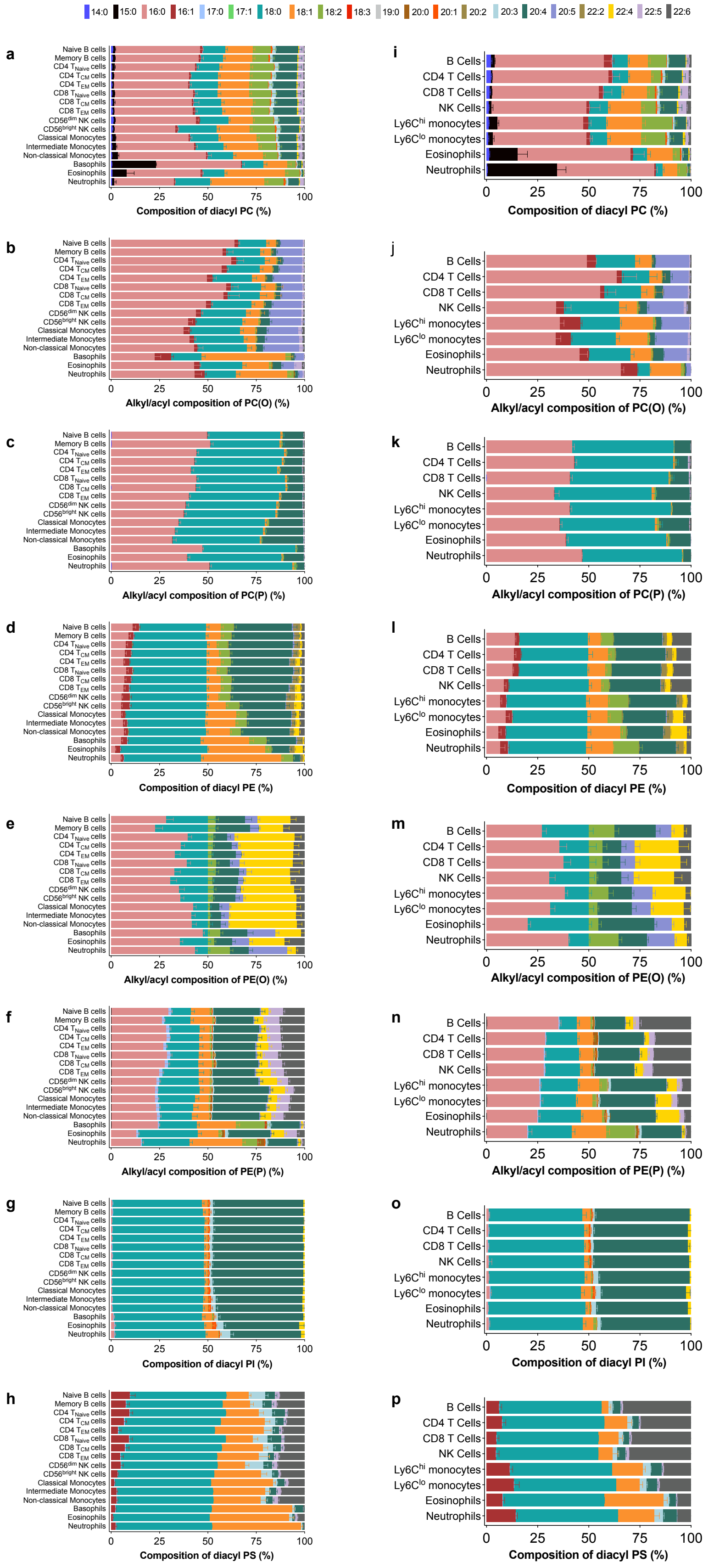

**Extended Data Figure 2. Changes in alkyl/acyl composition within specific PL subclasses. a-p,** Alkyl/acyl composition for the indicated PL subclasses in human (**a-h**) and mouse (**i-p**) immune cells. Data are shown as mean + S.D. from n=11-14 individual human donors (**a-h**) or n=8-10 individual mice (**i-p**). In each figure, statistically significant differences in lipid features between cell types was determined by 1-way ANOVA with Tukey's HSD test after false discovery rate correction (5%; Benjamini-Hochberg). P values for all pairwise comparisons are shown in Supplementary Excel files 1 and 2.

**Extended Data Figure 3**

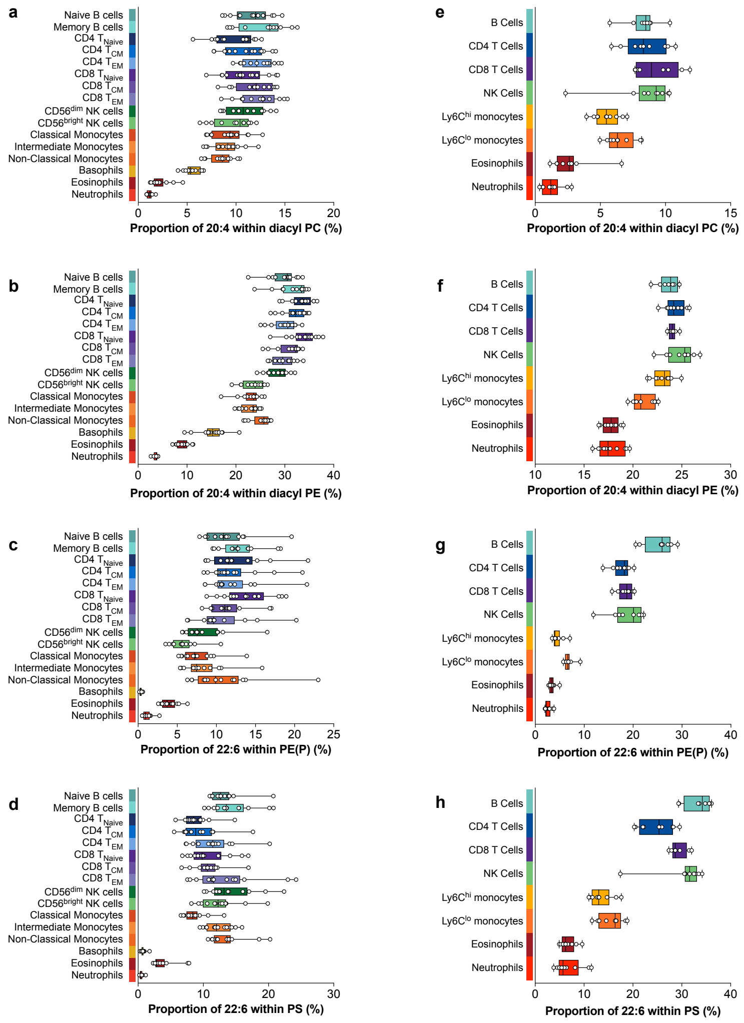


**Extended Data Figure 3. Levels of PUFA-containing PL vary between immune cell types.**

**a-h,** Levels of specific lipid features in human (**a-d**) and mouse (**e-h**) immune cells. **a-d,** n=11-14 individual donors for each cell type; **e-h,** n=8-10 individual mice for each cell type. Data are shown as a box and whiskers, with the lower and upper limits of the box corresponding to the 25$^{th}$ and 75$^{th}$ percentile, the line within the box being the median, and the whiskers extending to the minimum and maximum values. Biological replicates from individual human donors or mice are shown. P values for pairwise comparisons are shown in Supplementary Excel files 1 and 2.

#### Extended Data Figure 4

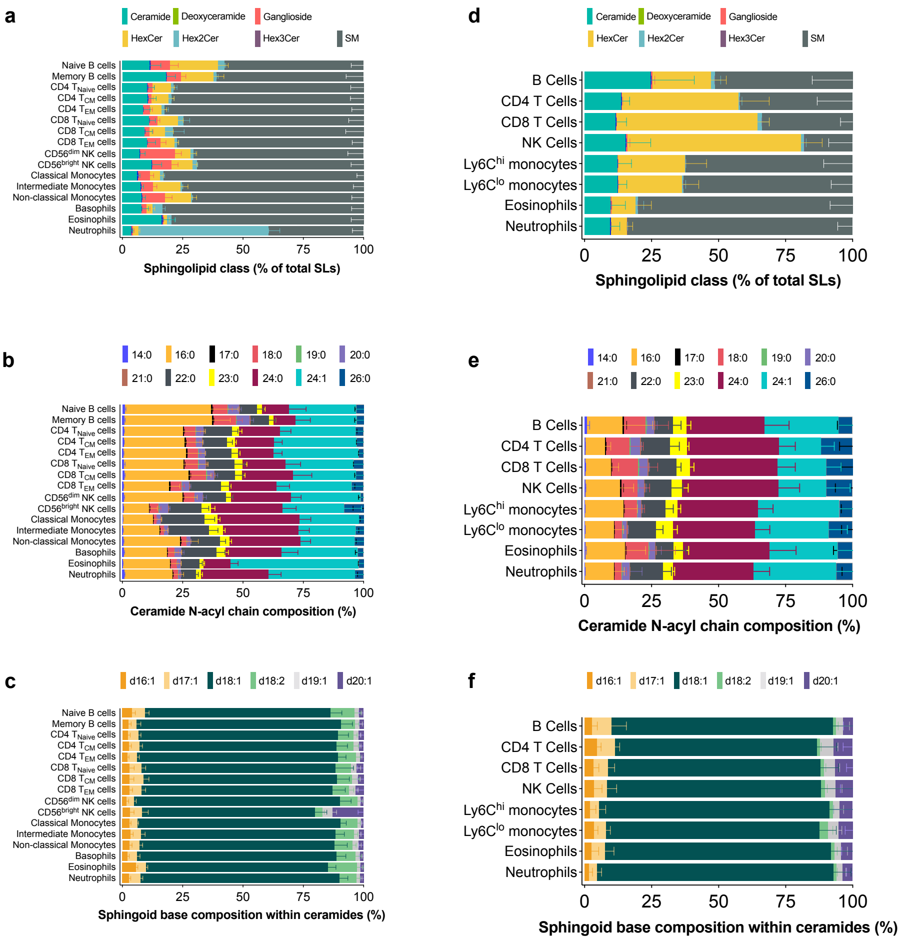


**Extended Data Figure 4. Sphingolipid changes in the cells of the human and mouse immune system.**

**a-f,** Breakdown of the indicated sphingolipid features in human (**a-c**) and mouse (**d-f**) immune cells. Data are shown as mean + S.D. from n=11-14 individual human donors (**a-c**) or n=8-10 individual mice (**d-f**). In each figure, statistically significant differences in lipid features between cell types was determined by 1-way ANOVA with Tukey's HSD test after false discovery rate correction (5%; Benjamini-Hochberg). P values for all pairwise comparisons are shown in Supplementary Excel files 1 and 2.

### Extended Data Figure 5

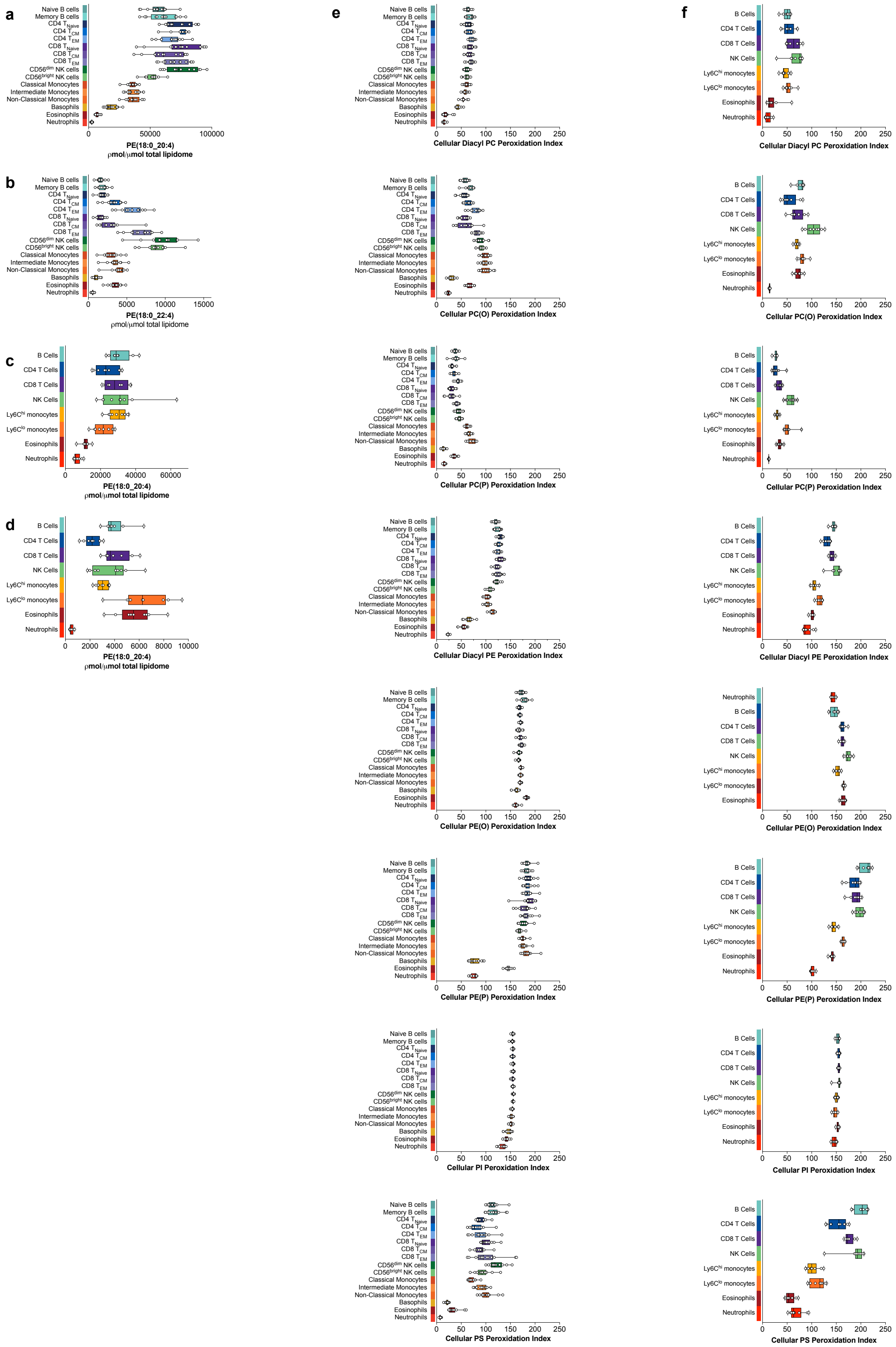

**Extended Data Figure 5. The lipidome of myeloid cells, most notably neutrophils, has a reduced potential for lipid peroxidation.**

**a-d,** Levels of canonical ferroptosis-inducing PL in human (**a,b**) and mouse (**c,d**) immune cells. **e,f,** Lipid peroxidation indexes of PL subclasses calculated based on the relative susceptibility of different fatty acids to lipid peroxidation. **a,b,e,** n=11-14 individual donors for each cell type; **c,d,f,** n=8-10 individual mice for each cell type. Data are shown as a box and whiskers, with the lower and upper limits of the box corresponding to the $25^{th}$ and $75^{th}$ percentile, the line within the box being the median, and the whiskers extending to the minimum and maximum values. Biological replicates from individual human donors or mice are shown.

#### Extended Data Figure 6

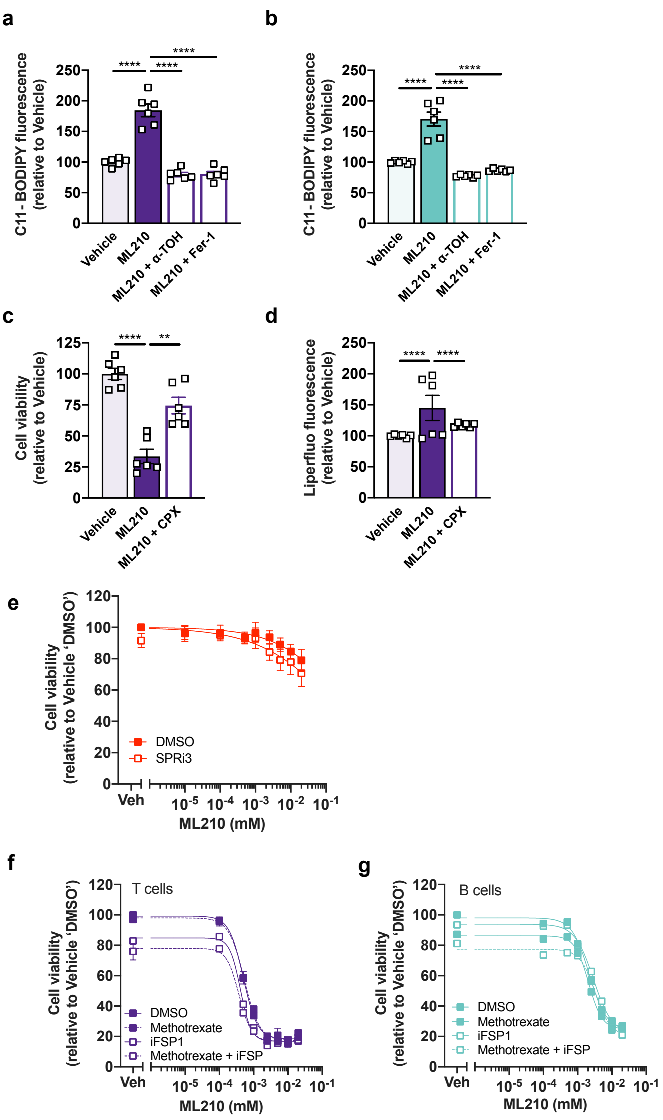


**Extended Data Figure 6.**

**a,b,** C11-BODIPY fluorescence in T cells (**a**) and B cells (**b**) treated with vehicle (DMSO), ML210 (1 μM) alone or ML210 in combination with either α-tocopherol (α-TOH; 200 μM) or ferrostatin (Fer-1; 1 μM) for 24 h. n=6 biological replicates. **c,d,** Cell viability (**c**) and liperfluo fluorescence (**d**) in T cells treated vehicle (DMSO), ML210 (1 μM) alone or ML210 in combination with ciclopirox (CPX; 10 μM) for 24 h. n=6 biological replicates. Data was analysed using a 1-way ANOVA with Tukey's HSD test. *$p<0.05$, **$p<0.01$, ***$p<0.001$, and

****p < 0.0001. **e,** Cell viability in bone marrow neutrophils treated with the indicated doses of ML210 for 24 h following pre-treatment with sepiapterin reductase inhibitor 3 (SPRi3; 50μM) for 16 h. n=5-6 biological replicates. **f,g,** Cell viability of T cells (**f**) and B cells (**g**) treated with the indicated doses of ML210 alone or in combination with methotrexate (1.5 μM), ferroptosis suppressor protein 1 inhibitor (iFSP1; 3 μM), or methotrexate + iFSP1 for 24 h. n=5-6 biological replicates.

### Extended Data Figure 7

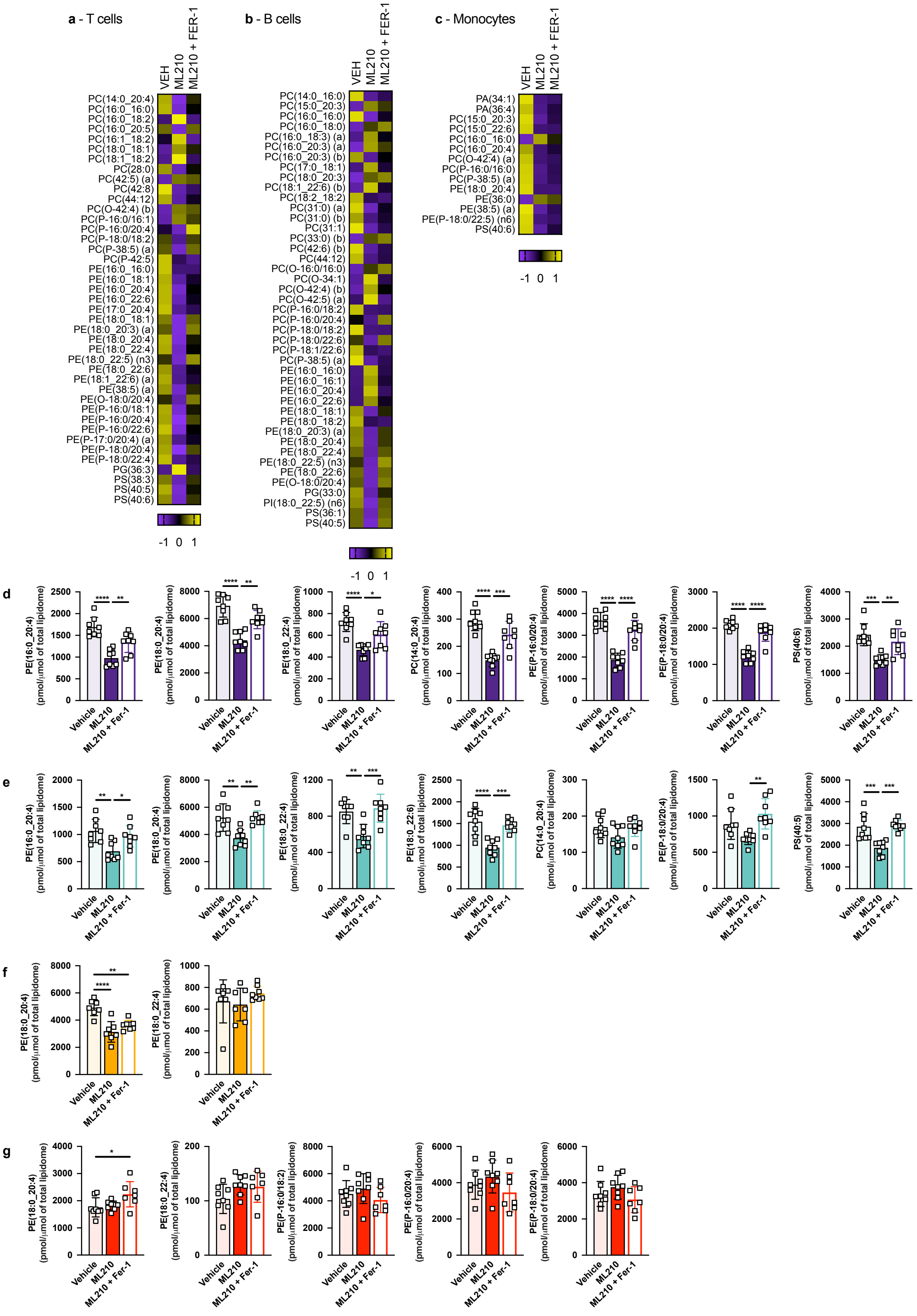

**Extended Data Figure 7. ML210 treatment in T cells, B cells, and monocytes differentially influences phospholipid composition.**

**a-c,** Heat maps showing significantly different phospholipid species in T cells (**a**), B cells (**b**) and monocytes (**c**) treated with vehicle (DMSO), ML210 (1 μM) or ML210 + Ferrostatin-1 (Fer-1; 1 μM) for 24 h. The phospholipid species shown are those that were significantly different (1-way ANOVA with Tukey's HSD test) after false discovery rate correction (Benjamini-Hochberg). Data are presented as z-scores. n=7-8 biological replicates. **d-g,** Individual PUFA-containing phospholipid species in FACS-sorted T cells (**d**), B cells (**e**), monocytes (**f**), and neutrophils (**g**) treated with vehicle (DMSO), ML210 (1 μM), or ML210 (1 μM) + Ferrostatin-1 (Fer-1; 1 μM) for 24 hours. Data was analysed using a 1-way ANOVA with Tukey's HSD test. *p<0.05, **p<0.01, ***p<0.001, and ****p < 0.0001. n=7-8 biological replicates.

#### Extended Data Figure 8

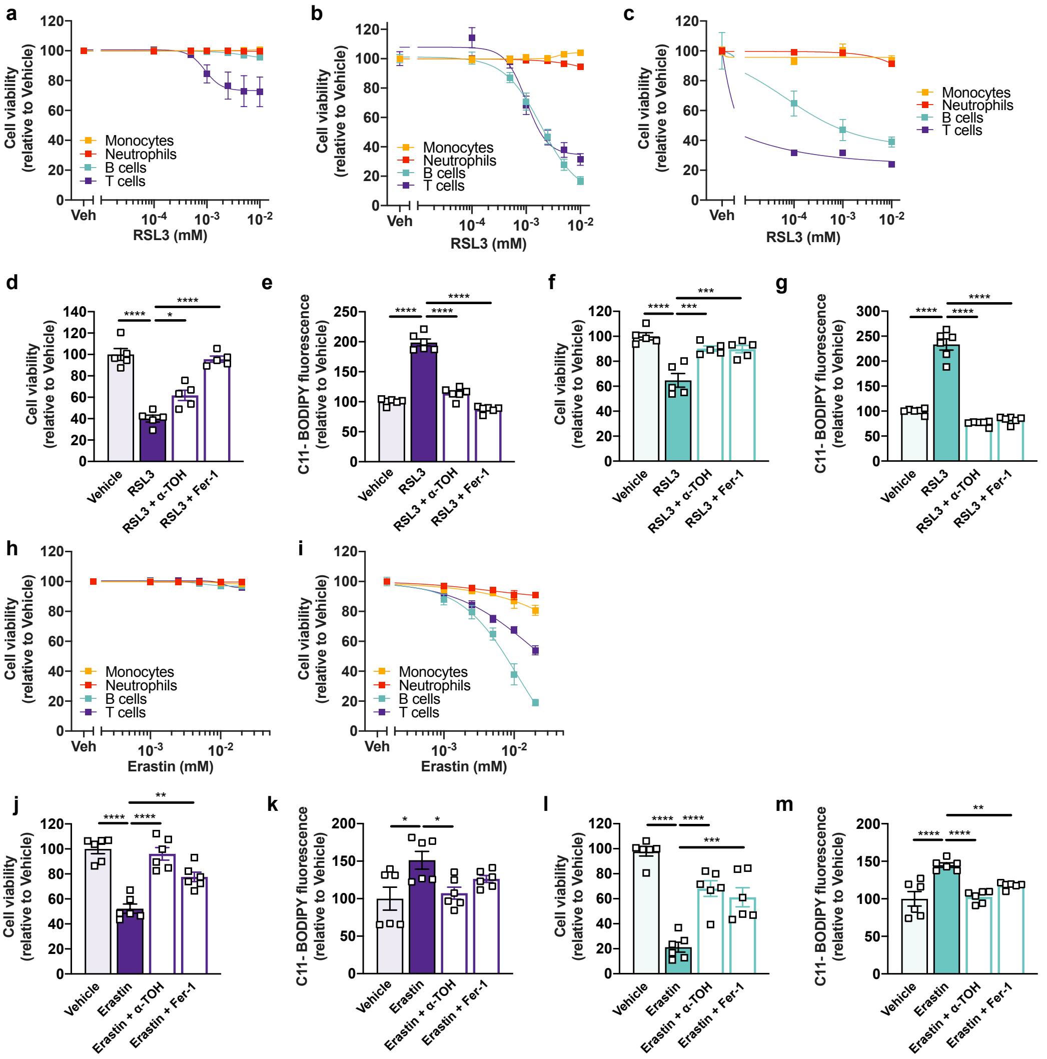


**Extended Data Figure 8. Lymphoid cells are susceptible while myeloid cells are resistant to RSL3- and erastin-induced ferroptosis.**

**a,b,** Cell viability of bone marrow neutrophils, monocytes, T and B cells treated with RSL3 at the indicated doses for either 6 h (**a**) or 24 h (**d**). **c,** Cell viability of FACS-sorted neutrophils, monocytes, T and B cells treated with the indicated doses of RSL3 for 24 h. **d-g,** Cell viability and C11-BODIPY fluorescence in T cells (**d,e**) and B cells (**f,g**) treated with vehicle (DMSO), RSL3 (2 μM) alone, or in combination with either α-TOH (200 μM) or Fer-1 (1 μM) for 24 h. **h,i,** Cell viability of bone marrow neutrophils, monocytes, T and B cells treated with erastin at the indicated doses for either 6 h (**h**) or 24 h (**i**). **j-m,** Cell viability and C11-BODIPY fluorescence in T cells (**j,k**) and B cells (**l,m**) treated with vehicle (DMSO), erastin (5 µM) alone, or in combination with either α-TOH (200 µM) or Fer-1 (1 µM) for 24 h. (**a-m**) n=6 biological replicates. Data was analysed using a 1-way ANOVA with Tukey's HSD test. *$p<0.05$, **$p<0.01$, ***$p<0.001$, and ****$p < 0.0001$. Data are shown as mean ± S.E.M.

#### Extended Data Figure 9

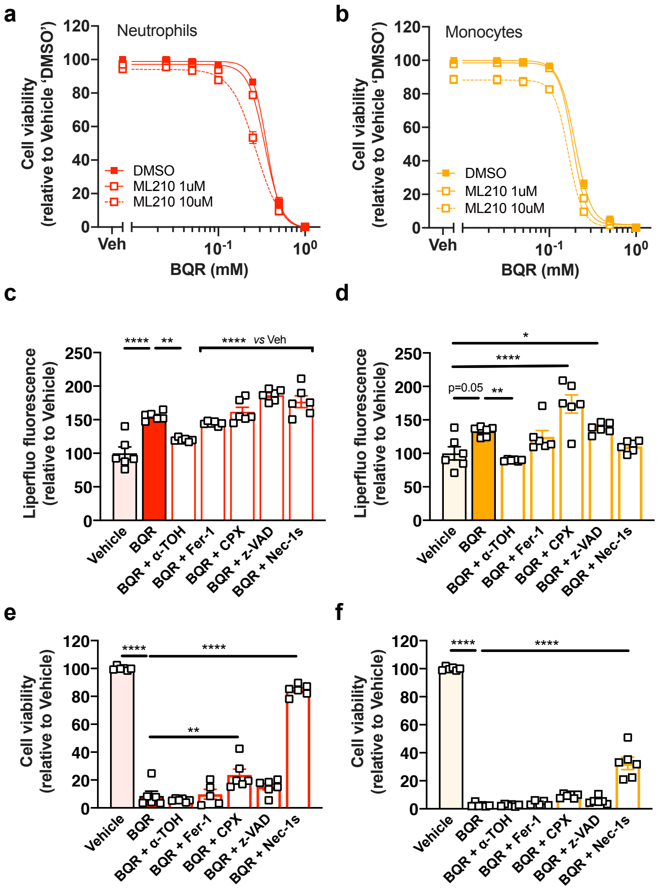


**Extended Data Figure 9. Inhibition of the DHODH pathway does not sensitize myeloid cells to lipid peroxidation and ferroptosis.**

**a,b,** Cell viability in bone marrow neutrophils (**a**) and monocytes (**b**) treated with brequniar (BQR) in the presence of ML210 (1 or 10 μM) or DMSO for 24 h. n=3-6 biological replicates. **c-f,** Liperfluo fluorescence and cell viability in bone marrow neutrophils (**c,e**) and monocytes (**d,f**) treated with BQR (500 μM) in the presence of α-TOH (200 μM), Fer-1 (1 μM), CPX (10 μM) or DMSO for 24 h or following pre-treatment with z-VAD (25 μM) or Nec-1s (10 μM) for an hour. n=5-6 biological replicates. Data was analysed using a 1-way ANOVA with Tukey's HSD test. *p<0.05, **p<0.01, ***p<0.001, and ****p < 0.0001. Data are shown as mean ± S.E.M.

#### Extended Data Figure 10

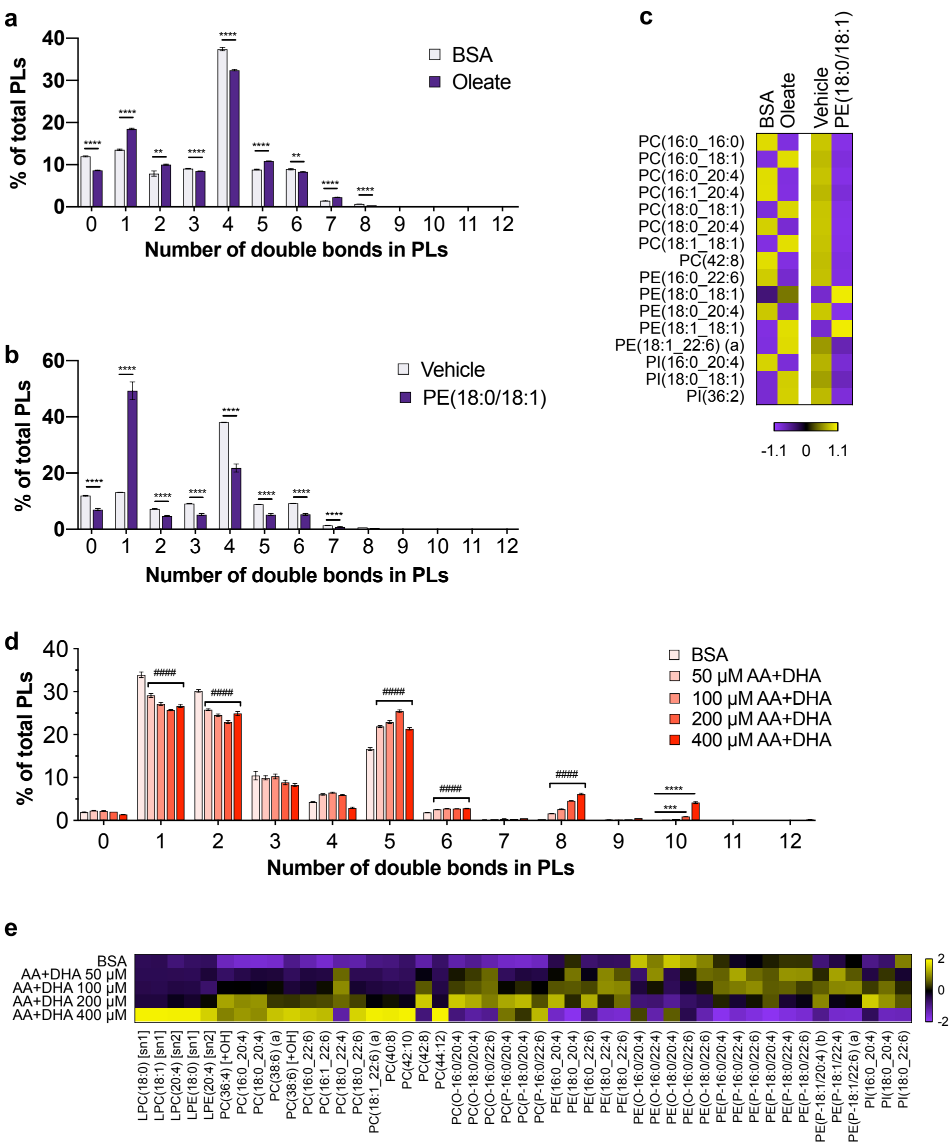


**Extended Data Figure 10. Lipid supplementation in T cells and neutrophils alters PL carbon-carbon double bond composition.**

**a,b,** Percentage of PLs with the indicated number of carbon-carbon double bonds following treatment of purified T cells with oleate (**a,** n=6 biological replicates) or PE(18:0/18:1) (**b,** n=5-6 biological replicates). **c,** Heatmap showing the main PL species altered in purified T cells following either oleate or PE(18:0/18:1) treatment. **d,** Percentage of PLs with the indicated number of carbon-carbon double bonds following treatment of purified neutrophils with the indicated doses of AA + DHA (n=4 biological replicates). **e,** Heatmap showing the main PL species altered in purified neutrophils following treatment with the indicated doses of AA + DHA. Data was analysed using either un-paired student's t-test (**a,b**) or a 1-way ANOVA with Tukey's HSD test (**d**). **p<0.01, ***p<0.001, and ****p < 0.0001 indicate significant differences between the indicated groups. #### p<0.00001 indicates a significant difference between BSA and all of the AA + DHA treated groups. **a,b,d,** Data are shown as mean ± S.E.M. **c,d,** Data are shown as z-scores.
