## Supplementary material for "An immune cell lipid atlas reveals the basis of susceptibility to ferroptosis": Human statistics

| Lipid class | **PE acyl chain** |
|---|---|
| Feature | 14:0 |
| One-way ANOVA *p-value* | 1.0 x $10^{-1}$ |
| FDR adjuested *p-value* | 1.0 x $10^{-1}$ |

| | Naive B | Memory B |
|---|---|---|
| Naive B | # | |
| Memory B | 1.0 x $10^{-1}$ | # |
| CD4 T Naive | 1.0 x $10^{-1}$ | 1.0 x $10^{-1}$ |
| CD4 T Central Memory | 1.0 x $10^{-1}$ | 1.0 x $10^{-1}$ |
| CD4 T Effector Memory | 1.0 x $10^{-1}$ | 1.0 x $10^{-1}$ |
| CD8 T Naive | 1.0 x $10^{-1}$ | 1.0 x $10^{-1}$ |
| CD8 T Central Memory | 1.0 x $10^{-1}$ | 1.0 x $10^{-1}$ |
| CD8 T Effector Memory | 1.0 x $10^{-1}$ | 1.0 x $10^{-1}$ |
| CD56 Dim NK | 1.0 x $10^{-1}$ | 1.0 x $10^{-1}$ |
| CD56 Bright NK | 1.0 x $10^{-1}$ | 1.0 x $10^{-1}$ |
| Classical Monocyte | 1.0 x $10^{-1}$ | 1.0 x $10^{-1}$ |
| Intermediate Monocyte | 1.0 x $10^{-1}$ | 1.0 x $10^{-1}$ |
| Non-classical Monocyte | 1.0 x $10^{-1}$ | 1.0 x $10^{-1}$ |
| Basophil | 1.0 x $10^{-1}$ | 1.0 x $10^{-1}$ |
| Eosinophil | 1.0 x $10^{-1}$ | 1.0 x $10^{-1}$ |
| Neutrophil | 1.0 x $10^{-1}$ | 1.0 x $10^{-1}$ |

| Lipid class | **PE acyl chain** |
|---|---|
| Feature | 15:0 |
| One-way ANOVA *p-value* | 1.0 x $10^{-1}$ |
| FDR adjuested *p-value* | 1.0 x $10^{-1}$ |

| | | |
|---|---|---|
| Naive B | # | |
| Memory B | 1.0 x $10^{-1}$ | # |
| CD4 T Naive | 1.0 x $10^{-1}$ | 1.0 x $10^{-1}$ |
| CD4 T Central Memory | 1.0 x $10^{-1}$ | 1.0 x $10^{-1}$ |
| CD4 T Effector Memory | 1.0 x $10^{-1}$ | 1.0 x $10^{-1}$ |
| CD8 T Naive | 1.0 x $10^{-1}$ | 1.0 x $10^{-1}$ |
| CD8 T Central Memory | 1.0 x $10^{-1}$ | 1.0 x $10^{-1}$ |
| CD8 T Effector Memory | 1.0 x $10^{-1}$ | 1.0 x $10^{-1}$ |
| CD56 Dim NK | 1.0 x $10^{-1}$ | 1.0 x $10^{-1}$ |
| CD56 Bright NK | 1.0 x $10^{-1}$ | 1.0 x $10^{-1}$ |
| Classical Monocyte | 1.0 x $10^{-1}$ | 1.0 x $10^{-1}$ |
| Intermediate Monocyte | 1.0 x $10^{-1}$ | 1.0 x $10^{-1}$ |
| Non-classical Monocyte | 1.0 x $10^{-1}$ | 1.0 x $10^{-1}$ |
| Basophil | 1.0 x $10^{-1}$ | 1.0 x $10^{-1}$ |
| Eosinophil | 1.0 x $10^{-1}$ | 1.0 x $10^{-1}$ |

| | Naive B | Memory B |
|---|---|---|
| Neutrophil | 1.0 x $10^{-1}$ | 1.0 x $10^{-1}$ |

| Lipid class | **PE acyl chain** |
|---|---|
| Feature | 16:0 |
| One-way ANOVA *p-value* | 6.2 x $10^{-65}$ |
| FDR adjuested *p-value* | 1.6 x $10^{-64}$ |

| | Naive B | Memory B |
|---|---|---|
| Naive B | # | |
| Memory B | 1.0 x $10^{-6}$ | # |
| CD4 T Naive | 8.3 x $10^{-14}$ | 1.4 x $10^{-4}$ |
| CD4 T Central Memory | < 1.0 x $10^{-16}$ | 1.4 x $10^{-4}$ |
| CD4 T Effector Memory | < 1.0 x $10^{-16}$ | 3.4 x $10^{-9}$ |
| CD8 T Naive | 8.9 x $10^{-14}$ | 1.6 x $10^{-1}$ |
| CD8 T Central Memory | < 1.0 x $10^{-16}$ | 2.5 x $10^{-5}$ |
| CD8 T Effector Memory | < 1.0 x $10^{-16}$ | 6.9 x $10^{-10}$ |
| CD56 Dim NK | < 1.0 x $10^{-16}$ | 9.2 x $10^{-14}$ |
| CD56 Bright NK | < 1.0 x $10^{-16}$ | 8.4 x $10^{-14}$ |
| Classical Monocyte | < 1.0 x $10^{-16}$ | 5.9 x $10^{-14}$ |
| Intermediate Monocyte | < 1.0 x $10^{-16}$ | 4.1 x $10^{-11}$ |
| Non-classical Monocyte | < 1.0 x $10^{-16}$ | 9.9 x $10^{-8}$ |
| Basophil | < 1.0 x $10^{-16}$ | 7.4 x $10^{-14}$ |
| Eosinophil | < 1.0 x $10^{-16}$ | < 1.0 x $10^{-16}$ |
| Neutrophil | < 1.0 x $10^{-16}$ | 7.3 x $10^{-14}$ |

| Lipid class | **PE acyl chain** |
|---|---|
| Feature | 16:1 |
| One-way ANOVA *p-value* | 2.1 x $10^{-38}$ |
| FDR adjuested *p-value* | 3.0 x $10^{-38}$ |

| | Naive B | Memory B |
|---|---|---|
| Naive B | # | |
| Memory B | 3.6 x $10^{-2}$ | # |
| CD4 T Naive | 1.0 x $10^{-1}$ | 6.1 x $10^{-1}$ |
| CD4 T Central Memory | 1.0 x $10^{-1}$ | 6.1 x $10^{-1}$ |
| CD4 T Effector Memory | 8.0 x $10^{-1}$ | 1.0 x $10^{-1}$ |
| CD8 T Naive | 1.0 x $10^{-1}$ | 4.5 x $10^{-1}$ |
| CD8 T Central Memory | 1.0 x $10^{-1}$ | 1.8 x $10^{-1}$ |
| CD8 T Effector Memory | 8.3 x $10^{-1}$ | 1.0 x $10^{-1}$ |
| CD56 Dim NK | 5.1 x $10^{-3}$ | 2.5 x $10^{-10}$ |

| | Naive B | Memory B |
|---|---|---|
| CD56 Bright NK | 8.3 x $10^{-4}$ | 1.7 x $10^{-11}$ |
| Classical Monocyte | 2.8 x $10^{-8}$ | 1.8 x $10^{-1}$ |
| Intermediate Monocyte | 2.1 x $10^{-7}$ | 3.9 x $10^{-1}$ |
| Non-classical Monocyte | 1.5 x $10^{-7}$ | 3.0 x $10^{-1}$ |
| Basophil | 7.4 x $10^{-1}$ | 1.0 x $10^{-1}$ |
| Eosinophil | 2.0 x $10^{-2}$ | 1.0 x $10^{-1}$ |
| Neutrophil | 9.0 x $10^{-14}$ | 3.7 x $10^{-6}$ |

| Lipid class | PE acyl chain |
|---|---|
| Feature | 17:0 |
| One-way ANOVA *p-value* | 1.9 x $10^{-80}$ |
| FDR adjuested *p-value* | 8.2 x $10^{-80}$ |

| | Naive B | Memory B |
|---|---|---|
| Naive B | # | |
| Memory B | 1.0 x $10^{-1}$ | # |
| CD4 T Naive | 2.8 x $10^{-3}$ | 8.3 x $10^{-1}$ |
| CD4 T Central Memory | 8.2 x $10^{-1}$ | 8.3 x $10^{-1}$ |
| CD4 T Effector Memory | 1.0 x $10^{-1}$ | 1.0 x $10^{-1}$ |
| CD8 T Naive | 1.6 x $10^{-2}$ | 1.9 x $10^{-2}$ |
| CD8 T Central Memory | 1.0 x $10^{-1}$ | 1.0 x $10^{-1}$ |
| CD8 T Effector Memory | 1.0 x $10^{-1}$ | 1.0 x $10^{-1}$ |
| CD56 Dim NK | 7.1 x $10^{-2}$ | 8.1 x $10^{-2}$ |
| CD56 Bright NK | 1.0 x $10^{-1}$ | 1.0 x $10^{-1}$ |
| Classical Monocyte | 3.5 x $10^{-3}$ | 5.0 x $10^{-3}$ |
| Intermediate Monocyte | 9.0 x $10^{-4}$ | 1.4 x $10^{-3}$ |
| Non-classical Monocyte | 6.2 x $10^{-1}$ | 6.6 x $10^{-1}$ |
| Basophil | 3.0 x $10^{-14}$ | 5.1 x $10^{-14}$ |
| Eosinophil | < 1.0 x $10^{-16}$ | 1.5 x $10^{-14}$ |
| Neutrophil | < 1.0 x $10^{-16}$ | < 1.0 x $10^{-16}$ |

| Lipid class | PE acyl chain |
|---|---|
| Feature | 17:1 |

| | |
|---|---|
| Naive B | # |

| | |
|---|---|
| One-way ANOVA *p-value* | $1.5 \times 10^{-18}$ |
| FDR adjuested *p-value* | $1.8 \times 10^{-18}$ |

| | Naive B | Memory B |
|---|---|---|
| Memory B | $1.1 \times 10^{-2}$ | # |
| CD4 T Naive | $1.1 \times 10^{-2}$ | $1.0 \times 10^{-1}$ |
| CD4 T Central Memory | $1.2 \times 10^{-4}$ | $1.0 \times 10^{-1}$ |
| CD4 T Effector Memory | $8.5 \times 10^{-8}$ | $4.6 \times 10^{-1}$ |
| CD8 T Naive | $9.1 \times 10^{-3}$ | $1.0 \times 10^{-1}$ |
| CD8 T Central Memory | $1.8 \times 10^{-4}$ | $1.0 \times 10^{-1}$ |
| CD8 T Effector Memory | $7.2 \times 10^{-9}$ | $2.0 \times 10^{-1}$ |
| CD56 Dim NK | $4.5 \times 10^{-9}$ | $1.3 \times 10^{-1}$ |
| CD56 Bright NK | $4.0 \times 10^{-6}$ | $8.8 \times 10^{-1}$ |
| Classical Monocyte | $7.1 \times 10^{-10}$ | $8.9 \times 10^{-2}$ |
| Intermediate Monocyte | $6.5 \times 10^{-7}$ | $7.8 \times 10^{-1}$ |
| Non-classical Monocyte | $2.5 \times 10^{-7}$ | $6.1 \times 10^{-1}$ |
| Basophil | $3.1 \times 10^{-1}$ | $1.0 \times 10^{-1}$ |
| Eosinophil | $8.7 \times 10^{-14}$ | $7.7 \times 10^{-7}$ |
| Neutrophil | $3.0 \times 10^{-9}$ | $8.8 \times 10^{-2}$ |

| Lipid class | **PE acyl chain** |
|---|---|
| Feature | 18:0 |
| One-way ANOVA *p-value* | $5.2 \times 10^{-55}$ |
| FDR adjuested *p-value* | $1.1 \times 10^{-54}$ |

| | | |
|---|---|---|
| Naive B | # | |
| Memory B | $3.6 \times 10^{-7}$ | # |
| CD4 T Naive | $9.3 \times 10^{-14}$ | $1.2 \times 10^{-2}$ |
| CD4 T Central Memory | $7.3 \times 10^{-14}$ | $1.2 \times 10^{-2}$ |
| CD4 T Effector Memory | $< 1.0 \times 10^{-16}$ | $5.6 \times 10^{-6}$ |
| CD8 T Naive | $2.5 \times 10^{-13}$ | $6.0 \times 10^{-1}$ |
| CD8 T Central Memory | $8.8 \times 10^{-14}$ | $3.1 \times 10^{-2}$ |
| CD8 T Effector Memory | $< 1.0 \times 10^{-16}$ | $1.2 \times 10^{-6}$ |
| CD56 Dim NK | $< 1.0 \times 10^{-16}$ | $4.5 \times 10^{-5}$ |
| CD56 Bright NK | $2.8 \times 10^{-14}$ | $4.1 \times 10^{-4}$ |
| Classical Monocyte | $< 1.0 \times 10^{-16}$ | $9.5 \times 10^{-14}$ |
| Intermediate Monocyte | $< 1.0 \times 10^{-16}$ | $1.1 \times 10^{-8}$ |
| Non-classical Monocyte | $< 1.0 \times 10^{-16}$ | $3.2 \times 10^{-8}$ |
| Basophil | $4.2 \times 10^{-11}$ | $1.0 \times 10^{-1}$ |
| Eosinophil | $< 1.0 \times 10^{-16}$ | $< 1.0 \times 10^{-16}$ |
| Neutrophil | $< 1.0 \times 10^{-16}$ | $2.7 \times 10^{-5}$ |

| | | | |
|---|---|---|---|
| | | Naive B | Memory B |

| Lipid class | PE acyl chain |
|---|---|
| Feature | 18:1 |
| One-way ANOVA *p-value* | $1.1 \times 10^{-162}$ |
| FDR adjuested *p-value* | $1.4 \times 10^{-161}$ |

| | Naive B | Memory B |
|---|---|---|
| Naive B | # | |
| Memory B | $1.0 \times 10^{-1}$ | # |
| CD4 T Naive | $2.8 \times 10^{-5}$ | $3.6 \times 10^{-1}$ |
| CD4 T Central Memory | $3.9 \times 10^{-1}$ | $3.6 \times 10^{-1}$ |
| CD4 T Effector Memory | $1.0 \times 10^{-1}$ | $1.0 \times 10^{-1}$ |
| CD8 T Naive | $9.0 \times 10^{-5}$ | $9.0 \times 10^{-5}$ |
| CD8 T Central Memory | $1.0 \times 10^{-1}$ | $1.0 \times 10^{-1}$ |
| CD8 T Effector Memory | $1.0 \times 10^{-1}$ | $1.0 \times 10^{-1}$ |
| CD56 Dim NK | $5.3 \times 10^{-2}$ | $4.8 \times 10^{-2}$ |
| CD56 Bright NK | $7.5 \times 10^{-4}$ | $1.5 \times 10^{-3}$ |
| Classical Monocyte | $< 1.0 \times 10^{-16}$ | $< 1.0 \times 10^{-16}$ |
| Intermediate Monocyte | $< 1.0 \times 10^{-16}$ | $< 1.0 \times 10^{-16}$ |
| Non-classical Monocyte | $1.0 \times 10^{-13}$ | $1.9 \times 10^{-13}$ |
| Basophil | $< 1.0 \times 10^{-16}$ | $< 1.0 \times 10^{-16}$ |
| Eosinophil | $< 1.0 \times 10^{-16}$ | $< 1.0 \times 10^{-16}$ |
| Neutrophil | $< 1.0 \times 10^{-16}$ | $< 1.0 \times 10^{-16}$ |
| | Naive B | Memory B |

| Lipid class | PE acyl chain |
|---|---|
| Feature | 18:2 |
| One-way ANOVA *p-value* | $8.0 \times 10^{-17}$ |
| FDR adjuested *p-value* | $8.7 \times 10^{-17}$ |

| | | |
|---|---|---|
| Naive B | # | |
| Memory B | $6.3 \times 10^{-1}$ | # |
| CD4 T Naive | $1.0 \times 10^{-1}$ | $1.0 \times 10^{-1}$ |
| CD4 T Central Memory | $4.4 \times 10^{-1}$ | $1.0 \times 10^{-1}$ |
| CD4 T Effector Memory | $7.7 \times 10^{-2}$ | $1.0 \times 10^{-1}$ |
| CD8 T Naive | $2.8 \times 10^{-1}$ | $1.0 \times 10^{-1}$ |
| CD8 T Central Memory | $4.8 \times 10^{-1}$ | $1.0 \times 10^{-1}$ |
| CD8 T Effector Memory | $2.2 \times 10^{-1}$ | $1.0 \times 10^{-1}$ |

| | Naive B | Memory B |
|---|---|---|
| CD56 Dim NK | $1.0 \times 10^{-1}$ | $1.0 \times 10^{-1}$ |
| CD56 Bright NK | $1.0 \times 10^{-1}$ | $4.5 \times 10^{-1}$ |
| Classical Monocyte | $3.4 \times 10^{-1}$ | $1.0 \times 10^{-1}$ |
| Intermediate Monocyte | $6.7 \times 10^{-1}$ | $1.0 \times 10^{-1}$ |
| Non-classical Monocyte | $4.8 \times 10^{-1}$ | $1.0 \times 10^{-1}$ |
| Basophil | $1.3 \times 10^{-2}$ | $6.9 \times 10^{-7}$ |
| Eosinophil | $8.7 \times 10^{-9}$ | $1.1 \times 10^{-3}$ |
| Neutrophil | $1.0 \times 10^{-1}$ | $1.0 \times 10^{-1}$ |

| Lipid class | **PE acyl chain** |
|---|---|
| Feature | 18:3 |
| One-way ANOVA *p-value* | $1.0 \times 10^{-1}$ |
| FDR adjuested *p-value* | $1.0 \times 10^{-1}$ |

| | Naive B | Memory B |
|---|---|---|
| Naive B | # | |
| Memory B | $1.0 \times 10^{-1}$ | # |
| CD4 T Naive | $1.0 \times 10^{-1}$ | $1.0 \times 10^{-1}$ |
| CD4 T Central Memory | $1.0 \times 10^{-1}$ | $1.0 \times 10^{-1}$ |
| CD4 T Effector Memory | $1.0 \times 10^{-1}$ | $1.0 \times 10^{-1}$ |
| CD8 T Naive | $1.0 \times 10^{-1}$ | $1.0 \times 10^{-1}$ |
| CD8 T Central Memory | $1.0 \times 10^{-1}$ | $1.0 \times 10^{-1}$ |
| CD8 T Effector Memory | $1.0 \times 10^{-1}$ | $1.0 \times 10^{-1}$ |
| CD56 Dim NK | $1.0 \times 10^{-1}$ | $1.0 \times 10^{-1}$ |
| CD56 Bright NK | $1.0 \times 10^{-1}$ | $1.0 \times 10^{-1}$ |
| Classical Monocyte | $1.0 \times 10^{-1}$ | $1.0 \times 10^{-1}$ |
| Intermediate Monocyte | $1.0 \times 10^{-1}$ | $1.0 \times 10^{-1}$ |
| Non-classical Monocyte | $1.0 \times 10^{-1}$ | $1.0 \times 10^{-1}$ |
| Basophil | $1.0 \times 10^{-1}$ | $1.0 \times 10^{-1}$ |
| Eosinophil | $1.0 \times 10^{-1}$ | $1.0 \times 10^{-1}$ |
| Neutrophil | $1.0 \times 10^{-1}$ | $1.0 \times 10^{-1}$ |

Lipid class **PE acyl chain**

| Feature | 19:0 |
|---|---|
| One-way ANOVA *p-value* | $1.0 \times 10^{-1}$ |
| FDR adjuested *p-value* | $1.0 \times 10^{-1}$ |

| | Naive B | Memory B |
|---|---|---|
| Naive B | # | |
| Memory B | $1.0 \times 10^{-1}$ | # |
| CD4 T Naive | $1.0 \times 10^{-1}$ | $1.0 \times 10^{-1}$ |
| CD4 T Central Memory | $1.0 \times 10^{-1}$ | $1.0 \times 10^{-1}$ |
| CD4 T Effector Memory | $1.0 \times 10^{-1}$ | $1.0 \times 10^{-1}$ |
| CD8 T Naive | $1.0 \times 10^{-1}$ | $1.0 \times 10^{-1}$ |
| CD8 T Central Memory | $1.0 \times 10^{-1}$ | $1.0 \times 10^{-1}$ |
| CD8 T Effector Memory | $1.0 \times 10^{-1}$ | $1.0 \times 10^{-1}$ |
| CD56 Dim NK | $1.0 \times 10^{-1}$ | $1.0 \times 10^{-1}$ |
| CD56 Bright NK | $1.0 \times 10^{-1}$ | $1.0 \times 10^{-1}$ |
| Classical Monocyte | $1.0 \times 10^{-1}$ | $1.0 \times 10^{-1}$ |
| Intermediate Monocyte | $1.0 \times 10^{-1}$ | $1.0 \times 10^{-1}$ |
| Non-classical Monocyte | $1.0 \times 10^{-1}$ | $1.0 \times 10^{-1}$ |
| Basophil | $1.0 \times 10^{-1}$ | $1.0 \times 10^{-1}$ |
| Eosinophil | $1.0 \times 10^{-1}$ | $1.0 \times 10^{-1}$ |
| Neutrophil | $1.0 \times 10^{-1}$ | $1.0 \times 10^{-1}$ |

| Lipid class | **PE acyl chain** |
|---|---|
| Feature | 20:0 |
| One-way ANOVA *p-value* | $1.0 \times 10^{-1}$ |
| FDR adjuested *p-value* | $1.0 \times 10^{-1}$ |

| | | |
|---|---|---|
| Naive B | # | |
| Memory B | $1.0 \times 10^{-1}$ | # |
| CD4 T Naive | $1.0 \times 10^{-1}$ | $1.0 \times 10^{-1}$ |
| CD4 T Central Memory | $1.0 \times 10^{-1}$ | $1.0 \times 10^{-1}$ |
| CD4 T Effector Memory | $1.0 \times 10^{-1}$ | $1.0 \times 10^{-1}$ |
| CD8 T Naive | $1.0 \times 10^{-1}$ | $1.0 \times 10^{-1}$ |
| CD8 T Central Memory | $1.0 \times 10^{-1}$ | $1.0 \times 10^{-1}$ |
| CD8 T Effector Memory | $1.0 \times 10^{-1}$ | $1.0 \times 10^{-1}$ |
| CD56 Dim NK | $1.0 \times 10^{-1}$ | $1.0 \times 10^{-1}$ |
| CD56 Bright NK | $1.0 \times 10^{-1}$ | $1.0 \times 10^{-1}$ |
| Classical Monocyte | $1.0 \times 10^{-1}$ | $1.0 \times 10^{-1}$ |
| Intermediate Monocyte | $1.0 \times 10^{-1}$ | $1.0 \times 10^{-1}$ |
| Non-classical Monocyte | $1.0 \times 10^{-1}$ | $1.0 \times 10^{-1}$ |
| Basophil | $1.0 \times 10^{-1}$ | $1.0 \times 10^{-1}$ |
| Eosinophil | $1.0 \times 10^{-1}$ | $1.0 \times 10^{-1}$ |
| Neutrophil | $1.0 \times 10^{-1}$ | $1.0 \times 10^{-1}$ |

Naive B

Memory B

| | |
|---|---|
| Lipid class | **PE acyl chain** |
| Feature | 20:1 |
| One-way ANOVA *p-value* | $1.0 \times 10^{-1}$ |
| FDR adjuested *p-value* | $1.0 \times 10^{-1}$ |

| | Naive B | Memory B |
|---|---|---|
| Naive B | # | |
| Memory B | $1.0 \times 10^{-1}$ | # |
| CD4 T Naive | $1.0 \times 10^{-1}$ | $1.0 \times 10^{-1}$ |
| CD4 T Central Memory | $1.0 \times 10^{-1}$ | $1.0 \times 10^{-1}$ |
| CD4 T Effector Memory | $1.0 \times 10^{-1}$ | $1.0 \times 10^{-1}$ |
| CD8 T Naive | $1.0 \times 10^{-1}$ | $1.0 \times 10^{-1}$ |
| CD8 T Central Memory | $1.0 \times 10^{-1}$ | $1.0 \times 10^{-1}$ |
| CD8 T Effector Memory | $1.0 \times 10^{-1}$ | $1.0 \times 10^{-1}$ |
| CD56 Dim NK | $1.0 \times 10^{-1}$ | $1.0 \times 10^{-1}$ |
| CD56 Bright NK | $1.0 \times 10^{-1}$ | $1.0 \times 10^{-1}$ |
| Classical Monocyte | $1.0 \times 10^{-1}$ | $1.0 \times 10^{-1}$ |
| Intermediate Monocyte | $1.0 \times 10^{-1}$ | $1.0 \times 10^{-1}$ |
| Non-classical Monocyte | $1.0 \times 10^{-1}$ | $1.0 \times 10^{-1}$ |
| Basophil | $1.0 \times 10^{-1}$ | $1.0 \times 10^{-1}$ |
| Eosinophil | $1.0 \times 10^{-1}$ | $1.0 \times 10^{-1}$ |
| Neutrophil | $1.0 \times 10^{-1}$ | $1.0 \times 10^{-1}$ |

| | |
|---|---|
| Lipid class | **PE acyl chain** |
| Feature | 20:2 |
| One-way ANOVA *p-value* | $2.8 \times 10^{-42}$ |
| FDR adjuested *p-value* | $5.1 \times 10^{-42}$ |

| | Naive B | Memory B |
|---|---|---|
| Naive B | # | |
| Memory B | $1.0 \times 10^{-1}$ | # |
| CD4 T Naive | $1.1 \times 10^{-5}$ | $2.0 \times 10^{-1}$ |
| CD4 T Central Memory | $3.0 \times 10^{-1}$ | $2.0 \times 10^{-1}$ |
| CD4 T Effector Memory | $1.0 \times 10^{-1}$ | $1.0 \times 10^{-1}$ |
| CD8 T Naive | $6.0 \times 10^{-14}$ | $6.0 \times 10^{-14}$ |
| CD8 T Central Memory | $2.4 \times 10^{-2}$ | $1.3 \times 10^{-2}$ |
| CD8 T Effector Memory | $9.8 \times 10^{-2}$ | $5.9 \times 10^{-2}$ |

| | Naive B | Memory B |
|---|---|---|
| CD56 Dim NK | 1.1 x $10^{-6}$ | 5.3 x $10^{-7}$ |
| CD56 Bright NK | 6.1 x $10^{-5}$ | 3.1 x $10^{-5}$ |
| Classical Monocyte | 8.1 x $10^{-3}$ | 4.3 x $10^{-3}$ |
| Intermediate Monocyte | 1.9 x $10^{-1}$ | 1.2 x $10^{-1}$ |
| Non-classical Monocyte | 1.0 x $10^{-1}$ | 9.3 x $10^{-1}$ |
| Basophil | 3.4 x $10^{-14}$ | 2.7 x $10^{-14}$ |
| Eosinophil | 1.0 x $10^{-1}$ | 1.0 x $10^{-1}$ |
| Neutrophil | 4.5 x $10^{-3}$ | 1.4 x $10^{-2}$ |

| Lipid class | PE acyl chain |
|---|---|
| Feature | 20:3 |
| One-way ANOVA *p-value* | 1.0 x $10^{-1}$ |
| FDR adjuested *p-value* | 1.0 x $10^{-1}$ |

| | Naive B | Memory B |
|---|---|---|
| Naive B | # | |
| Memory B | 1.0 x $10^{-1}$ | # |
| CD4 T Naive | 1.0 x $10^{-1}$ | 1.0 x $10^{-1}$ |
| CD4 T Central Memory | 1.0 x $10^{-1}$ | 1.0 x $10^{-1}$ |
| CD4 T Effector Memory | 1.0 x $10^{-1}$ | 1.0 x $10^{-1}$ |
| CD8 T Naive | 1.0 x $10^{-1}$ | 1.0 x $10^{-1}$ |
| CD8 T Central Memory | 1.0 x $10^{-1}$ | 1.0 x $10^{-1}$ |
| CD8 T Effector Memory | 1.0 x $10^{-1}$ | 1.0 x $10^{-1}$ |
| CD56 Dim NK | 1.0 x $10^{-1}$ | 1.0 x $10^{-1}$ |
| CD56 Bright NK | 1.0 x $10^{-1}$ | 1.0 x $10^{-1}$ |
| Classical Monocyte | 1.0 x $10^{-1}$ | 1.0 x $10^{-1}$ |
| Intermediate Monocyte | 1.0 x $10^{-1}$ | 1.0 x $10^{-1}$ |
| Non-classical Monocyte | 1.0 x $10^{-1}$ | 1.0 x $10^{-1}$ |
| Basophil | 1.0 x $10^{-1}$ | 1.0 x $10^{-1}$ |
| Eosinophil | 1.0 x $10^{-1}$ | 1.0 x $10^{-1}$ |
| Neutrophil | 1.0 x $10^{-1}$ | 1.0 x $10^{-1}$ |

Lipid class **PE acyl chain**

| Feature | 20:4 |
|---|---|
| One-way ANOVA *p-value* | $5.0 \times 10^{-13}$ |
| FDR adjuested *p-value* | $3.2 \times 10^{-12}$ |

| | Naive B | Memory B |
|---|---|---|
| Naive B | # | |
| Memory B | $7.7 \times 10^{-1}$ | # |
| CD4 T Naive | $1.2 \times 10^{-2}$ | $1.0 \times 10^{-1}$ |
| CD4 T Central Memory | $5.3 \times 10^{-1}$ | $1.0 \times 10^{-1}$ |
| CD4 T Effector Memory | $1.0 \times 10^{-1}$ | $8.9 \times 10^{-1}$ |
| CD8 T Naive | $1.1 \times 10^{-3}$ | $5.9 \times 10^{-1}$ |
| CD8 T Central Memory | $1.0 \times 10^{-1}$ | $1.0 \times 10^{-1}$ |
| CD8 T Effector Memory | $1.0 \times 10^{-1}$ | $7.3 \times 10^{-1}$ |
| CD56 Dim NK | $1.0 \times 10^{-1}$ | $6.1 \times 10^{-2}$ |
| CD56 Bright NK | $2.5 \times 10^{-8}$ | $4.1 \times 10^{-13}$ |
| Classical Monocyte | $1.2 \times 10^{-10}$ | $8.4 \times 10^{-14}$ |
| Intermediate Monocyte | $1.1 \times 10^{-10}$ | $8.8 \times 10^{-14}$ |
| Non-classical Monocyte | $1.1 \times 10^{-4}$ | $4.5 \times 10^{-9}$ |
| Basophil | $< 1.0 \times 10^{-16}$ | $< 1.0 \times 10^{-16}$ |
| Eosinophil | $< 1.0 \times 10^{-16}$ | $< 1.0 \times 10^{-16}$ |
| Neutrophil | $< 1.0 \times 10^{-16}$ | $< 1.0 \times 10^{-16}$ |

| Lipid class | **PE acyl chain** |
|---|---|
| Feature | 20:5 |
| One-way ANOVA *p-value* | $2.4 \times 10^{-8}$ |
| FDR adjuested *p-value* | $2.4 \times 10^{-8}$ |

| | | |
|---|---|---|
| Naive B | # | |
| Memory B | $2.7 \times 10^{-1}$ | # |
| CD4 T Naive | $9.6 \times 10^{-2}$ | $1.0 \times 10^{-1}$ |
| CD4 T Central Memory | $1.8 \times 10^{-2}$ | $1.0 \times 10^{-1}$ |
| CD4 T Effector Memory | $4.0 \times 10^{-3}$ | $1.0 \times 10^{-1}$ |
| CD8 T Naive | $4.3 \times 10^{-2}$ | $1.0 \times 10^{-1}$ |
| CD8 T Central Memory | $4.4 \times 10^{-3}$ | $1.0 \times 10^{-1}$ |
| CD8 T Effector Memory | $1.0 \times 10^{-2}$ | $1.0 \times 10^{-1}$ |
| CD56 Dim NK | $4.6 \times 10^{-2}$ | $1.0 \times 10^{-1}$ |
| CD56 Bright NK | $3.2 \times 10^{-4}$ | $8.1 \times 10^{-1}$ |
| Classical Monocyte | $8.6 \times 10^{-6}$ | $3.5 \times 10^{-1}$ |
| Intermediate Monocyte | $6.1 \times 10^{-6}$ | $3.1 \times 10^{-1}$ |
| Non-classical Monocyte | $4.1 \times 10^{-6}$ | $2.3 \times 10^{-1}$ |
| Basophil | $1.0 \times 10^{-2}$ | $1.0 \times 10^{-1}$ |
| Eosinophil | $8.5 \times 10^{-8}$ | $3.3 \times 10^{-2}$ |
| Neutrophil | $6.5 \times 10^{-7}$ | $6.9 \times 10^{-2}$ |

| | Naive B | Memory B |
|---|---|---|

| | |
|---|---|
| Lipid class | **PE acyl chain** |
| Feature | 22:2 |
| One-way ANOVA *p-value* | $1.9 \times 10^{-38}$ |
| FDR adjuested *p-value* | $3.0 \times 10^{-38}$ |

| | Naive B | Memory B |
|---|---|---|
| Naive B | # | |
| Memory B | $3.9 \times 10^{-2}$ | # |
| CD4 T Naive | $1.0 \times 10^{-1}$ | $5.5 \times 10^{-1}$ |
| CD4 T Central Memory | $1.0 \times 10^{-1}$ | $5.5 \times 10^{-1}$ |
| CD4 T Effector Memory | $8.6 \times 10^{-1}$ | $1.0 \times 10^{-1}$ |
| CD8 T Naive | $1.0 \times 10^{-1}$ | $3.9 \times 10^{-1}$ |
| CD8 T Central Memory | $1.0 \times 10^{-1}$ | $1.5 \times 10^{-1}$ |
| CD8 T Effector Memory | $8.7 \times 10^{-1}$ | $1.0 \times 10^{-1}$ |
| CD56 Dim NK | $4.5 \times 10^{-3}$ | $2.4 \times 10^{-10}$ |
| CD56 Bright NK | $6.2 \times 10^{-4}$ | $1.3 \times 10^{-11}$ |
| Classical Monocyte | $4.3 \times 10^{-8}$ | $2.1 \times 10^{-1}$ |
| Intermediate Monocyte | $2.7 \times 10^{-7}$ | $4.0 \times 10^{-1}$ |
| Non-classical Monocyte | $2.0 \times 10^{-7}$ | $3.2 \times 10^{-1}$ |
| Basophil | $7.8 \times 10^{-1}$ | $1.0 \times 10^{-1}$ |
| Eosinophil | $3.1 \times 10^{-2}$ | $1.0 \times 10^{-1}$ |
| Neutrophil | $9.7 \times 10^{-14}$ | $3.9 \times 10^{-6}$ |
| | Naive B | Memory B |

| | |
|---|---|
| Lipid class | **PE acyl chain** |
| Feature | 22:4 |
| One-way ANOVA *p-value* | $1.6 \times 10^{-68}$ |
| FDR adjuested *p-value* | $5.3 \times 10^{-68}$ |

| | | |
|---|---|---|
| Naive B | # | |
| Memory B | $1.0 \times 10^{-1}$ | # |
| CD4 T Naive | $1.0 \times 10^{-1}$ | $9.3 \times 10^{-1}$ |
| CD4 T Central Memory | $3.6 \times 10^{-1}$ | $9.3 \times 10^{-1}$ |
| CD4 T Effector Memory | $3.1 \times 10^{-12}$ | $1.2 \times 10^{-9}$ |
| CD8 T Naive | $1.0 \times 10^{-1}$ | $1.0 \times 10^{-1}$ |
| CD8 T Central Memory | $2.7 \times 10^{-1}$ | $8.5 \times 10^{-1}$ |
| CD8 T Effector Memory | $9.3 \times 10^{-14}$ | $2.4 \times 10^{-13}$ |

| | Naive B | Memory B |
|---|---|---|
| CD56 Dim NK | < 1.0 x $10^{-16}$ | < 1.0 x $10^{-16}$ |
| CD56 Bright NK | < 1.0 x $10^{-16}$ | < 1.0 x $10^{-16}$ |
| Classical Monocyte | 6.6 x $10^{-6}$ | 5.8 x $10^{-4}$ |
| Intermediate Monocyte | 1.1 x $10^{-8}$ | 2.3 x $10^{-6}$ |
| Non-classical Monocyte | 9.2 x $10^{-14}$ | 1.2 x $10^{-12}$ |
| Basophil | 1.0 x $10^{-1}$ | 1.0 x $10^{-1}$ |
| Eosinophil | < 1.0 x $10^{-16}$ | < 1.0 x $10^{-16}$ |
| Neutrophil | 1.0 x $10^{-1}$ | 1.0 x $10^{-1}$ |

| Lipid class | **PE acyl chain** |
|---|---|
| Feature | 22:5 |
| One-way ANOVA *p-value* | 1.0 x $10^{-1}$ |
| FDR adjuested *p-value* | 1.0 x $10^{-1}$ |

| | Naive B | Memory B |
|---|---|---|
| Naive B | # | |
| Memory B | 1.0 x $10^{-1}$ | # |
| CD4 T Naive | 1.0 x $10^{-1}$ | 1.0 x $10^{-1}$ |
| CD4 T Central Memory | 1.0 x $10^{-1}$ | 1.0 x $10^{-1}$ |
| CD4 T Effector Memory | 1.0 x $10^{-1}$ | 1.0 x $10^{-1}$ |
| CD8 T Naive | 1.0 x $10^{-1}$ | 1.0 x $10^{-1}$ |
| CD8 T Central Memory | 1.0 x $10^{-1}$ | 1.0 x $10^{-1}$ |
| CD8 T Effector Memory | 1.0 x $10^{-1}$ | 1.0 x $10^{-1}$ |
| CD56 Dim NK | 1.0 x $10^{-1}$ | 1.0 x $10^{-1}$ |
| CD56 Bright NK | 1.0 x $10^{-1}$ | 1.0 x $10^{-1}$ |
| Classical Monocyte | 1.0 x $10^{-1}$ | 1.0 x $10^{-1}$ |
| Intermediate Monocyte | 1.0 x $10^{-1}$ | 1.0 x $10^{-1}$ |
| Non-classical Monocyte | 1.0 x $10^{-1}$ | 1.0 x $10^{-1}$ |
| Basophil | 1.0 x $10^{-1}$ | 1.0 x $10^{-1}$ |
| Eosinophil | 1.0 x $10^{-1}$ | 1.0 x $10^{-1}$ |
| Neutrophil | 1.0 x $10^{-1}$ | 1.0 x $10^{-1}$ |

Lipid class **PE acyl chain**

| Feature | 22:6 |
|---|---|
| One-way ANOVA *p-value* | $7.3 \times 10^{-27}$ |
| FDR adjuested *p-value* | $9.5 \times 10^{-27}$ |

| | Naive B | Memory B |
|---|---|---|
| Naive B | # | |
| Memory B | $1.0 \times 10^{-1}$ | # |
| CD4 T Naive | $1.0 \times 10^{-1}$ | $1.0 \times 10^{-1}$ |
| CD4 T Central Memory | $1.0 \times 10^{-1}$ | $1.0 \times 10^{-1}$ |
| CD4 T Effector Memory | $4.5 \times 10^{-1}$ | $1.0 \times 10^{-1}$ |
| CD8 T Naive | $1.0 \times 10^{-1}$ | $1.0 \times 10^{-1}$ |
| CD8 T Central Memory | $1.0 \times 10^{-1}$ | $1.0 \times 10^{-1}$ |
| CD8 T Effector Memory | $8.9 \times 10^{-1}$ | $1.0 \times 10^{-1}$ |
| CD56 Dim NK | $1.0 \times 10^{-1}$ | $1.0 \times 10^{-1}$ |
| CD56 Bright NK | $1.0 \times 10^{-1}$ | $1.0 \times 10^{-1}$ |
| Classical Monocyte | $4.0 \times 10^{-3}$ | $9.0 \times 10^{-2}$ |
| Intermediate Monocyte | $2.8 \times 10^{-2}$ | $3.2 \times 10^{-1}$ |
| Non-classical Monocyte | $2.7 \times 10^{-4}$ | $1.1 \times 10^{-2}$ |
| Basophil | $3.3 \times 10^{-5}$ | $7.4 \times 10^{-7}$ |
| Eosinophil | $2.7 \times 10^{-1}$ | $3.3 \times 10^{-2}$ |
| Neutrophil | $1.1 \times 10^{-4}$ | $3.4 \times 10^{-6}$ |

**One-way ANOVA results of <u>diacyl PE acyl chain</u> composition in 16 different hum**

| CD4 T Naive | CD4 T Central Memory | CD4 T Effector Memory | CD8 T Naive | CD8 T Central Memory | CD8 T Effector Memory | CD56 Dim NK |
|---|---|---|---|---|---|---|
| # | | | | | | |
| 1.0 x $10^{-1}$ | # | | | | | |
| 1.0 x $10^{-1}$ | 1.0 x $10^{-1}$ | # | | | | |
| 1.0 x $10^{-1}$ | 1.0 x $10^{-1}$ | 1.0 x $10^{-1}$ | # | | | |
| 1.0 x $10^{-1}$ | 1.0 x $10^{-1}$ | 1.0 x $10^{-1}$ | 1.0 x $10^{-1}$ | # | | |
| 1.0 x $10^{-1}$ | 1.0 x $10^{-1}$ | 1.0 x $10^{-1}$ | 1.0 x $10^{-1}$ | 1.0 x $10^{-1}$ | # | |
| 1.0 x $10^{-1}$ | 1.0 x $10^{-1}$ | 1.0 x $10^{-1}$ | 1.0 x $10^{-1}$ | 1.0 x $10^{-1}$ | 1.0 x $10^{-1}$ | # |
| 1.0 x $10^{-1}$ | 1.0 x $10^{-1}$ | 1.0 x $10^{-1}$ | 1.0 x $10^{-1}$ | 1.0 x $10^{-1}$ | 1.0 x $10^{-1}$ | 1.0 x $10^{-1}$ |
| 1.0 x $10^{-1}$ | 1.0 x $10^{-1}$ | 1.0 x $10^{-1}$ | 1.0 x $10^{-1}$ | 1.0 x $10^{-1}$ | 1.0 x $10^{-1}$ | 1.0 x $10^{-1}$ |
| 1.0 x $10^{-1}$ | 1.0 x $10^{-1}$ | 1.0 x $10^{-1}$ | 1.0 x $10^{-1}$ | 1.0 x $10^{-1}$ | 1.0 x $10^{-1}$ | 1.0 x $10^{-1}$ |
| 1.0 x $10^{-1}$ | 1.0 x $10^{-1}$ | 1.0 x $10^{-1}$ | 1.0 x $10^{-1}$ | 1.0 x $10^{-1}$ | 1.0 x $10^{-1}$ | 1.0 x $10^{-1}$ |
| 1.0 x $10^{-1}$ | 1.0 x $10^{-1}$ | 1.0 x $10^{-1}$ | 1.0 x $10^{-1}$ | 1.0 x $10^{-1}$ | 1.0 x $10^{-1}$ | 1.0 x $10^{-1}$ |
| 1.0 x $10^{-1}$ | 1.0 x $10^{-1}$ | 1.0 x $10^{-1}$ | 1.0 x $10^{-1}$ | 1.0 x $10^{-1}$ | 1.0 x $10^{-1}$ | 1.0 x $10^{-1}$ |
| 1.0 x $10^{-1}$ | 1.0 x $10^{-1}$ | 1.0 x $10^{-1}$ | 1.0 x $10^{-1}$ | 1.0 x $10^{-1}$ | 1.0 x $10^{-1}$ | 1.0 x $10^{-1}$ |

| | | | | | | |
|---|---|---|---|---|---|---|
| # | | | | | | |
| 1.0 x $10^{-1}$ | # | | | | | |
| 1.0 x $10^{-1}$ | 1.0 x $10^{-1}$ | # | | | | |
| 1.0 x $10^{-1}$ | 1.0 x $10^{-1}$ | 1.0 x $10^{-1}$ | # | | | |
| 1.0 x $10^{-1}$ | 1.0 x $10^{-1}$ | 1.0 x $10^{-1}$ | 1.0 x $10^{-1}$ | # | | |
| 1.0 x $10^{-1}$ | 1.0 x $10^{-1}$ | 1.0 x $10^{-1}$ | 1.0 x $10^{-1}$ | 1.0 x $10^{-1}$ | # | |
| 1.0 x $10^{-1}$ | 1.0 x $10^{-1}$ | 1.0 x $10^{-1}$ | 1.0 x $10^{-1}$ | 1.0 x $10^{-1}$ | 1.0 x $10^{-1}$ | # |
| 1.0 x $10^{-1}$ | 1.0 x $10^{-1}$ | 1.0 x $10^{-1}$ | 1.0 x $10^{-1}$ | 1.0 x $10^{-1}$ | 1.0 x $10^{-1}$ | 1.0 x $10^{-1}$ |
| 1.0 x $10^{-1}$ | 1.0 x $10^{-1}$ | 1.0 x $10^{-1}$ | 1.0 x $10^{-1}$ | 1.0 x $10^{-1}$ | 1.0 x $10^{-1}$ | 1.0 x $10^{-1}$ |
| 1.0 x $10^{-1}$ | 1.0 x $10^{-1}$ | 1.0 x $10^{-1}$ | 1.0 x $10^{-1}$ | 1.0 x $10^{-1}$ | 1.0 x $10^{-1}$ | 1.0 x $10^{-1}$ |
| 1.0 x $10^{-1}$ | 1.0 x $10^{-1}$ | 1.0 x $10^{-1}$ | 1.0 x $10^{-1}$ | 1.0 x $10^{-1}$ | 1.0 x $10^{-1}$ | 1.0 x $10^{-1}$ |
| 1.0 x $10^{-1}$ | 1.0 x $10^{-1}$ | 1.0 x $10^{-1}$ | 1.0 x $10^{-1}$ | 1.0 x $10^{-1}$ | 1.0 x $10^{-1}$ | 1.0 x $10^{-1}$ |
| 1.0 x $10^{-1}$ | 1.0 x $10^{-1}$ | 1.0 x $10^{-1}$ | 1.0 x $10^{-1}$ | 1.0 x $10^{-1}$ | 1.0 x $10^{-1}$ | 1.0 x $10^{-1}$ |

| CD4 T Naive | CD4 T Central Memory | CD4 T Effector Memory | CD8 T Naive | CD8 T Central Memory | CD8 T Effector Memory | CD56 Dim NK |
|---|---|---|---|---|---|---|
| $1.0 \times 10^{-1}$ | $1.0 \times 10^{-1}$ | $1.0 \times 10^{-1}$ | $1.0 \times 10^{-1}$ | $1.0 \times 10^{-1}$ | $1.0 \times 10^{-1}$ | $1.0 \times 10^{-1}$ |

| CD4 T Naive | CD4 T Central Memory | CD4 T Effector Memory | CD8 T Naive | CD8 T Central Memory | CD8 T Effector Memory | CD56 Dim NK |
|---|---|---|---|---|---|---|
| # | | | | | | |
| $1.0 \times 10^{-1}$ | # | | | | | |
| $9.7 \times 10^{-2}$ | $7.0 \times 10^{-1}$ | # | | | | |
| $1.0 \times 10^{-1}$ | $8.2 \times 10^{-1}$ | $5.0 \times 10^{-3}$ | # | | | |
| $1.0 \times 10^{-1}$ | $1.0 \times 10^{-1}$ | $1.0 \times 10^{-1}$ | $4.9 \times 10^{-1}$ | # | | |
| $4.4 \times 10^{-2}$ | $5.0 \times 10^{-1}$ | $1.0 \times 10^{-1}$ | $1.7 \times 10^{-3}$ | $9.1 \times 10^{-1}$ | # | |
| $3.4 \times 10^{-6}$ | $3.5 \times 10^{-4}$ | $4.0 \times 10^{-1}$ | $2.4 \times 10^{-8}$ | $6.5 \times 10^{-3}$ | $5.9 \times 10^{-1}$ | # |
| $2.3 \times 10^{-7}$ | $3.1 \times 10^{-5}$ | $1.3 \times 10^{-1}$ | $1.2 \times 10^{-9}$ | $8.6 \times 10^{-4}$ | $2.4 \times 10^{-1}$ | $1.0 \times 10^{-1}$ |
| $3.8 \times 10^{-8}$ | $7.4 \times 10^{-6}$ | $7.5 \times 10^{-2}$ | $1.3 \times 10^{-10}$ | $2.9 \times 10^{-4}$ | $1.6 \times 10^{-1}$ | $1.0 \times 10^{-1}$ |
| $1.1 \times 10^{-2}$ | $2.4 \times 10^{-1}$ | $1.0 \times 10^{-1}$ | $2.7 \times 10^{-4}$ | $7.1 \times 10^{-1}$ | $1.0 \times 10^{-1}$ | $7.9 \times 10^{-1}$ |
| $3.8 \times 10^{-1}$ | $1.0 \times 10^{-1}$ | $1.0 \times 10^{-1}$ | $4.2 \times 10^{-2}$ | $1.0 \times 10^{-1}$ | $1.0 \times 10^{-1}$ | $1.1 \times 10^{-1}$ |
| $1.3 \times 10^{-7}$ | $2.3 \times 10^{-5}$ | $1.4 \times 10^{-1}$ | $5.0 \times 10^{-10}$ | $7.5 \times 10^{-4}$ | $2.6 \times 10^{-1}$ | $1.0 \times 10^{-1}$ |
| $< 1.0 \times 10^{-16}$ | $< 1.0 \times 10^{-16}$ | $< 1.0 \times 10^{-16}$ | $< 1.0 \times 10^{-16}$ | $< 1.0 \times 10^{-16}$ | $< 1.0 \times 10^{-16}$ | $1.0 \times 10^{-13}$ |
| $1.9 \times 10^{-8}$ | $3.0 \times 10^{-6}$ | $2.8 \times 10^{-2}$ | $9.2 \times 10^{-11}$ | $1.0 \times 10^{-4}$ | $6.3 \times 10^{-2}$ | $1.0 \times 10^{-1}$ |

| | | | | | | |
|---|---|---|---|---|---|---|
| # | | | | | | |
| $1.0 \times 10^{-1}$ | # | | | | | |
| $1.0 \times 10^{-1}$ | $1.0 \times 10^{-1}$ | # | | | | |
| $1.0 \times 10^{-1}$ | $1.0 \times 10^{-1}$ | $1.0 \times 10^{-1}$ | # | | | |
| $1.0 \times 10^{-1}$ | $1.0 \times 10^{-1}$ | $1.0 \times 10^{-1}$ | $1.0 \times 10^{-1}$ | # | | |
| $1.0 \times 10^{-1}$ | $1.0 \times 10^{-1}$ | $1.0 \times 10^{-1}$ | $1.0 \times 10^{-1}$ | $1.0 \times 10^{-1}$ | # | |
| $4.7 \times 10^{-4}$ | $2.5 \times 10^{-5}$ | $9.6 \times 10^{-7}$ | $6.8 \times 10^{-5}$ | $1.7 \times 10^{-3}$ | $1.2 \times 10^{-6}$ | # |

| CD4 T Naive | CD4 T Central Memory | CD4 T Effector Memory | CD8 T Naive | CD8 T Central Memory | CD8 T Effector Memory | CD56 Dim NK |
|---|---|---|---|---|---|---|
| $6.3 \times 10^{-5}$ | $2.7 \times 10^{-6}$ | $8.8 \times 10^{-8}$ | $7.6 \times 10^{-6}$ | $2.7 \times 10^{-4}$ | $1.2 \times 10^{-7}$ | $1.0 \times 10^{-1}$ |
| $2.1 \times 10^{-6}$ | $2.2 \times 10^{-5}$ | $1.1 \times 10^{-3}$ | $7.5 \times 10^{-6}$ | $1.2 \times 10^{-6}$ | $9.0 \times 10^{-4}$ | $6.1 \times 10^{-14}$ |
| $1.3 \times 10^{-5}$ | $1.2 \times 10^{-4}$ | $4.8 \times 10^{-3}$ | $4.6 \times 10^{-5}$ | $7.5 \times 10^{-6}$ | $3.9 \times 10^{-3}$ | $8.5 \times 10^{-14}$ |
| $8.8 \times 10^{-6}$ | $8.1 \times 10^{-5}$ | $3.1 \times 10^{-3}$ | $3.0 \times 10^{-5}$ | $5.0 \times 10^{-6}$ | $2.6 \times 10^{-3}$ | $8.4 \times 10^{-14}$ |
| $1.0 \times 10^{-1}$ | $1.0 \times 10^{-1}$ | $1.0 \times 10^{-1}$ | $1.0 \times 10^{-1}$ | $1.0 \times 10^{-1}$ | $1.0 \times 10^{-1}$ | $4.3 \times 10^{-7}$ |
| $1.7 \times 10^{-1}$ | $4.8 \times 10^{-1}$ | $9.5 \times 10^{-1}$ | $3.3 \times 10^{-1}$ | $1.1 \times 10^{-1}$ | $9.3 \times 10^{-1}$ | $8.9 \times 10^{-11}$ |
| $7.7 \times 10^{-13}$ | $8.4 \times 10^{-12}$ | $1.4 \times 10^{-9}$ | $2.4 \times 10^{-12}$ | $5.6 \times 10^{-13}$ | $1.1 \times 10^{-9}$ | $< 1.0 \times 10^{-16}$ |

| CD4 T Naive | CD4 T Central Memory | CD4 T Effector Memory | CD8 T Naive | CD8 T Central Memory | CD8 T Effector Memory | CD56 Dim NK |
|---|---|---|---|---|---|---|
| # | | | | | | |
| $6.0 \times 10^{-1}$ | # | | | | | |
| $2.2 \times 10^{-1}$ | $1.0 \times 10^{-1}$ | # | | | | |
| $1.0 \times 10^{-1}$ | $9.1 \times 10^{-1}$ | $5.4 \times 10^{-1}$ | # | | | |
| $4.4 \times 10^{-2}$ | $1.0 \times 10^{-1}$ | $1.0 \times 10^{-1}$ | $1.6 \times 10^{-1}$ | # | | |
| $1.6 \times 10^{-1}$ | $1.0 \times 10^{-1}$ | $1.0 \times 10^{-1}$ | $4.2 \times 10^{-1}$ | $1.0 \times 10^{-1}$ | # | |
| $1.0 \times 10^{-1}$ | $1.0 \times 10^{-1}$ | $8.2 \times 10^{-1}$ | $1.0 \times 10^{-1}$ | $3.9 \times 10^{-1}$ | $7.2 \times 10^{-1}$ | # |
| $2.3 \times 10^{-2}$ | $1.0 \times 10^{-1}$ | $1.0 \times 10^{-1}$ | $9.3 \times 10^{-2}$ | $1.0 \times 10^{-1}$ | $1.0 \times 10^{-1}$ | $2.6 \times 10^{-1}$ |
| $8.9 \times 10^{-13}$ | $3.1 \times 10^{-7}$ | $1.4 \times 10^{-5}$ | $9.1 \times 10^{-12}$ | $5.3 \times 10^{-4}$ | $3.0 \times 10^{-5}$ | $4.1 \times 10^{-10}$ |
| $1.9 \times 10^{-13}$ | $5.0 \times 10^{-8}$ | $2.7 \times 10^{-6}$ | $1.2 \times 10^{-12}$ | $1.3 \times 10^{-4}$ | $6.0 \times 10^{-6}$ | $6.1 \times 10^{-11}$ |
| $8.8 \times 10^{-8}$ | $3.1 \times 10^{-3}$ | $3.6 \times 10^{-2}$ | $8.3 \times 10^{-7}$ | $2.6 \times 10^{-1}$ | $5.8 \times 10^{-2}$ | $1.3 \times 10^{-5}$ |
| $< 1.0 \times 10^{-16}$ | $< 1.0 \times 10^{-16}$ | $< 1.0 \times 10^{-16}$ | $< 1.0 \times 10^{-16}$ | $1.1 \times 10^{-14}$ | $< 1.0 \times 10^{-16}$ | $< 1.0 \times 10^{-16}$ |
| $< 1.0 \times 10^{-16}$ | $< 1.0 \times 10^{-16}$ | $< 1.0 \times 10^{-16}$ | $< 1.0 \times 10^{-16}$ | $< 1.0 \times 10^{-16}$ | $< 1.0 \times 10^{-16}$ | $< 1.0 \times 10^{-16}$ |
| $< 1.0 \times 10^{-16}$ | $< 1.0 \times 10^{-16}$ | $< 1.0 \times 10^{-16}$ | $< 1.0 \times 10^{-16}$ | $< 1.0 \times 10^{-16}$ | $< 1.0 \times 10^{-16}$ | $< 1.0 \times 10^{-16}$ |

| CD4 T Naive | CD4 T Central Memory | CD4 T Effector Memory | CD8 T Naive | CD8 T Central Memory | CD8 T Effector Memory | CD56 Dim NK |
|---|---|---|---|---|---|---|
| # | | | | | | |
| $1.0 \times 10^{-1}$ | # | | | | | |
| $4.7 \times 10^{-1}$ | $1.0 \times 10^{-1}$ | # | | | | |
| $1.0 \times 10^{-1}$ | $1.0 \times 10^{-1}$ | $4.2 \times 10^{-1}$ | # | | | |
| $1.0 \times 10^{-1}$ | $1.0 \times 10^{-1}$ | $1.0 \times 10^{-1}$ | $1.0 \times 10^{-1}$ | # | | |
| $2.0 \times 10^{-1}$ | $8.3 \times 10^{-1}$ | $1.0 \times 10^{-1}$ | $1.7 \times 10^{-1}$ | $9.1 \times 10^{-1}$ | # | |
| $1.4 \times 10^{-1}$ | $7.2 \times 10^{-1}$ | $1.0 \times 10^{-1}$ | $1.1 \times 10^{-1}$ | $8.3 \times 10^{-1}$ | $1.0 \times 10^{-1}$ | # |
| $8.9 \times 10^{-1}$ | $1.0 \times 10^{-1}$ | $1.0 \times 10^{-1}$ | $8.6 \times 10^{-1}$ | $1.0 \times 10^{-1}$ | $1.0 \times 10^{-1}$ | $1.0 \times 10^{-1}$ |
| $9.1 \times 10^{-2}$ | $6.3 \times 10^{-1}$ | $1.0 \times 10^{-1}$ | $7.1 \times 10^{-2}$ | $7.6 \times 10^{-1}$ | $1.0 \times 10^{-1}$ | $1.0 \times 10^{-1}$ |
| $7.9 \times 10^{-1}$ | $1.0 \times 10^{-1}$ | $1.0 \times 10^{-1}$ | $7.5 \times 10^{-1}$ | $1.0 \times 10^{-1}$ | $1.0 \times 10^{-1}$ | $1.0 \times 10^{-1}$ |
| $6.2 \times 10^{-1}$ | $1.0 \times 10^{-1}$ | $1.0 \times 10^{-1}$ | $5.7 \times 10^{-1}$ | $1.0 \times 10^{-1}$ | $1.0 \times 10^{-1}$ | $1.0 \times 10^{-1}$ |
| $1.0 \times 10^{-1}$ | $6.3 \times 10^{-1}$ | $1.8 \times 10^{-2}$ | $1.0 \times 10^{-1}$ | $6.2 \times 10^{-1}$ | $3.6 \times 10^{-3}$ | $2.1 \times 10^{-3}$ |
| $8.0 \times 10^{-7}$ | $6.9 \times 10^{-5}$ | $3.8 \times 10^{-2}$ | $3.9 \times 10^{-7}$ | $2.7 \times 10^{-4}$ | $1.3 \times 10^{-1}$ | $2.6 \times 10^{-1}$ |
| $9.0 \times 10^{-2}$ | $5.9 \times 10^{-1}$ | $1.0 \times 10^{-1}$ | $7.2 \times 10^{-2}$ | $7.1 \times 10^{-1}$ | $1.0 \times 10^{-1}$ | $1.0 \times 10^{-1}$ |

| | | | | | | |
|---|---|---|---|---|---|---|
| # | | | | | | |
| $1.0 \times 10^{-1}$ | # | | | | | |
| $2.8 \times 10^{-1}$ | $8.8 \times 10^{-1}$ | # | | | | |
| $1.0 \times 10^{-1}$ | $1.0 \times 10^{-1}$ | $5.2 \times 10^{-2}$ | # | | | |
| $1.0 \times 10^{-1}$ | $1.0 \times 10^{-1}$ | $8.5 \times 10^{-1}$ | $1.0 \times 10^{-1}$ | # | | |
| $1.4 \times 10^{-1}$ | $7.0 \times 10^{-1}$ | $1.0 \times 10^{-1}$ | $1.9 \times 10^{-2}$ | $6.7 \times 10^{-1}$ | # | |
| $5.3 \times 10^{-1}$ | $1.0 \times 10^{-1}$ | $1.0 \times 10^{-1}$ | $1.5 \times 10^{-1}$ | $1.0 \times 10^{-1}$ | $1.0 \times 10^{-1}$ | # |
| $8.5 \times 10^{-1}$ | $1.0 \times 10^{-1}$ | $1.0 \times 10^{-1}$ | $4.3 \times 10^{-1}$ | $1.0 \times 10^{-1}$ | $1.0 \times 10^{-1}$ | $1.0 \times 10^{-1}$ |
| $5.0 \times 10^{-8}$ | $4.5 \times 10^{-6}$ | $2.2 \times 10^{-2}$ | $7.6 \times 10^{-10}$ | $7.4 \times 10^{-6}$ | $5.8 \times 10^{-2}$ | $9.4 \times 10^{-3}$ |
| $1.1 \times 10^{-2}$ | $1.7 \times 10^{-1}$ | $1.0 \times 10^{-1}$ | $7.9 \times 10^{-4}$ | $1.7 \times 10^{-1}$ | $1.0 \times 10^{-1}$ | $1.0 \times 10^{-1}$ |
| $1.8 \times 10^{-2}$ | $2.3 \times 10^{-1}$ | $1.0 \times 10^{-1}$ | $1.5 \times 10^{-3}$ | $2.2 \times 10^{-1}$ | $1.0 \times 10^{-1}$ | $1.0 \times 10^{-1}$ |
| $1.0 \times 10^{-1}$ | $4.8 \times 10^{-1}$ | $2.4 \times 10^{-3}$ | $1.0 \times 10^{-1}$ | $6.4 \times 10^{-1}$ | $6.6 \times 10^{-4}$ | $1.1 \times 10^{-2}$ |
| $< 1.0 \times 10^{-16}$ | $< 1.0 \times 10^{-16}$ | $6.0 \times 10^{-14}$ | $< 1.0 \times 10^{-16}$ | $< 1.0 \times 10^{-16}$ | $7.0 \times 10^{-14}$ | $4.5 \times 10^{-14}$ |
| $4.0 \times 10^{-1}$ | $9.4 \times 10^{-1}$ | $1.0 \times 10^{-1}$ | $1.0 \times 10^{-1}$ | $9.2 \times 10^{-1}$ | $1.0 \times 10^{-1}$ | $1.0 \times 10^{-1}$ |

| CD4 T Naive | CD4 T Central Memory | CD4 T Effector Memory | CD8 T Naive | CD8 T Central Memory | CD8 T Effector Memory | CD56 Dim NK |
|---|---|---|---|---|---|---|
| # | | | | | | |
| $2.9 \times 10^{-1}$ | # | | | | | |
| $3.8 \times 10^{-4}$ | $7.7 \times 10^{-1}$ | # | | | | |
| $1.0 \times 10^{-1}$ | $4.9 \times 10^{-1}$ | $1.1 \times 10^{-3}$ | # | | | |
| $8.7 \times 10^{-4}$ | $8.5 \times 10^{-1}$ | $1.0 \times 10^{-1}$ | $2.5 \times 10^{-3}$ | # | | |
| $2.1 \times 10^{-3}$ | $1.0 \times 10^{-1}$ | $1.0 \times 10^{-1}$ | $5.8 \times 10^{-3}$ | $1.0 \times 10^{-1}$ | # | |
| $9.3 \times 10^{-1}$ | $1.0 \times 10^{-1}$ | $2.1 \times 10^{-1}$ | $1.0 \times 10^{-1}$ | $2.9 \times 10^{-1}$ | $4.6 \times 10^{-1}$ | # |
| $8.4 \times 10^{-14}$ | $2.1 \times 10^{-9}$ | $1.4 \times 10^{-4}$ | $9.3 \times 10^{-14}$ | $1.4 \times 10^{-4}$ | $2.1 \times 10^{-5}$ | $5.1 \times 10^{-11}$ |
| $< 1.0 \times 10^{-16}$ | $< 1.0 \times 10^{-16}$ | $< 1.0 \times 10^{-16}$ | $< 1.0 \times 10^{-16}$ | $< 1.0 \times 10^{-16}$ | $< 1.0 \times 10^{-16}$ | $< 1.0 \times 10^{-16}$ |
| $< 1.0 \times 10^{-16}$ | $< 1.0 \times 10^{-16}$ | $< 1.0 \times 10^{-16}$ | $< 1.0 \times 10^{-16}$ | $< 1.0 \times 10^{-16}$ | $< 1.0 \times 10^{-16}$ | $< 1.0 \times 10^{-16}$ |
| $< 1.0 \times 10^{-16}$ | $< 1.0 \times 10^{-16}$ | $8.8 \times 10^{-14}$ | $< 1.0 \times 10^{-16}$ | $9.3 \times 10^{-14}$ | $9.2 \times 10^{-14}$ | $< 1.0 \times 10^{-16}$ |
| $< 1.0 \times 10^{-16}$ | $< 1.0 \times 10^{-16}$ | $< 1.0 \times 10^{-16}$ | $< 1.0 \times 10^{-16}$ | $< 1.0 \times 10^{-16}$ | $< 1.0 \times 10^{-16}$ | $< 1.0 \times 10^{-16}$ |
| $< 1.0 \times 10^{-16}$ | $< 1.0 \times 10^{-16}$ | $< 1.0 \times 10^{-16}$ | $< 1.0 \times 10^{-16}$ | $< 1.0 \times 10^{-16}$ | $< 1.0 \times 10^{-16}$ | $< 1.0 \times 10^{-16}$ |
| $< 1.0 \times 10^{-16}$ | $< 1.0 \times 10^{-16}$ | $< 1.0 \times 10^{-16}$ | $< 1.0 \times 10^{-16}$ | $< 1.0 \times 10^{-16}$ | $< 1.0 \times 10^{-16}$ | $< 1.0 \times 10^{-16}$ |

CD4 T Naive | CD4 T Central Memory | CD4 T Effector Memory | CD8 T Naive | CD8 T Central Memory | CD8 T Effector Memory | CD56 Dim NK

| | | | | | |
|---|---|---|---|---|---|
| # | | | | | |
| $1.0 \times 10^{-1}$ | # | | | | |
| $9.2 \times 10^{-1}$ | $1.0 \times 10^{-1}$ | # | | | |
| $1.0 \times 10^{-1}$ | $1.0 \times 10^{-1}$ | $1.0 \times 10^{-1}$ | # | | |
| $1.0 \times 10^{-1}$ | $1.0 \times 10^{-1}$ | $1.0 \times 10^{-1}$ | $1.0 \times 10^{-1}$ | # | |
| $1.0 \times 10^{-1}$ | $1.0 \times 10^{-1}$ | $1.0 \times 10^{-1}$ | $1.0 \times 10^{-1}$ | $1.0 \times 10^{-1}$ | # |

| CD4 T Naive | CD4 T Central Memory | CD4 T Effector Memory | CD8 T Naive | CD8 T Central Memory | CD8 T Effector Memory | CD56 Dim NK |
|---|---|---|---|---|---|---|
| $1.0 \times 10^{-1}$ | $1.0 \times 10^{-1}$ | $9.0 \times 10^{-1}$ | $1.0 \times 10^{-1}$ | $1.0 \times 10^{-1}$ | $1.0 \times 10^{-1}$ | # |
| $9.2 \times 10^{-1}$ | $2.8 \times 10^{-1}$ | $4.2 \times 10^{-2}$ | $1.7 \times 10^{-1}$ | $3.2 \times 10^{-1}$ | $1.3 \times 10^{-1}$ | $1.0 \times 10^{-1}$ |
| $1.0 \times 10^{-1}$ | $1.0 \times 10^{-1}$ | $1.0 \times 10^{-1}$ | $1.0 \times 10^{-1}$ | $1.0 \times 10^{-1}$ | $1.0 \times 10^{-1}$ | $1.0 \times 10^{-1}$ |
| $1.0 \times 10^{-1}$ | $1.0 \times 10^{-1}$ | $1.0 \times 10^{-1}$ | $1.0 \times 10^{-1}$ | $1.0 \times 10^{-1}$ | $1.0 \times 10^{-1}$ | $1.0 \times 10^{-1}$ |
| $1.0 \times 10^{-1}$ | $1.0 \times 10^{-1}$ | $1.0 \times 10^{-1}$ | $1.0 \times 10^{-1}$ | $1.0 \times 10^{-1}$ | $1.0 \times 10^{-1}$ | $1.0 \times 10^{-1}$ |
| $3.5 \times 10^{-5}$ | $1.1 \times 10^{-7}$ | $2.3 \times 10^{-9}$ | $3.0 \times 10^{-8}$ | $3.5 \times 10^{-7}$ | $2.5 \times 10^{-8}$ | $9.2 \times 10^{-5}$ |
| $3.5 \times 10^{-5}$ | $1.9 \times 10^{-3}$ | $4.4 \times 10^{-2}$ | $4.9 \times 10^{-3}$ | $4.3 \times 10^{-3}$ | $1.1 \times 10^{-2}$ | $3.8 \times 10^{-5}$ |
| $1.0 \times 10^{-1}$ | $1.0 \times 10^{-1}$ | $8.5 \times 10^{-1}$ | $1.0 \times 10^{-1}$ | $1.0 \times 10^{-1}$ | $1.0 \times 10^{-1}$ | $1.0 \times 10^{-1}$ |

| CD4 T Naive | CD4 T Central Memory | CD4 T Effector Memory | CD8 T Naive | CD8 T Central Memory | CD8 T Effector Memory | CD56 Dim NK |
|---|---|---|---|---|---|---|
| # | | | | | | |
| $1.0 \times 10^{-1}$ | # | | | | | |
| $1.0 \times 10^{-1}$ | $1.0 \times 10^{-1}$ | # | | | | |
| $1.0 \times 10^{-1}$ | $1.0 \times 10^{-1}$ | $1.0 \times 10^{-1}$ | # | | | |
| $1.0 \times 10^{-1}$ | $1.0 \times 10^{-1}$ | $1.0 \times 10^{-1}$ | $1.0 \times 10^{-1}$ | # | | |
| $1.0 \times 10^{-1}$ | $1.0 \times 10^{-1}$ | $1.0 \times 10^{-1}$ | $1.0 \times 10^{-1}$ | $1.0 \times 10^{-1}$ | # | |
| $1.0 \times 10^{-1}$ | $1.0 \times 10^{-1}$ | $1.0 \times 10^{-1}$ | $1.0 \times 10^{-1}$ | $1.0 \times 10^{-1}$ | $1.0 \times 10^{-1}$ | # |
| $1.0 \times 10^{-1}$ | $1.0 \times 10^{-1}$ | $1.0 \times 10^{-1}$ | $1.0 \times 10^{-1}$ | $1.0 \times 10^{-1}$ | $1.0 \times 10^{-1}$ | $1.0 \times 10^{-1}$ |
| $1.0 \times 10^{-1}$ | $1.0 \times 10^{-1}$ | $1.0 \times 10^{-1}$ | $1.0 \times 10^{-1}$ | $1.0 \times 10^{-1}$ | $1.0 \times 10^{-1}$ | $1.0 \times 10^{-1}$ |
| $1.0 \times 10^{-1}$ | $1.0 \times 10^{-1}$ | $1.0 \times 10^{-1}$ | $1.0 \times 10^{-1}$ | $1.0 \times 10^{-1}$ | $1.0 \times 10^{-1}$ | $1.0 \times 10^{-1}$ |
| $1.0 \times 10^{-1}$ | $1.0 \times 10^{-1}$ | $1.0 \times 10^{-1}$ | $1.0 \times 10^{-1}$ | $1.0 \times 10^{-1}$ | $1.0 \times 10^{-1}$ | $1.0 \times 10^{-1}$ |
| $1.0 \times 10^{-1}$ | $1.0 \times 10^{-1}$ | $1.0 \times 10^{-1}$ | $1.0 \times 10^{-1}$ | $1.0 \times 10^{-1}$ | $1.0 \times 10^{-1}$ | $1.0 \times 10^{-1}$ |
| $1.0 \times 10^{-1}$ | $1.0 \times 10^{-1}$ | $1.0 \times 10^{-1}$ | $1.0 \times 10^{-1}$ | $1.0 \times 10^{-1}$ | $1.0 \times 10^{-1}$ | $1.0 \times 10^{-1}$ |
| $1.0 \times 10^{-1}$ | $1.0 \times 10^{-1}$ | $1.0 \times 10^{-1}$ | $1.0 \times 10^{-1}$ | $1.0 \times 10^{-1}$ | $1.0 \times 10^{-1}$ | $1.0 \times 10^{-1}$ |

| CD4 T Naive | CD4 T Central Memory | CD4 T Effector Memory | CD8 T Naive | CD8 T Central Memory | CD8 T Effector Memory | CD56 Dim NK |
|---|---|---|---|---|---|---|
| # | | | | | | |
| $1.0 \times 10^{-1}$ | # | | | | | |
| $1.0 \times 10^{-1}$ | $1.0 \times 10^{-1}$ | # | | | | |
| $1.0 \times 10^{-1}$ | $1.0 \times 10^{-1}$ | $1.0 \times 10^{-1}$ | # | | | |
| $1.0 \times 10^{-1}$ | $1.0 \times 10^{-1}$ | $1.0 \times 10^{-1}$ | $1.0 \times 10^{-1}$ | # | | |
| $1.0 \times 10^{-1}$ | $1.0 \times 10^{-1}$ | $1.0 \times 10^{-1}$ | $1.0 \times 10^{-1}$ | $1.0 \times 10^{-1}$ | # | |
| $1.0 \times 10^{-1}$ | $1.0 \times 10^{-1}$ | $1.0 \times 10^{-1}$ | $1.0 \times 10^{-1}$ | $1.0 \times 10^{-1}$ | $1.0 \times 10^{-1}$ | # |
| $1.0 \times 10^{-1}$ | $1.0 \times 10^{-1}$ | $1.0 \times 10^{-1}$ | $1.0 \times 10^{-1}$ | $1.0 \times 10^{-1}$ | $1.0 \times 10^{-1}$ | $1.0 \times 10^{-1}$ |
| $1.0 \times 10^{-1}$ | $1.0 \times 10^{-1}$ | $1.0 \times 10^{-1}$ | $1.0 \times 10^{-1}$ | $1.0 \times 10^{-1}$ | $1.0 \times 10^{-1}$ | $1.0 \times 10^{-1}$ |
| $1.0 \times 10^{-1}$ | $1.0 \times 10^{-1}$ | $1.0 \times 10^{-1}$ | $1.0 \times 10^{-1}$ | $1.0 \times 10^{-1}$ | $1.0 \times 10^{-1}$ | $1.0 \times 10^{-1}$ |
| $1.0 \times 10^{-1}$ | $1.0 \times 10^{-1}$ | $1.0 \times 10^{-1}$ | $1.0 \times 10^{-1}$ | $1.0 \times 10^{-1}$ | $1.0 \times 10^{-1}$ | $1.0 \times 10^{-1}$ |
| $1.0 \times 10^{-1}$ | $1.0 \times 10^{-1}$ | $1.0 \times 10^{-1}$ | $1.0 \times 10^{-1}$ | $1.0 \times 10^{-1}$ | $1.0 \times 10^{-1}$ | $1.0 \times 10^{-1}$ |
| $1.0 \times 10^{-1}$ | $1.0 \times 10^{-1}$ | $1.0 \times 10^{-1}$ | $1.0 \times 10^{-1}$ | $1.0 \times 10^{-1}$ | $1.0 \times 10^{-1}$ | $1.0 \times 10^{-1}$ |
| $1.0 \times 10^{-1}$ | $1.0 \times 10^{-1}$ | $1.0 \times 10^{-1}$ | $1.0 \times 10^{-1}$ | $1.0 \times 10^{-1}$ | $1.0 \times 10^{-1}$ | $1.0 \times 10^{-1}$ |

| | | | | | | |
|---|---|---|---|---|---|---|
| # | | | | | | |
| $1.0 \times 10^{-1}$ | # | | | | | |
| $1.0 \times 10^{-1}$ | $1.0 \times 10^{-1}$ | # | | | | |
| $1.0 \times 10^{-1}$ | $1.0 \times 10^{-1}$ | $1.0 \times 10^{-1}$ | # | | | |
| $1.0 \times 10^{-1}$ | $1.0 \times 10^{-1}$ | $1.0 \times 10^{-1}$ | $1.0 \times 10^{-1}$ | # | | |
| $1.0 \times 10^{-1}$ | $1.0 \times 10^{-1}$ | $1.0 \times 10^{-1}$ | $1.0 \times 10^{-1}$ | $1.0 \times 10^{-1}$ | # | |
| $1.0 \times 10^{-1}$ | $1.0 \times 10^{-1}$ | $1.0 \times 10^{-1}$ | $1.0 \times 10^{-1}$ | $1.0 \times 10^{-1}$ | $1.0 \times 10^{-1}$ | # |
| $1.0 \times 10^{-1}$ | $1.0 \times 10^{-1}$ | $1.0 \times 10^{-1}$ | $1.0 \times 10^{-1}$ | $1.0 \times 10^{-1}$ | $1.0 \times 10^{-1}$ | $1.0 \times 10^{-1}$ |
| $1.0 \times 10^{-1}$ | $1.0 \times 10^{-1}$ | $1.0 \times 10^{-1}$ | $1.0 \times 10^{-1}$ | $1.0 \times 10^{-1}$ | $1.0 \times 10^{-1}$ | $1.0 \times 10^{-1}$ |
| $1.0 \times 10^{-1}$ | $1.0 \times 10^{-1}$ | $1.0 \times 10^{-1}$ | $1.0 \times 10^{-1}$ | $1.0 \times 10^{-1}$ | $1.0 \times 10^{-1}$ | $1.0 \times 10^{-1}$ |
| $1.0 \times 10^{-1}$ | $1.0 \times 10^{-1}$ | $1.0 \times 10^{-1}$ | $1.0 \times 10^{-1}$ | $1.0 \times 10^{-1}$ | $1.0 \times 10^{-1}$ | $1.0 \times 10^{-1}$ |
| $1.0 \times 10^{-1}$ | $1.0 \times 10^{-1}$ | $1.0 \times 10^{-1}$ | $1.0 \times 10^{-1}$ | $1.0 \times 10^{-1}$ | $1.0 \times 10^{-1}$ | $1.0 \times 10^{-1}$ |
| $1.0 \times 10^{-1}$ | $1.0 \times 10^{-1}$ | $1.0 \times 10^{-1}$ | $1.0 \times 10^{-1}$ | $1.0 \times 10^{-1}$ | $1.0 \times 10^{-1}$ | $1.0 \times 10^{-1}$ |
| $1.0 \times 10^{-1}$ | $1.0 \times 10^{-1}$ | $1.0 \times 10^{-1}$ | $1.0 \times 10^{-1}$ | $1.0 \times 10^{-1}$ | $1.0 \times 10^{-1}$ | $1.0 \times 10^{-1}$ |

| CD4 T Naive | CD4 T Central Memory | CD4 T Effector Memory | CD8 T Naive | CD8 T Central Memory | CD8 T Effector Memory | CD56 Dim NK |
|---|---|---|---|---|---|---|
| # | | | | | | |
| $1.0 \times 10^{-1}$ | # | | | | | |
| $1.0 \times 10^{-1}$ | $1.0 \times 10^{-1}$ | # | | | | |
| $1.0 \times 10^{-1}$ | $1.0 \times 10^{-1}$ | $1.0 \times 10^{-1}$ | # | | | |
| $1.0 \times 10^{-1}$ | $1.0 \times 10^{-1}$ | $1.0 \times 10^{-1}$ | $1.0 \times 10^{-1}$ | # | | |
| $1.0 \times 10^{-1}$ | $1.0 \times 10^{-1}$ | $1.0 \times 10^{-1}$ | $1.0 \times 10^{-1}$ | $1.0 \times 10^{-1}$ | # | |
| $1.0 \times 10^{-1}$ | $1.0 \times 10^{-1}$ | $1.0 \times 10^{-1}$ | $1.0 \times 10^{-1}$ | $1.0 \times 10^{-1}$ | $1.0 \times 10^{-1}$ | # |
| $1.0 \times 10^{-1}$ | $1.0 \times 10^{-1}$ | $1.0 \times 10^{-1}$ | $1.0 \times 10^{-1}$ | $1.0 \times 10^{-1}$ | $1.0 \times 10^{-1}$ | $1.0 \times 10^{-1}$ |
| $1.0 \times 10^{-1}$ | $1.0 \times 10^{-1}$ | $1.0 \times 10^{-1}$ | $1.0 \times 10^{-1}$ | $1.0 \times 10^{-1}$ | $1.0 \times 10^{-1}$ | $1.0 \times 10^{-1}$ |
| $1.0 \times 10^{-1}$ | $1.0 \times 10^{-1}$ | $1.0 \times 10^{-1}$ | $1.0 \times 10^{-1}$ | $1.0 \times 10^{-1}$ | $1.0 \times 10^{-1}$ | $1.0 \times 10^{-1}$ |
| $1.0 \times 10^{-1}$ | $1.0 \times 10^{-1}$ | $1.0 \times 10^{-1}$ | $1.0 \times 10^{-1}$ | $1.0 \times 10^{-1}$ | $1.0 \times 10^{-1}$ | $1.0 \times 10^{-1}$ |
| $1.0 \times 10^{-1}$ | $1.0 \times 10^{-1}$ | $1.0 \times 10^{-1}$ | $1.0 \times 10^{-1}$ | $1.0 \times 10^{-1}$ | $1.0 \times 10^{-1}$ | $1.0 \times 10^{-1}$ |
| $1.0 \times 10^{-1}$ | $1.0 \times 10^{-1}$ | $1.0 \times 10^{-1}$ | $1.0 \times 10^{-1}$ | $1.0 \times 10^{-1}$ | $1.0 \times 10^{-1}$ | $1.0 \times 10^{-1}$ |
| $1.0 \times 10^{-1}$ | $1.0 \times 10^{-1}$ | $1.0 \times 10^{-1}$ | $1.0 \times 10^{-1}$ | $1.0 \times 10^{-1}$ | $1.0 \times 10^{-1}$ | $1.0 \times 10^{-1}$ |

| CD4 T Naive | CD4 T Central Memory | CD4 T Effector Memory | CD8 T Naive | CD8 T Central Memory | CD8 T Effector Memory | CD56 Dim NK |
|---|---|---|---|---|---|---|
| # | | | | | | |
| $2.6 \times 10^{-1}$ | # | | | | | |
| $4.0 \times 10^{-3}$ | $1.0 \times 10^{-1}$ | # | | | | |
| $3.7 \times 10^{-4}$ | $5.5 \times 10^{-11}$ | $1.0 \times 10^{-13}$ | # | | | |
| $9.4 \times 10^{-1}$ | $1.0 \times 10^{-1}$ | $5.4 \times 10^{-1}$ | $1.2 \times 10^{-7}$ | # | | |
| $6.4 \times 10^{-1}$ | $1.0 \times 10^{-1}$ | $8.6 \times 10^{-1}$ | $2.8 \times 10^{-9}$ | $1.0 \times 10^{-1}$ | # | |

| CD4 T Naive | CD4 T Central Memory | CD4 T Effector Memory | CD8 T Naive | CD8 T Central Memory | CD8 T Effector Memory | CD56 Dim NK |
|---|---|---|---|---|---|---|
| $1.0 \times 10^{-1}$ | $7.0 \times 10^{-2}$ | $5.7 \times 10^{-4}$ | $5.8 \times 10^{-3}$ | $6.8 \times 10^{-1}$ | $2.8 \times 10^{-1}$ | # |
| $1.0 \times 10^{-1}$ | $4.5 \times 10^{-1}$ | $1.3 \times 10^{-2}$ | $1.9 \times 10^{-4}$ | $1.0 \times 10^{-1}$ | $8.3 \times 10^{-1}$ | $1.0 \times 10^{-1}$ |
| $1.0 \times 10^{-1}$ | $1.0 \times 10^{-1}$ | $3.6 \times 10^{-1}$ | $7.2 \times 10^{-8}$ | $1.0 \times 10^{-1}$ | $1.0 \times 10^{-1}$ | $7.3 \times 10^{-1}$ |
| $3.8 \times 10^{-1}$ | $1.0 \times 10^{-1}$ | $1.0 \times 10^{-1}$ | $1.8 \times 10^{-10}$ | $1.0 \times 10^{-1}$ | $1.0 \times 10^{-1}$ | $1.2 \times 10^{-1}$ |
| $1.0 \times 10^{-2}$ | $1.0 \times 10^{-1}$ | $1.0 \times 10^{-1}$ | $1.8 \times 10^{-13}$ | $7.2 \times 10^{-1}$ | $1.0 \times 10^{-1}$ | $1.6 \times 10^{-3}$ |
| $7.3 \times 10^{-5}$ | $5.9 \times 10^{-12}$ | $9.0 \times 10^{-14}$ | $1.0 \times 10^{-1}$ | $1.8 \times 10^{-8}$ | $3.4 \times 10^{-10}$ | $1.4 \times 10^{-3}$ |
| $2.1 \times 10^{-8}$ | $7.8 \times 10^{-3}$ | $4.5 \times 10^{-1}$ | $< 1.0 \times 10^{-16}$ | $2.2 \times 10^{-4}$ | $1.4 \times 10^{-3}$ | $1.8 \times 10^{-9}$ |
| $8.4 \times 10^{-14}$ | $1.8 \times 10^{-8}$ | $3.7 \times 10^{-5}$ | $< 1.0 \times 10^{-16}$ | $2.5 \times 10^{-10}$ | $2.1 \times 10^{-9}$ | $8.4 \times 10^{-14}$ |

| CD4 T Naive | CD4 T Central Memory | CD4 T Effector Memory | CD8 T Naive | CD8 T Central Memory | CD8 T Effector Memory | CD56 Dim NK |
|---|---|---|---|---|---|---|
| # | | | | | | |
| $1.0 \times 10^{-1}$ | # | | | | | |
| $1.0 \times 10^{-1}$ | $1.0 \times 10^{-1}$ | # | | | | |
| $1.0 \times 10^{-1}$ | $1.0 \times 10^{-1}$ | $1.0 \times 10^{-1}$ | # | | | |
| $1.0 \times 10^{-1}$ | $1.0 \times 10^{-1}$ | $1.0 \times 10^{-1}$ | $1.0 \times 10^{-1}$ | # | | |
| $1.0 \times 10^{-1}$ | $1.0 \times 10^{-1}$ | $1.0 \times 10^{-1}$ | $1.0 \times 10^{-1}$ | $1.0 \times 10^{-1}$ | # | |
| $1.0 \times 10^{-1}$ | $1.0 \times 10^{-1}$ | $1.0 \times 10^{-1}$ | $1.0 \times 10^{-1}$ | $1.0 \times 10^{-1}$ | $1.0 \times 10^{-1}$ | # |
| $1.0 \times 10^{-1}$ | $1.0 \times 10^{-1}$ | $1.0 \times 10^{-1}$ | $1.0 \times 10^{-1}$ | $1.0 \times 10^{-1}$ | $1.0 \times 10^{-1}$ | $1.0 \times 10^{-1}$ |
| $1.0 \times 10^{-1}$ | $1.0 \times 10^{-1}$ | $1.0 \times 10^{-1}$ | $1.0 \times 10^{-1}$ | $1.0 \times 10^{-1}$ | $1.0 \times 10^{-1}$ | $1.0 \times 10^{-1}$ |
| $1.0 \times 10^{-1}$ | $1.0 \times 10^{-1}$ | $1.0 \times 10^{-1}$ | $1.0 \times 10^{-1}$ | $1.0 \times 10^{-1}$ | $1.0 \times 10^{-1}$ | $1.0 \times 10^{-1}$ |
| $1.0 \times 10^{-1}$ | $1.0 \times 10^{-1}$ | $1.0 \times 10^{-1}$ | $1.0 \times 10^{-1}$ | $1.0 \times 10^{-1}$ | $1.0 \times 10^{-1}$ | $1.0 \times 10^{-1}$ |
| $1.0 \times 10^{-1}$ | $1.0 \times 10^{-1}$ | $1.0 \times 10^{-1}$ | $1.0 \times 10^{-1}$ | $1.0 \times 10^{-1}$ | $1.0 \times 10^{-1}$ | $1.0 \times 10^{-1}$ |
| $1.0 \times 10^{-1}$ | $1.0 \times 10^{-1}$ | $1.0 \times 10^{-1}$ | $1.0 \times 10^{-1}$ | $1.0 \times 10^{-1}$ | $1.0 \times 10^{-1}$ | $1.0 \times 10^{-1}$ |
| $1.0 \times 10^{-1}$ | $1.0 \times 10^{-1}$ | $1.0 \times 10^{-1}$ | $1.0 \times 10^{-1}$ | $1.0 \times 10^{-1}$ | $1.0 \times 10^{-1}$ | $1.0 \times 10^{-1}$ |

| CD4 T Naive | CD4 T Central Memory | CD4 T Effector Memory | CD8 T Naive | CD8 T Central Memory | CD8 T Effector Memory | CD56 Dim NK |
|---|---|---|---|---|---|---|
| # | | | | | | |
| $1.0 \times 10^{-1}$ | # | | | | | |
| $2.9 \times 10^{-2}$ | $6.9 \times 10^{-1}$ | # | | | | |
| $1.0 \times 10^{-1}$ | $7.7 \times 10^{-1}$ | $3.4 \times 10^{-3}$ | # | | | |
| $4.5 \times 10^{-1}$ | $1.0 \times 10^{-1}$ | $1.0 \times 10^{-1}$ | $1.4 \times 10^{-1}$ | # | | |
| $1.1 \times 10^{-2}$ | $4.8 \times 10^{-1}$ | $1.0 \times 10^{-1}$ | $1.1 \times 10^{-3}$ | $1.0 \times 10^{-1}$ | # | |
| $7.2 \times 10^{-5}$ | $2.0 \times 10^{-2}$ | $1.0 \times 10^{-1}$ | $3.7 \times 10^{-6}$ | $4.3 \times 10^{-1}$ | $1.0 \times 10^{-1}$ | # |
| $7.3 \times 10^{-14}$ | $1.0 \times 10^{-13}$ | $1.8 \times 10^{-8}$ | $2.0 \times 10^{-14}$ | $1.0 \times 10^{-10}$ | $8.0 \times 10^{-8}$ | $9.4 \times 10^{-5}$ |
| $< 1.0 \times 10^{-16}$ | $8.3 \times 10^{-14}$ | $9.7 \times 10^{-11}$ | $< 1.0 \times 10^{-16}$ | $4.8 \times 10^{-13}$ | $5.2 \times 10^{-10}$ | $1.8 \times 10^{-6}$ |
| $< 1.0 \times 10^{-16}$ | $7.8 \times 10^{-14}$ | $8.5 \times 10^{-11}$ | $< 1.0 \times 10^{-16}$ | $4.3 \times 10^{-13}$ | $4.6 \times 10^{-10}$ | $1.6 \times 10^{-6}$ |
| $1.8 \times 10^{-13}$ | $3.7 \times 10^{-10}$ | $7.3 \times 10^{-5}$ | $8.1 \times 10^{-14}$ | $7.2 \times 10^{-7}$ | $2.6 \times 10^{-4}$ | $5.1 \times 10^{-2}$ |
| $< 1.0 \times 10^{-16}$ | $< 1.0 \times 10^{-16}$ | $< 1.0 \times 10^{-16}$ | $< 1.0 \times 10^{-16}$ | $< 1.0 \times 10^{-16}$ | $< 1.0 \times 10^{-16}$ | $< 1.0 \times 10^{-16}$ |
| $< 1.0 \times 10^{-16}$ | $< 1.0 \times 10^{-16}$ | $< 1.0 \times 10^{-16}$ | $< 1.0 \times 10^{-16}$ | $< 1.0 \times 10^{-16}$ | $< 1.0 \times 10^{-16}$ | $< 1.0 \times 10^{-16}$ |
| $< 1.0 \times 10^{-16}$ | $< 1.0 \times 10^{-16}$ | $< 1.0 \times 10^{-16}$ | $< 1.0 \times 10^{-16}$ | $< 1.0 \times 10^{-16}$ | $< 1.0 \times 10^{-16}$ | $< 1.0 \times 10^{-16}$ |

| | | | | | | |
|---|---|---|---|---|---|---|
| # | | | | | | |
| $1.0 \times 10^{-1}$ | # | | | | | |
| $1.0 \times 10^{-1}$ | $1.0 \times 10^{-1}$ | # | | | | |
| $1.0 \times 10^{-1}$ | $1.0 \times 10^{-1}$ | $1.0 \times 10^{-1}$ | # | | | |
| $1.0 \times 10^{-1}$ | $1.0 \times 10^{-1}$ | $1.0 \times 10^{-1}$ | $1.0 \times 10^{-1}$ | # | | |
| $1.0 \times 10^{-1}$ | $1.0 \times 10^{-1}$ | $1.0 \times 10^{-1}$ | $1.0 \times 10^{-1}$ | $1.0 \times 10^{-1}$ | # | |
| $1.0 \times 10^{-1}$ | $1.0 \times 10^{-1}$ | $1.0 \times 10^{-1}$ | $1.0 \times 10^{-1}$ | $1.0 \times 10^{-1}$ | $1.0 \times 10^{-1}$ | # |
| $1.0 \times 10^{-1}$ | $1.0 \times 10^{-1}$ | $1.0 \times 10^{-1}$ | $1.0 \times 10^{-1}$ | $1.0 \times 10^{-1}$ | $1.0 \times 10^{-1}$ | $1.0 \times 10^{-1}$ |
| $6.6 \times 10^{-1}$ | $9.1 \times 10^{-1}$ | $1.0 \times 10^{-1}$ | $7.8 \times 10^{-1}$ | $1.0 \times 10^{-1}$ | $1.0 \times 10^{-1}$ | $8.8 \times 10^{-1}$ |
| $6.1 \times 10^{-1}$ | $8.9 \times 10^{-1}$ | $1.0 \times 10^{-1}$ | $7.3 \times 10^{-1}$ | $1.0 \times 10^{-1}$ | $1.0 \times 10^{-1}$ | $8.5 \times 10^{-1}$ |
| $5.1 \times 10^{-1}$ | $8.1 \times 10^{-1}$ | $1.0 \times 10^{-1}$ | $6.3 \times 10^{-1}$ | $1.0 \times 10^{-1}$ | $9.3 \times 10^{-1}$ | $7.7 \times 10^{-1}$ |
| $1.0 \times 10^{-1}$ | $1.0 \times 10^{-1}$ | $1.0 \times 10^{-1}$ | $1.0 \times 10^{-1}$ | $1.0 \times 10^{-1}$ | $1.0 \times 10^{-1}$ | $1.0 \times 10^{-1}$ |
| $1.1 \times 10^{-1}$ | $3.0 \times 10^{-1}$ | $6.7 \times 10^{-1}$ | $1.7 \times 10^{-1}$ | $7.4 \times 10^{-1}$ | $4.8 \times 10^{-1}$ | $2.8 \times 10^{-1}$ |
| $2.0 \times 10^{-1}$ | $4.4 \times 10^{-1}$ | $7.9 \times 10^{-1}$ | $2.7 \times 10^{-1}$ | $8.5 \times 10^{-1}$ | $6.3 \times 10^{-1}$ | $4.1 \times 10^{-1}$ |

| CD4 T Naive | CD4 T Central Memory | CD4 T Effector Memory | CD8 T Naive | CD8 T Central Memory | CD8 T Effector Memory | CD56 Dim NK |
|---|---|---|---|---|---|---|
| # | | | | | | |
| $1.0 \times 10^{-1}$ | # | | | | | |
| $1.0 \times 10^{-1}$ | $1.0 \times 10^{-1}$ | # | | | | |
| $1.0 \times 10^{-1}$ | $1.0 \times 10^{-1}$ | $1.0 \times 10^{-1}$ | # | | | |
| $1.0 \times 10^{-1}$ | $1.0 \times 10^{-1}$ | $1.0 \times 10^{-1}$ | $1.0 \times 10^{-1}$ | # | | |
| $1.0 \times 10^{-1}$ | $1.0 \times 10^{-1}$ | $1.0 \times 10^{-1}$ | $1.0 \times 10^{-1}$ | $1.0 \times 10^{-1}$ | # | |
| $6.5 \times 10^{-4}$ | $3.5 \times 10^{-5}$ | $1.4 \times 10^{-6}$ | $9.4 \times 10^{-5}$ | $2.2 \times 10^{-3}$ | $1.6 \times 10^{-6}$ | # |
| $7.7 \times 10^{-5}$ | $3.1 \times 10^{-6}$ | $1.1 \times 10^{-7}$ | $9.1 \times 10^{-6}$ | $3.1 \times 10^{-4}$ | $1.3 \times 10^{-7}$ | $1.0 \times 10^{-1}$ |
| $1.7 \times 10^{-6}$ | $1.9 \times 10^{-5}$ | $9.4 \times 10^{-4}$ | $6.4 \times 10^{-6}$ | $1.1 \times 10^{-6}$ | $8.4 \times 10^{-4}$ | $6.1 \times 10^{-14}$ |
| $9.3 \times 10^{-6}$ | $9.4 \times 10^{-5}$ | $3.6 \times 10^{-3}$ | $3.4 \times 10^{-5}$ | $5.6 \times 10^{-6}$ | $3.2 \times 10^{-3}$ | $8.2 \times 10^{-14}$ |
| $6.6 \times 10^{-6}$ | $6.6 \times 10^{-5}$ | $2.5 \times 10^{-3}$ | $2.4 \times 10^{-5}$ | $4.0 \times 10^{-6}$ | $2.2 \times 10^{-3}$ | $7.7 \times 10^{-14}$ |
| $1.0 \times 10^{-1}$ | $1.0 \times 10^{-1}$ | $1.0 \times 10^{-1}$ | $1.0 \times 10^{-1}$ | $1.0 \times 10^{-1}$ | $1.0 \times 10^{-1}$ | $4.7 \times 10^{-7}$ |
| $1.8 \times 10^{-1}$ | $5.0 \times 10^{-1}$ | $1.0 \times 10^{-1}$ | $3.5 \times 10^{-1}$ | $1.2 \times 10^{-1}$ | $9.5 \times 10^{-1}$ | $1.6 \times 10^{-10}$ |
| $5.2 \times 10^{-13}$ | $5.7 \times 10^{-12}$ | $9.4 \times 10^{-10}$ | $1.6 \times 10^{-12}$ | $4.0 \times 10^{-13}$ | $8.0 \times 10^{-10}$ | $< 1.0 \times 10^{-16}$ |

| CD4 T Naive | CD4 T Central Memory | CD4 T Effector Memory | CD8 T Naive | CD8 T Central Memory | CD8 T Effector Memory | CD56 Dim NK |
|---|---|---|---|---|---|---|
| # | | | | | | |
| $4.7 \times 10^{-1}$ | # | | | | | |
| $1.4 \times 10^{-11}$ | $7.4 \times 10^{-6}$ | # | | | | |
| $1.0 \times 10^{-1}$ | $1.7 \times 10^{-1}$ | $4.1 \times 10^{-13}$ | # | | | |
| $3.6 \times 10^{-1}$ | $1.0 \times 10^{-1}$ | $6.9 \times 10^{-5}$ | $1.2 \times 10^{-1}$ | # | | |
| $8.4 \times 10^{-14}$ | $2.3 \times 10^{-9}$ | $1.0 \times 10^{-1}$ | $8.0 \times 10^{-14}$ | $4.4 \times 10^{-8}$ | # | |

| CD4 T Naive | CD4 T Central Memory | CD4 T Effector Memory | CD8 T Naive | CD8 T Central Memory | CD8 T Effector Memory | CD56 Dim NK |
|---|---|---|---|---|---|---|
| $< 1.0 \times 10^{-16}$ | $4.7 \times 10^{-14}$ | $8.2 \times 10^{-5}$ | $< 1.0 \times 10^{-16}$ | $7.8 \times 10^{-14}$ | $2.6 \times 10^{-2}$ | # |
| $< 1.0 \times 10^{-16}$ | $< 1.0 \times 10^{-16}$ | $1.1 \times 10^{-7}$ | $< 1.0 \times 10^{-16}$ | $< 1.0 \times 10^{-16}$ | $1.5 \times 10^{-4}$ | $1.0 \times 10^{-1}$ |
| $1.8 \times 10^{-5}$ | $1.9 \times 10^{-1}$ | $3.4 \times 10^{-1}$ | $1.1 \times 10^{-6}$ | $4.0 \times 10^{-1}$ | $4.0 \times 10^{-3}$ | $3.2 \times 10^{-11}$ |
| $4.0 \times 10^{-8}$ | $4.3 \times 10^{-3}$ | $1.0 \times 10^{-1}$ | $1.4 \times 10^{-9}$ | $2.0 \times 10^{-2}$ | $1.7 \times 10^{-1}$ | $2.6 \times 10^{-8}$ |
| $9.5 \times 10^{-14}$ | $1.5 \times 10^{-8}$ | $1.0 \times 10^{-1}$ | $9.0 \times 10^{-14}$ | $2.5 \times 10^{-7}$ | $1.0 \times 10^{-1}$ | $8.8 \times 10^{-3}$ |
| $1.0 \times 10^{-1}$ | $6.4 \times 10^{-1}$ | $3.1 \times 10^{-11}$ | $1.0 \times 10^{-1}$ | $5.2 \times 10^{-1}$ | $8.7 \times 10^{-14}$ | $< 1.0 \times 10^{-16}$ |
| $< 1.0 \times 10^{-16}$ | $< 1.0 \times 10^{-16}$ | $8.8 \times 10^{-14}$ | $< 1.0 \times 10^{-16}$ | $< 1.0 \times 10^{-16}$ | $6.5 \times 10^{-12}$ | $3.0 \times 10^{-3}$ |
| $1.0 \times 10^{-1}$ | $5.7 \times 10^{-1}$ | $1.3 \times 10^{-10}$ | $1.0 \times 10^{-1}$ | $4.6 \times 10^{-1}$ | $1.1 \times 10^{-13}$ | $< 1.0 \times 10^{-16}$ |

| CD4 T Naive | CD4 T Central Memory | CD4 T Effector Memory | CD8 T Naive | CD8 T Central Memory | CD8 T Effector Memory | CD56 Dim NK |
|---|---|---|---|---|---|---|
| # | | | | | | |
| $1.0 \times 10^{-1}$ | # | | | | | |
| $1.0 \times 10^{-1}$ | $1.0 \times 10^{-1}$ | # | | | | |
| $1.0 \times 10^{-1}$ | $1.0 \times 10^{-1}$ | $1.0 \times 10^{-1}$ | # | | | |
| $1.0 \times 10^{-1}$ | $1.0 \times 10^{-1}$ | $1.0 \times 10^{-1}$ | $1.0 \times 10^{-1}$ | # | | |
| $1.0 \times 10^{-1}$ | $1.0 \times 10^{-1}$ | $1.0 \times 10^{-1}$ | $1.0 \times 10^{-1}$ | $1.0 \times 10^{-1}$ | # | |
| $1.0 \times 10^{-1}$ | $1.0 \times 10^{-1}$ | $1.0 \times 10^{-1}$ | $1.0 \times 10^{-1}$ | $1.0 \times 10^{-1}$ | $1.0 \times 10^{-1}$ | # |
| $1.0 \times 10^{-1}$ | $1.0 \times 10^{-1}$ | $1.0 \times 10^{-1}$ | $1.0 \times 10^{-1}$ | $1.0 \times 10^{-1}$ | $1.0 \times 10^{-1}$ | $1.0 \times 10^{-1}$ |
| $1.0 \times 10^{-1}$ | $1.0 \times 10^{-1}$ | $1.0 \times 10^{-1}$ | $1.0 \times 10^{-1}$ | $1.0 \times 10^{-1}$ | $1.0 \times 10^{-1}$ | $1.0 \times 10^{-1}$ |
| $1.0 \times 10^{-1}$ | $1.0 \times 10^{-1}$ | $1.0 \times 10^{-1}$ | $1.0 \times 10^{-1}$ | $1.0 \times 10^{-1}$ | $1.0 \times 10^{-1}$ | $1.0 \times 10^{-1}$ |
| $1.0 \times 10^{-1}$ | $1.0 \times 10^{-1}$ | $1.0 \times 10^{-1}$ | $1.0 \times 10^{-1}$ | $1.0 \times 10^{-1}$ | $1.0 \times 10^{-1}$ | $1.0 \times 10^{-1}$ |
| $1.0 \times 10^{-1}$ | $1.0 \times 10^{-1}$ | $1.0 \times 10^{-1}$ | $1.0 \times 10^{-1}$ | $1.0 \times 10^{-1}$ | $1.0 \times 10^{-1}$ | $1.0 \times 10^{-1}$ |
| $1.0 \times 10^{-1}$ | $1.0 \times 10^{-1}$ | $1.0 \times 10^{-1}$ | $1.0 \times 10^{-1}$ | $1.0 \times 10^{-1}$ | $1.0 \times 10^{-1}$ | $1.0 \times 10^{-1}$ |
| $1.0 \times 10^{-1}$ | $1.0 \times 10^{-1}$ | $1.0 \times 10^{-1}$ | $1.0 \times 10^{-1}$ | $1.0 \times 10^{-1}$ | $1.0 \times 10^{-1}$ | $1.0 \times 10^{-1}$ |

| CD4 T Naive | CD4 T Central Memory | CD4 T Effector Memory | CD8 T Naive | CD8 T Central Memory | CD8 T Effector Memory | CD56 Dim NK |
|---|---|---|---|---|---|---|
| # | | | | | | |
| $1.0 \times 10^{-1}$ | # | | | | | |
| $1.6 \times 10^{-1}$ | $7.9 \times 10^{-1}$ | # | | | | |
| $1.0 \times 10^{-1}$ | $1.0 \times 10^{-1}$ | $3.5 \times 10^{-1}$ | # | | | |
| $1.0 \times 10^{-1}$ | $1.0 \times 10^{-1}$ | $7.1 \times 10^{-1}$ | $1.0 \times 10^{-1}$ | # | | |
| $5.7 \times 10^{-1}$ | $1.0 \times 10^{-1}$ | $1.0 \times 10^{-1}$ | $8.3 \times 10^{-1}$ | $1.0 \times 10^{-1}$ | # | |
| $1.0 \times 10^{-1}$ | $1.0 \times 10^{-1}$ | $4.4 \times 10^{-1}$ | $1.0 \times 10^{-1}$ | $1.0 \times 10^{-1}$ | $8.8 \times 10^{-1}$ | # |
| $1.0 \times 10^{-1}$ | $1.0 \times 10^{-1}$ | $3.5 \times 10^{-1}$ | $1.0 \times 10^{-1}$ | $1.0 \times 10^{-1}$ | $8.1 \times 10^{-1}$ | $1.0 \times 10^{-1}$ |
| $5.4 \times 10^{-4}$ | $2.3 \times 10^{-2}$ | $1.0 \times 10^{-1}$ | $2.3 \times 10^{-3}$ | $1.9 \times 10^{-2}$ | $6.4 \times 10^{-1}$ | $5.0 \times 10^{-3}$ |
| $4.8 \times 10^{-3}$ | $1.2 \times 10^{-1}$ | $1.0 \times 10^{-1}$ | $1.7 \times 10^{-2}$ | $9.4 \times 10^{-2}$ | $9.3 \times 10^{-1}$ | $3.2 \times 10^{-2}$ |
| $3.1 \times 10^{-5}$ | $2.0 \times 10^{-3}$ | $6.4 \times 10^{-1}$ | $1.4 \times 10^{-4}$ | $1.7 \times 10^{-3}$ | $2.0 \times 10^{-1}$ | $3.8 \times 10^{-4}$ |
| $6.6 \times 10^{-4}$ | $3.0 \times 10^{-6}$ | $5.7 \times 10^{-11}$ | $6.3 \times 10^{-5}$ | $2.5 \times 10^{-5}$ | $3.3 \times 10^{-9}$ | $1.4 \times 10^{-4}$ |
| $6.9 \times 10^{-1}$ | $8.4 \times 10^{-2}$ | $4.4 \times 10^{-5}$ | $3.5 \times 10^{-1}$ | $1.9 \times 10^{-1}$ | $8.9 \times 10^{-4}$ | $4.0 \times 10^{-1}$ |
| $1.6 \times 10^{-3}$ | $1.3 \times 10^{-5}$ | $6.0 \times 10^{-10}$ | $2.0 \times 10^{-4}$ | $7.9 \times 10^{-5}$ | $2.4 \times 10^{-8}$ | $3.7 \times 10^{-4}$ |

**ıan immune cells**

| | | | | | | |
|---|---|---|---|---|---|---|
| # | | | | | | |
| 1.0 x 10⁻¹ | # | | | | | |
| 1.0 x 10⁻¹ | 1.0 x 10⁻¹ | # | | | | |
| 1.0 x 10⁻¹ | 1.0 x 10⁻¹ | 1.0 x 10⁻¹ | # | | | |
| 1.0 x 10⁻¹ | 1.0 x 10⁻¹ | 1.0 x 10⁻¹ | 1.0 x 10⁻¹ | # | | |
| 1.0 x 10⁻¹ | 1.0 x 10⁻¹ | 1.0 x 10⁻¹ | 1.0 x 10⁻¹ | 1.0 x 10⁻¹ | # | |
| 1.0 x 10⁻¹ | 1.0 x 10⁻¹ | 1.0 x 10⁻¹ | 1.0 x 10⁻¹ | 1.0 x 10⁻¹ | 1.0 x 10⁻¹ | # |
| CD56 Bright NK | Classical Monocyte | Intermediate Monocyte | Non-classical Monocyte | Basophil | Eosinophil | Neutrophil |

| | | | | | |
|---|---|---|---|---|---|
| # | | | | | |
| 1.0 x 10⁻¹ | # | | | | |
| 1.0 x 10⁻¹ | 1.0 x 10⁻¹ | # | | | |
| 1.0 x 10⁻¹ | 1.0 x 10⁻¹ | 1.0 x 10⁻¹ | # | | |
| 1.0 x 10⁻¹ | 1.0 x 10⁻¹ | 1.0 x 10⁻¹ | 1.0 x 10⁻¹ | # | |
| 1.0 x 10⁻¹ | 1.0 x 10⁻¹ | 1.0 x 10⁻¹ | 1.0 x 10⁻¹ | 1.0 x 10⁻¹ | # |

| CD56 Bright NK | Classical Monocyte | Intermediate Monocyte | Non-classical Monocyte | Basophil | Eosinophil | Neutrophil |
|---|---|---|---|---|---|---|
| $1.0 \times 10^{-1}$ | $1.0 \times 10^{-1}$ | $1.0 \times 10^{-1}$ | $1.0 \times 10^{-1}$ | $1.0 \times 10^{-1}$ | $1.0 \times 10^{-1}$ | # |

| | CD56 Bright NK | Classical Monocyte | Intermediate Monocyte | Non-classical Monocyte | Basophil | Eosinophil | Neutrophil |
|---|---|---|---|---|---|---|---|
| | # | | | | | | |
| | $1.0 \times 10^{-1}$ | # | | | | | |
| | $4.2 \times 10^{-1}$ | $3.0 \times 10^{-1}$ | # | | | | |
| | $2.2 \times 10^{-2}$ | $1.0 \times 10^{-2}$ | $1.0 \times 10^{-1}$ | # | | | |
| | $1.0 \times 10^{-1}$ | $1.0 \times 10^{-1}$ | $4.5 \times 10^{-1}$ | $2.2 \times 10^{-2}$ | # | | |
| | $4.1 \times 10^{-13}$ | $1.4 \times 10^{-13}$ | $< 1.0 \times 10^{-16}$ | $< 1.0 \times 10^{-16}$ | $1.0 \times 10^{-13}$ | # | |
| | $1.0 \times 10^{-1}$ | $1.0 \times 10^{-1}$ | $1.3 \times 10^{-1}$ | $3.5 \times 10^{-3}$ | $1.0 \times 10^{-1}$ | $4.3 \times 10^{-11}$ | # |

| CD56 Bright NK | Classical Monocyte | Intermediate Monocyte | Non-classical Monocyte | Basophil | Eosinophil | Neutrophil |
|---|---|---|---|---|---|---|
| # | | | | | | |
| $1.8 \times 10^{-14}$ | # | | | | | |
| $5.4 \times 10^{-14}$ | $1.0 \times 10^{-1}$ | # | | | | |
| $5.4 \times 10^{-14}$ | $1.0 \times 10^{-1}$ | $1.0 \times 10^{-1}$ | # | | | |
| $3.7 \times 10^{-8}$ | $9.5 \times 10^{-4}$ | $4.3 \times 10^{-3}$ | $2.8 \times 10^{-3}$ | # | | |
| $6.0 \times 10^{-12}$ | $2.7 \times 10^{-1}$ | $5.2 \times 10^{-1}$ | $4.2 \times 10^{-1}$ | $1.0 \times 10^{-1}$ | # | |
| $< 1.0 \times 10^{-16}$ | $1.8 \times 10^{-1}$ | $7.0 \times 10^{-2}$ | $1.3 \times 10^{-1}$ | $9.4 \times 10^{-10}$ | $8.7 \times 10^{-6}$ | # |

| CD56 Bright NK | Classical Monocyte | Intermediate Monocyte | Non-classical Monocyte | Basophil | Eosinophil | Neutrophil |
|---|---|---|---|---|---|---|
| # | | | | | | |
| $1.3 \times 10^{-3}$ | # | | | | | |
| $3.3 \times 10^{-4}$ | $1.0 \times 10^{-1}$ | # | | | | |
| $3.9 \times 10^{-1}$ | $8.9 \times 10^{-1}$ | $7.1 \times 10^{-1}$ | # | | | |
| $3.1 \times 10^{-14}$ | $2.8 \times 10^{-8}$ | $1.8 \times 10^{-7}$ | $1.1 \times 10^{-12}$ | # | | |
| $< 1.0 \times 10^{-16}$ | $1.8 \times 10^{-9}$ | $1.2 \times 10^{-8}$ | $1.4 \times 10^{-13}$ | $1.0 \times 10^{-1}$ | # | |
| $< 1.0 \times 10^{-16}$ | $< 1.0 \times 10^{-16}$ | $< 1.0 \times 10^{-16}$ | $< 1.0 \times 10^{-16}$ | $< 1.0 \times 10^{-16}$ | $< 1.0 \times 10^{-16}$ | # |

| CD56 Bright NK | Classical Monocyte | Intermediate Monocyte | Non-classical Monocyte | Basophil | Eosinophil | Neutrophil |
|---|---|---|---|---|---|---|
| # | | | | | | |
| $1.0 \times 10^{-1}$ | # | | | | | |
| $1.0 \times 10^{-1}$ | $1.0 \times 10^{-1}$ | # | | | | |
| $1.0 \times 10^{-1}$ | $1.0 \times 10^{-1}$ | $1.0 \times 10^{-1}$ | # | | | |
| $1.4 \times 10^{-1}$ | $8.8 \times 10^{-4}$ | $7.0 \times 10^{-2}$ | $3.5 \times 10^{-2}$ | # | | |
| $6.5 \times 10^{-3}$ | $2.2 \times 10^{-1}$ | $5.6 \times 10^{-3}$ | $2.0 \times 10^{-2}$ | $4.6 \times 10^{-10}$ | # | |
| $1.0 \times 10^{-1}$ | $1.0 \times 10^{-1}$ | $1.0 \times 10^{-1}$ | $1.0 \times 10^{-1}$ | $1.2 \times 10^{-3}$ | $4.5 \times 10^{-1}$ | # |

| | | | | | | |
|---|---|---|---|---|---|---|
| # | | | | | | |
| $1.4 \times 10^{-3}$ | # | | | | | |
| $8.9 \times 10^{-1}$ | $3.3 \times 10^{-1}$ | # | | | | |
| $9.2 \times 10^{-1}$ | $3.2 \times 10^{-1}$ | $1.0 \times 10^{-1}$ | # | | | |
| $5.7 \times 10^{-2}$ | $3.5 \times 10^{-12}$ | $1.3 \times 10^{-5}$ | $3.0 \times 10^{-5}$ | # | | |
| $1.1 \times 10^{-15}$ | $2.9 \times 10^{-9}$ | $9.1 \times 10^{-14}$ | $8.8 \times 10^{-14}$ | $< 1.0 \times 10^{-16}$ | # | |
| $1.0 \times 10^{-1}$ | $2.9 \times 10^{-2}$ | $1.0 \times 10^{-1}$ | $1.0 \times 10^{-1}$ | $6.7 \times 10^{-3}$ | $8.2 \times 10^{-14}$ | # |

| CD56 Bright NK | Classical Monocyte | Intermediate Monocyte | Non-classical Monocyte | Basophil | Eosinophil | Neutrophil |
|---|---|---|---|---|---|---|
| # | | | | | | |
| $4.8 \times 10^{-14}$ | # | | | | | |
| $7.2 \times 10^{-14}$ | $1.0 \times 10^{-1}$ | # | | | | |
| $4.7 \times 10^{-3}$ | $2.7 \times 10^{-8}$ | $1.7 \times 10^{-7}$ | # | | | |
| $< 1.0 \times 10^{-16}$ | $< 1.0 \times 10^{-16}$ | $< 1.0 \times 10^{-16}$ | $< 1.0 \times 10^{-16}$ | # | | |
| $< 1.0 \times 10^{-16}$ | $< 1.0 \times 10^{-16}$ | $< 1.0 \times 10^{-16}$ | $< 1.0 \times 10^{-16}$ | $9.1 \times 10^{-14}$ | # | |
| $< 1.0 \times 10^{-16}$ | $< 1.0 \times 10^{-16}$ | $< 1.0 \times 10^{-16}$ | $< 1.0 \times 10^{-16}$ | $< 1.0 \times 10^{-16}$ | $< 1.0 \times 10^{-16}$ | # |
| CD56 Bright NK | Classical Monocyte | Intermediate Monocyte | Non-classical Monocyte | Basophil | Eosinophil | Neutrophil |

| CD56 Bright NK | Classical Monocyte | Intermediate Monocyte | Non-classical Monocyte | Basophil | Eosinophil | Neutrophil |
|---|---|---|---|---|---|---|
| # | | | | | | |
| $2.1 \times 10^{-1}$ | # | | | | | |
| $4.9 \times 10^{-1}$ | $1.0 \times 10^{-1}$ | # | | | | |
| $3.1 \times 10^{-1}$ | $1.0 \times 10^{-1}$ | $1.0 \times 10^{-1}$ | # | | | |
| $6.4 \times 10^{-2}$ | $5.1 \times 10^{-8}$ | $6.2 \times 10^{-7}$ | $2.3 \times 10^{-7}$ | # | | |
| $5.8 \times 10^{-9}$ | $3.4 \times 10^{-3}$ | $5.3 \times 10^{-4}$ | $2.5 \times 10^{-3}$ | $5.1 \times 10^{-14}$ | # | |
| $1.0 \times 10^{-1}$ | $1.0 \times 10^{-1}$ | $1.0 \times 10^{-1}$ | $1.0 \times 10^{-1}$ | $3.0 \times 10^{-4}$ | $3.4 \times 10^{-5}$ | # |

| CD56 Bright NK | Classical Monocyte | Intermediate Monocyte | Non-classical Monocyte | Basophil | Eosinophil | Neutrophil |
|---|---|---|---|---|---|---|
| # | | | | | | |
| $1.0 \times 10^{-1}$ | # | | | | | |
| $1.0 \times 10^{-1}$ | $1.0 \times 10^{-1}$ | # | | | | |
| $1.0 \times 10^{-1}$ | $1.0 \times 10^{-1}$ | $1.0 \times 10^{-1}$ | # | | | |
| $1.0 \times 10^{-1}$ | $1.0 \times 10^{-1}$ | $1.0 \times 10^{-1}$ | $1.0 \times 10^{-1}$ | # | | |
| $1.0 \times 10^{-1}$ | $1.0 \times 10^{-1}$ | $1.0 \times 10^{-1}$ | $1.0 \times 10^{-1}$ | $1.0 \times 10^{-1}$ | # | |
| $1.0 \times 10^{-1}$ | $1.0 \times 10^{-1}$ | $1.0 \times 10^{-1}$ | $1.0 \times 10^{-1}$ | $1.0 \times 10^{-1}$ | $1.0 \times 10^{-1}$ | # |

| | | | | | | |
|---|---|---|---|---|---|---|
| # | | | | | | |
| 1.0 x $10^{-1}$ | # | | | | | |
| 1.0 x $10^{-1}$ | 1.0 x $10^{-1}$ | # | | | | |
| 1.0 x $10^{-1}$ | 1.0 x $10^{-1}$ | 1.0 x $10^{-1}$ | # | | | |
| 1.0 x $10^{-1}$ | 1.0 x $10^{-1}$ | 1.0 x $10^{-1}$ | 1.0 x $10^{-1}$ | # | | |
| 1.0 x $10^{-1}$ | 1.0 x $10^{-1}$ | 1.0 x $10^{-1}$ | 1.0 x $10^{-1}$ | 1.0 x $10^{-1}$ | # | |
| 1.0 x $10^{-1}$ | 1.0 x $10^{-1}$ | 1.0 x $10^{-1}$ | 1.0 x $10^{-1}$ | 1.0 x $10^{-1}$ | 1.0 x $10^{-1}$ | # |
| CD56 Bright NK | Classical Monocyte | Intermediate Monocyte | Non-classical Monocyte | Basophil | Eosinophil | Neutrophil |

| | | | | | | |
|---|---|---|---|---|---|---|
| # | | | | | | |
| 1.0 x $10^{-1}$ | # | | | | | |
| 1.0 x $10^{-1}$ | 1.0 x $10^{-1}$ | # | | | | |
| 1.0 x $10^{-1}$ | 1.0 x $10^{-1}$ | 1.0 x $10^{-1}$ | # | | | |
| 1.0 x $10^{-1}$ | 1.0 x $10^{-1}$ | 1.0 x $10^{-1}$ | 1.0 x $10^{-1}$ | # | | |
| 1.0 x $10^{-1}$ | 1.0 x $10^{-1}$ | 1.0 x $10^{-1}$ | 1.0 x $10^{-1}$ | 1.0 x $10^{-1}$ | # | |
| 1.0 x $10^{-1}$ | 1.0 x $10^{-1}$ | 1.0 x $10^{-1}$ | 1.0 x $10^{-1}$ | 1.0 x $10^{-1}$ | 1.0 x $10^{-1}$ | # |

| | | | | | | | | |
|---|---|---|---|---|---|---|---|---|
| CD56 Bright NK | $1.0 \times 10^{-1}$ | $1.0 \times 10^{-1}$ | $1.0 \times 10^{-1}$ | $1.0 \times 10^{-1}$ | $1.0 \times 10^{-1}$ | $1.0 \times 10^{-1}$ | # | CD56 Bright NK |
| Classical Monocyte | $1.0 \times 10^{-1}$ | $1.0 \times 10^{-1}$ | $1.0 \times 10^{-1}$ | $1.0 \times 10^{-1}$ | $1.0 \times 10^{-1}$ | # | | Classical Monocyte |
| Intermediate Monocyte | $1.0 \times 10^{-1}$ | $1.0 \times 10^{-1}$ | $1.0 \times 10^{-1}$ | $1.0 \times 10^{-1}$ | # | | | Intermediate Monocyte |
| Non-classical Monocyte | $1.0 \times 10^{-1}$ | $1.0 \times 10^{-1}$ | $1.0 \times 10^{-1}$ | # | | | | Non-classical Monocyte |
| Basophil | $1.0 \times 10^{-1}$ | $1.0 \times 10^{-1}$ | # | | | | | Basophil |
| Eosinophil | $1.0 \times 10^{-1}$ | # | | | | | | Eosinophil |
| Neutrophil | # | | | | | | | Neutrophil |

| CD56 Bright NK | Classical Monocyte | Intermediate Monocyte | Non-classical Monocyte | Basophil | Eosinophil | Neutrophil |
|---|---|---|---|---|---|---|
| # | | | | | | |
| $1.0 \times 10^{-1}$ | # | | | | | |
| $6.0 \times 10^{-1}$ | $1.0 \times 10^{-1}$ | # | | | | |
| $3.1 \times 10^{-2}$ | $5.4 \times 10^{-1}$ | $1.0 \times 10^{-1}$ | # | | | |
| $3.8 \times 10^{-5}$ | $9.3 \times 10^{-9}$ | $2.0 \times 10^{-11}$ | $9.0 \times 10^{-14}$ | # | | |
| $1.7 \times 10^{-7}$ | $4.5 \times 10^{-5}$ | $3.7 \times 10^{-3}$ | $2.8 \times 10^{-1}$ | $< 1.0 \times 10^{-16}$ | # | |
| $1.3 \times 10^{-13}$ | $2.0 \times 10^{-11}$ | $6.3 \times 10^{-9}$ | $1.1 \times 10^{-5}$ | $< 1.0 \times 10^{-16}$ | $2.4 \times 10^{-1}$ | # |

| CD56 Bright NK | Classical Monocyte | Intermediate Monocyte | Non-classical Monocyte | Basophil | Eosinophil | Neutrophil |
|---|---|---|---|---|---|---|
| # | | | | | | |
| $1.0 \times 10^{-1}$ | # | | | | | |
| $1.0 \times 10^{-1}$ | $1.0 \times 10^{-1}$ | # | | | | |
| $1.0 \times 10^{-1}$ | $1.0 \times 10^{-1}$ | $1.0 \times 10^{-1}$ | # | | | |
| $1.0 \times 10^{-1}$ | $1.0 \times 10^{-1}$ | $1.0 \times 10^{-1}$ | $1.0 \times 10^{-1}$ | # | | |
| $1.0 \times 10^{-1}$ | $1.0 \times 10^{-1}$ | $1.0 \times 10^{-1}$ | $1.0 \times 10^{-1}$ | $1.0 \times 10^{-1}$ | # | |
| $1.0 \times 10^{-1}$ | $1.0 \times 10^{-1}$ | $1.0 \times 10^{-1}$ | $1.0 \times 10^{-1}$ | $1.0 \times 10^{-1}$ | $1.0 \times 10^{-1}$ | # |

| | | | | | | |
|---|---|---|---|---|---|---|
| # | | | | | | |
| $1.0 \times 10^{-1}$ | # | | | | | |
| $1.0 \times 10^{-1}$ | $1.0 \times 10^{-1}$ | # | | | | |
| $9.4 \times 10^{-1}$ | $5.3 \times 10^{-1}$ | $5.1 \times 10^{-1}$ | # | | | |
| $1.1 \times 10^{-13}$ | $2.2 \times 10^{-13}$ | $2.4 \times 10^{-13}$ | $2.0 \times 10^{-14}$ | # | | |
| $< 1.0 \times 10^{-16}$ | $< 1.0 \times 10^{-16}$ | $< 1.0 \times 10^{-16}$ | $< 1.0 \times 10^{-16}$ | $4.9 \times 10^{-8}$ | # | |
| $< 1.0 \times 10^{-16}$ | $< 1.0 \times 10^{-16}$ | $< 1.0 \times 10^{-16}$ | $< 1.0 \times 10^{-16}$ | $< 1.0 \times 10^{-16}$ | $7.2 \times 10^{-6}$ | # |
| CD56 Bright NK | Classical Monocyte | Intermediate Monocyte | Non-classical Monocyte | Basophil | Eosinophil | Neutrophil |

| | | | | | | |
|---|---|---|---|---|---|---|
| # | | | | | | |
| $1.0 \times 10^{-1}$ | # | | | | | |
| $1.0 \times 10^{-1}$ | $1.0 \times 10^{-1}$ | # | | | | |
| $1.0 \times 10^{-1}$ | $1.0 \times 10^{-1}$ | $1.0 \times 10^{-1}$ | # | | | |
| $1.0 \times 10^{-1}$ | $1.0 \times 10^{-1}$ | $9.4 \times 10^{-1}$ | $8.9 \times 10^{-1}$ | # | | |
| $1.0 \times 10^{-1}$ | $1.0 \times 10^{-1}$ | $1.0 \times 10^{-1}$ | $1.0 \times 10^{-1}$ | $4.0 \times 10^{-1}$ | # | |
| $1.0 \times 10^{-1}$ | $1.0 \times 10^{-1}$ | $1.0 \times 10^{-1}$ | $1.0 \times 10^{-1}$ | $5.5 \times 10^{-1}$ | $1.0 \times 10^{-1}$ | # |

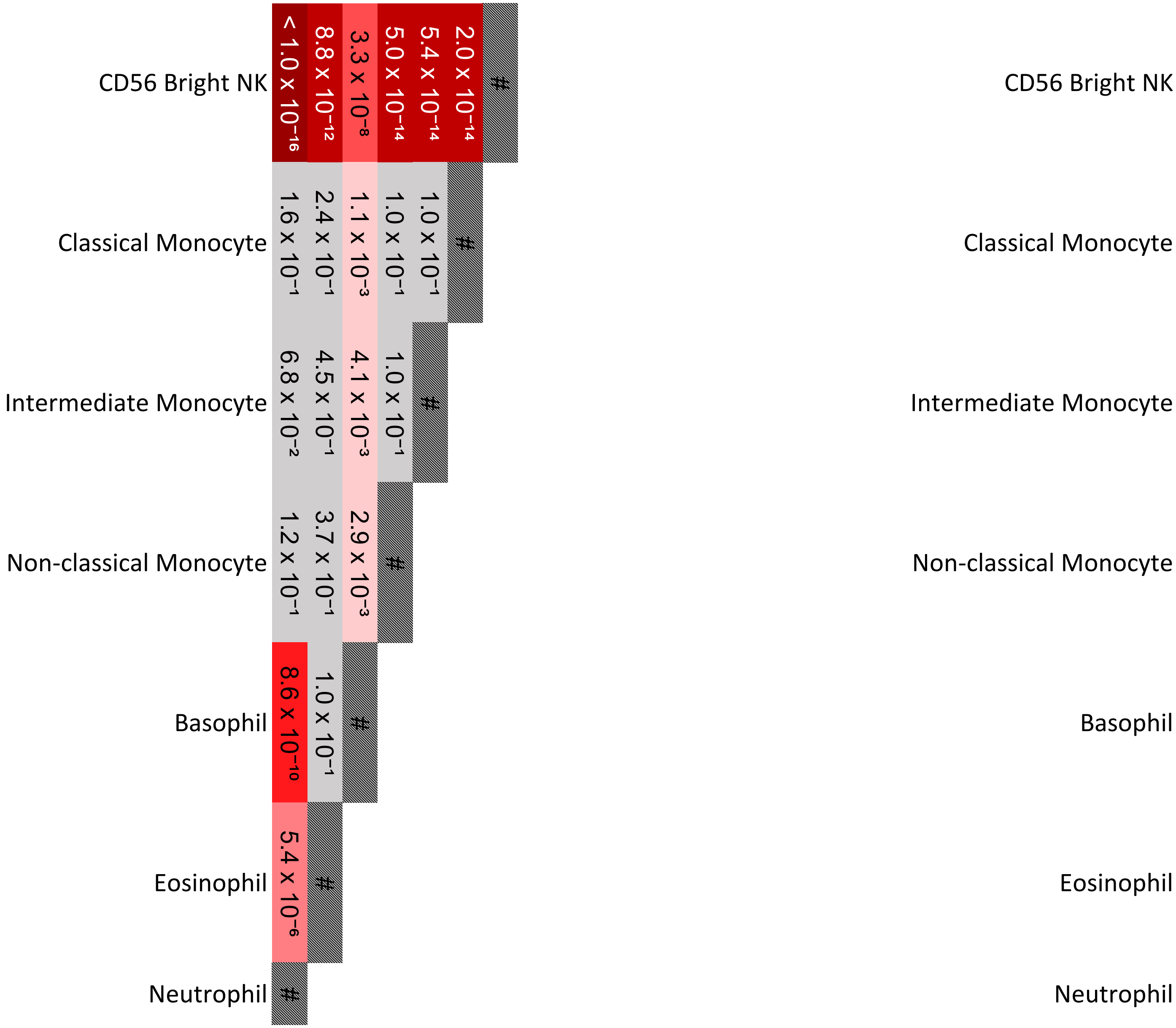
CD56 Bright NK
Classical Monocyte
Intermediate Monocyte
Non-classical Monocyte
Basophil
Eosinophil
Neutrophil
< 1.0 x 10⁻¹⁶
8.8 x 10⁻¹²
3.3 x 10⁻⁸
5.0 x 10⁻¹⁴
5.4 x 10⁻¹⁴
2.0 x 10⁻¹⁴
1.6 x 10⁻¹
2.4 x 10⁻¹
1.1 x 10⁻³
1.0 x 10⁻¹
1.0 x 10⁻¹
6.8 x 10⁻²
4.5 x 10⁻¹
4.1 x 10⁻³
1.0 x 10⁻¹
1.2 x 10⁻¹
3.7 x 10⁻¹
2.9 x 10⁻³
8.6 x 10⁻¹⁰
1.0 x 10⁻¹
5.4 x 10⁻⁶
#
CD56 Bright NK
Classical Monocyte
Intermediate Monocyte
Non-classical Monocyte
Basophil
Eosinophil
Neutrophil

| | CD56 Bright NK | Classical Monocyte | Intermediate Monocyte | Non-classical Monocyte | Basophil | Eosinophil | Neutrophil |
|---|---|---|---|---|---|---|---|
| | # | | | | | | |
| | $8.8 \times 10^{-14}$ | # | | | | | |
| | $1.1 \times 10^{-11}$ | $1.0 \times 10^{-1}$ | # | | | | |
| | $3.5 \times 10^{-5}$ | $1.3 \times 10^{-2}$ | $3.4 \times 10^{-1}$ | # | | | |
| | $< 1.0 \times 10^{-16}$ | $4.3 \times 10^{-5}$ | $9.6 \times 10^{-8}$ | $1.0 \times 10^{-13}$ | # | | |
| | $1.9 \times 10^{-1}$ | $< 1.0 \times 10^{-16}$ | $< 1.0 \times 10^{-16}$ | $9.6 \times 10^{-13}$ | $< 1.0 \times 10^{-16}$ | # | |
| | $< 1.0 \times 10^{-16}$ | $6.7 \times 10^{-5}$ | $2.4 \times 10^{-7}$ | $2.4 \times 10^{-13}$ | $1.0 \times 10^{-1}$ | $< 1.0 \times 10^{-16}$ | # |

| | CD56 Bright NK | Classical Monocyte | Intermediate Monocyte | Non-classical Monocyte | Basophil | Eosinophil | Neutrophil |
|---|---|---|---|---|---|---|---|
| | # | | | | | | |
| | $1.0 \times 10^{-1}$ | # | | | | | |
| | $1.0 \times 10^{-1}$ | $1.0 \times 10^{-1}$ | # | | | | |
| | $1.0 \times 10^{-1}$ | $1.0 \times 10^{-1}$ | $1.0 \times 10^{-1}$ | # | | | |
| | $1.0 \times 10^{-1}$ | $1.0 \times 10^{-1}$ | $1.0 \times 10^{-1}$ | $1.0 \times 10^{-1}$ | # | | |
| | $1.0 \times 10^{-1}$ | $1.0 \times 10^{-1}$ | $1.0 \times 10^{-1}$ | $1.0 \times 10^{-1}$ | $1.0 \times 10^{-1}$ | # | |
| | $1.0 \times 10^{-1}$ | $1.0 \times 10^{-1}$ | $1.0 \times 10^{-1}$ | $1.0 \times 10^{-1}$ | $1.0 \times 10^{-1}$ | $1.0 \times 10^{-1}$ | # |

| CD56 Bright NK | Classical Monocyte | Intermediate Monocyte | Non-classical Monocyte | Basophil | Eosinophil | Neutrophil |
|---|---|---|---|---|---|---|
| # | | | | | | |
| $2.9 \times 10^{-3}$ | # | | | | | |
| $2.0 \times 10^{-2}$ | $1.0 \times 10^{-1}$ | # | | | | |
| $2.1 \times 10^{-4}$ | $1.0 \times 10^{-1}$ | $1.0 \times 10^{-1}$ | # | | | |
| $2.6 \times 10^{-4}$ | $9.3 \times 10^{-14}$ | $1.2 \times 10^{-13}$ | $7.8 \times 10^{-14}$ | # | | |
| $5.0 \times 10^{-1}$ | $6.8 \times 10^{-9}$ | $1.3 \times 10^{-7}$ | $2.2 \times 10^{-10}$ | $5.4 \times 10^{-1}$ | # | |
| $6.5 \times 10^{-4}$ | $1.1 \times 10^{-13}$ | $7.4 \times 10^{-13}$ | $9.0 \times 10^{-14}$ | $1.0 \times 10^{-1}$ | $6.0 \times 10^{-1}$ | # |

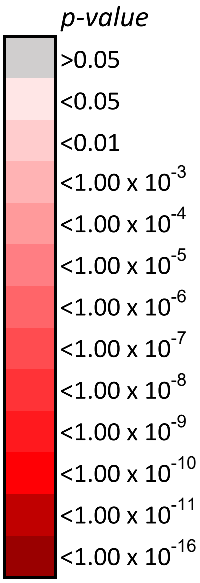
*p-value*
>0.05
<0.05
<0.01
<1.00 x $10^{-3}$
<1.00 x $10^{-4}$
<1.00 x $10^{-5}$
<1.00 x $10^{-6}$
<1.00 x $10^{-7}$
<1.00 x $10^{-8}$
<1.00 x $10^{-9}$
<1.00 x $10^{-10}$
<1.00 x $10^{-11}$
<1.00 x $10^{-16}$

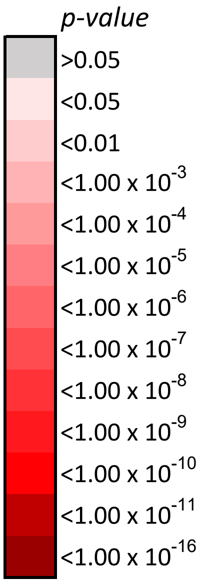
*p-value*
>0.05
<0.05
<0.01
<1.00 x $10^{-3}$
<1.00 x $10^{-4}$
<1.00 x $10^{-5}$
<1.00 x $10^{-6}$
<1.00 x $10^{-7}$
<1.00 x $10^{-8}$
<1.00 x $10^{-9}$
<1.00 x $10^{-10}$
<1.00 x $10^{-11}$
<1.00 x $10^{-16}$

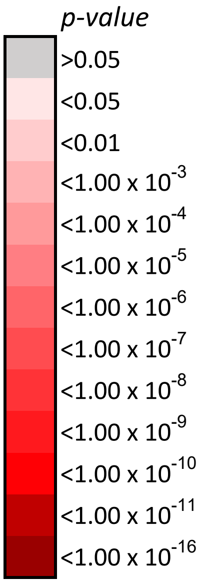
p-value
>0.05
<0.05
<0.01
<1.00 x 10-3
<1.00 x 10-4
<1.00 x 10-5
<1.00 x 10-6
<1.00 x 10-7
<1.00 x 10-8
<1.00 x 10-9
<1.00 x 10-10
<1.00 x 10-11
<1.00 x 10-16

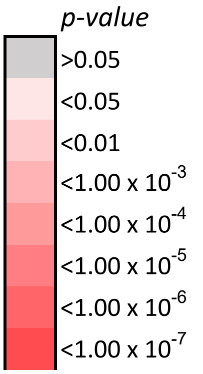
p-value
>0.05
<0.05
<0.01
<1.00 x 10-3
<1.00 x 10-4
<1.00 x 10-5
<1.00 x 10-6
<1.00 x 10-7

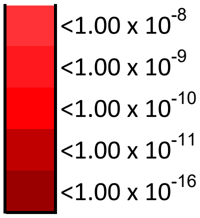

<1.00 x 10-8
<1.00 x 10-9
<1.00 x 10-10
<1.00 x 10-11
<1.00 x 10-16


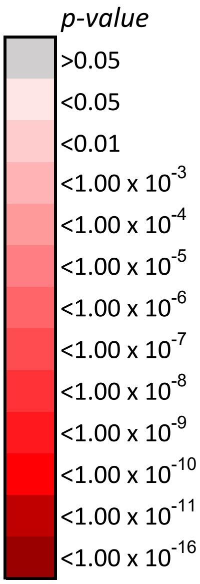

p-value
>0.05
<0.05
<0.01
<1.00 x 10-3
<1.00 x 10-4
<1.00 x 10-5
<1.00 x 10-6
<1.00 x 10-7
<1.00 x 10-8
<1.00 x 10-9
<1.00 x 10-10
<1.00 x 10-11
<1.00 x 10-16


*p-value*

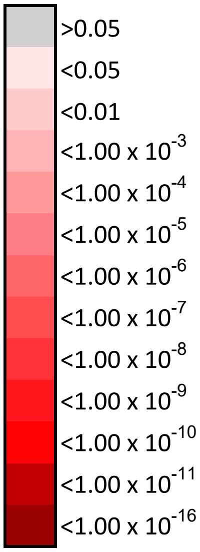
>0.05
<0.05
<0.01
<1.00 x 10-3
<1.00 x 10-4
<1.00 x 10-5
<1.00 x 10-6
<1.00 x 10-7
<1.00 x 10-8
<1.00 x 10-9
<1.00 x 10-10
<1.00 x 10-11
<1.00 x 10-16

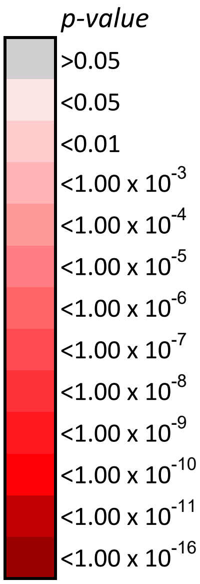
p-value
>0.05
<0.05
<0.01
<1.00 x 10-3
<1.00 x 10-4
<1.00 x 10-5
<1.00 x 10-6
<1.00 x 10-7
<1.00 x 10-8
<1.00 x 10-9
<1.00 x 10-10
<1.00 x 10-11
<1.00 x 10-16

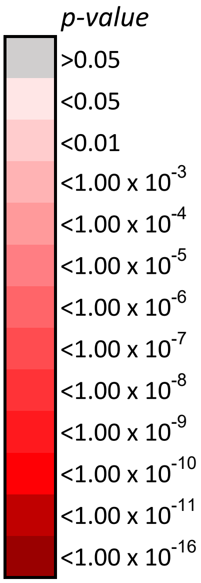
p-value
>0.05
<0.05
<0.01
<1.00 x 10-3
<1.00 x 10-4
<1.00 x 10-5
<1.00 x 10-6
<1.00 x 10-7
<1.00 x 10-8
<1.00 x 10-9
<1.00 x 10-10
<1.00 x 10-11
<1.00 x 10-16

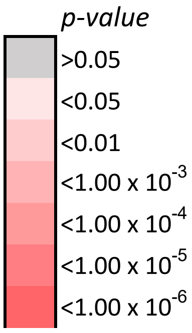
p-value
>0.05
<0.05
<0.01
<1.00 x 10-3
<1.00 x 10-4
<1.00 x 10-5
<1.00 x 10-6

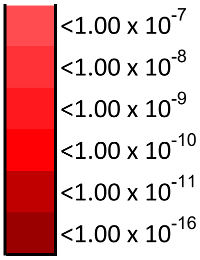

<1.00 x 10-7
<1.00 x 10-8
<1.00 x 10-9
<1.00 x 10-10
<1.00 x 10-11
<1.00 x 10-16


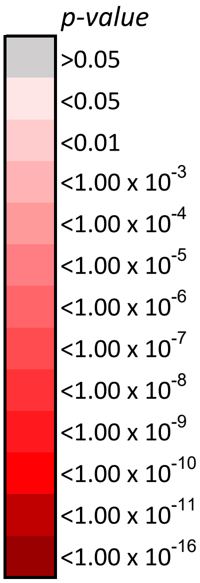

p-value
>0.05
<0.05
<0.01
<1.00 x 10-3
<1.00 x 10-4
<1.00 x 10-5
<1.00 x 10-6
<1.00 x 10-7
<1.00 x 10-8
<1.00 x 10-9
<1.00 x 10-10
<1.00 x 10-11
<1.00 x 10-16

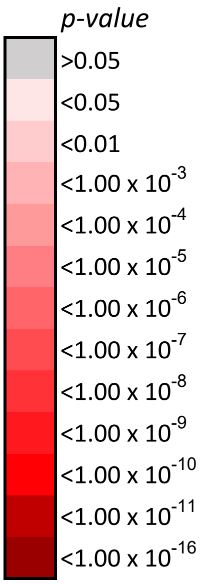
p-value
>0.05
<0.05
<0.01
<1.00 x 10-3
<1.00 x 10-4
<1.00 x 10-5
<1.00 x 10-6
<1.00 x 10-7
<1.00 x 10-8
<1.00 x 10-9
<1.00 x 10-10
<1.00 x 10-11
<1.00 x 10-16

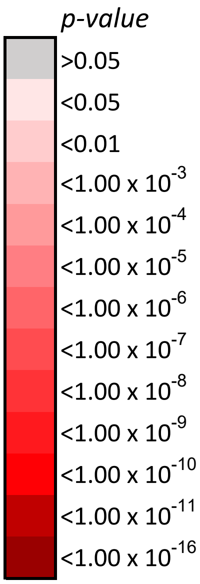
p-value
>0.05
<0.05
<0.01
<1.00 x 10-3
<1.00 x 10-4
<1.00 x 10-5
<1.00 x 10-6
<1.00 x 10-7
<1.00 x 10-8
<1.00 x 10-9
<1.00 x 10-10
<1.00 x 10-11
<1.00 x 10-16

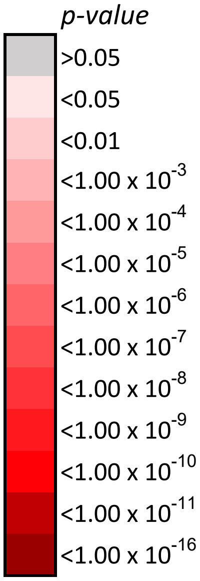
p-value
>0.05
<0.05
<0.01
<1.00 x 10-3
<1.00 x 10-4
<1.00 x 10-5
<1.00 x 10-6
<1.00 x 10-7
<1.00 x 10-8
<1.00 x 10-9
<1.00 x 10-10
<1.00 x 10-11
<1.00 x 10-16

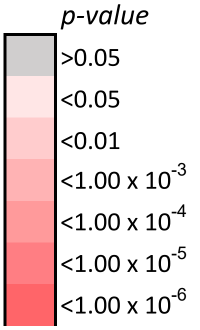
p-value
>0.05
<0.05
<0.01
<1.00 x 10-3
<1.00 x 10-4
<1.00 x 10-5
<1.00 x 10-6

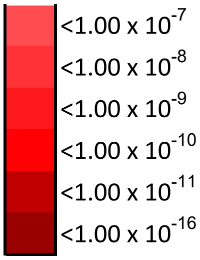

<1.00 x 10$^{-7}$
<1.00 x 10$^{-8}$
<1.00 x 10$^{-9}$
<1.00 x 10$^{-10}$
<1.00 x 10$^{-11}$
<1.00 x 10$^{-16}$


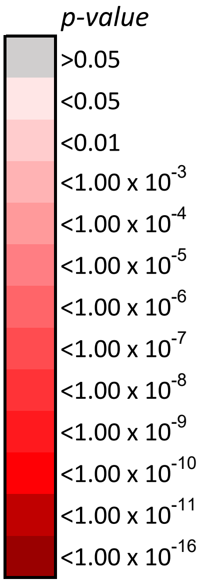

*p-value*
>0.05
<0.05
<0.01
<1.00 x 10$^{-3}$
<1.00 x 10$^{-4}$
<1.00 x 10$^{-5}$
<1.00 x 10$^{-6}$
<1.00 x 10$^{-7}$
<1.00 x 10$^{-8}$
<1.00 x 10$^{-9}$
<1.00 x 10$^{-10}$
<1.00 x 10$^{-11}$
<1.00 x 10$^{-16}$

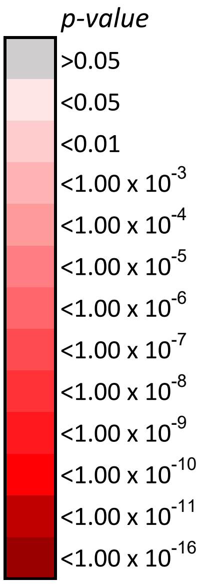
p-value
>0.05
<0.05
<0.01
<1.00 x 10-3
<1.00 x 10-4
<1.00 x 10-5
<1.00 x 10-6
<1.00 x 10-7
<1.00 x 10-8
<1.00 x 10-9
<1.00 x 10-10
<1.00 x 10-11
<1.00 x 10-16

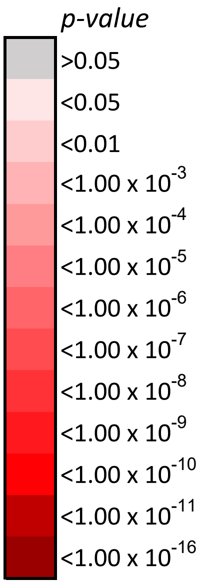
p-value
>0.05
<0.05
<0.01
<1.00 x 10-3
<1.00 x 10-4
<1.00 x 10-5
<1.00 x 10-6
<1.00 x 10-7
<1.00 x 10-8
<1.00 x 10-9
<1.00 x 10-10
<1.00 x 10-11
<1.00 x 10-16

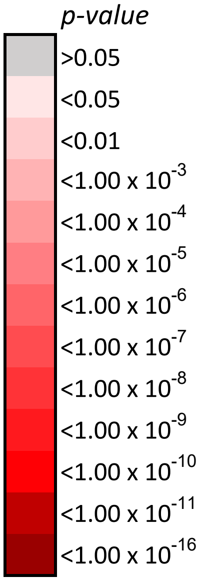
p-value
>0.05
<0.05
<0.01
<1.00 x 10-3
<1.00 x 10-4
<1.00 x 10-5
<1.00 x 10-6
<1.00 x 10-7
<1.00 x 10-8
<1.00 x 10-9
<1.00 x 10-10
<1.00 x 10-11
<1.00 x 10-16

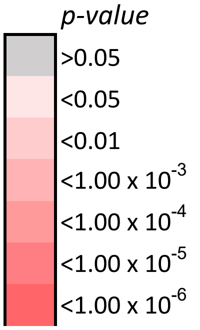
p-value
>0.05
<0.05
<0.01
<1.00 x 10-3
<1.00 x 10-4
<1.00 x 10-5
<1.00 x 10-6

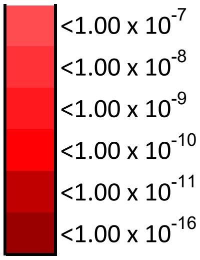
<1.00 x 10-7
<1.00 x 10-8
<1.00 x 10-9
<1.00 x 10-10
<1.00 x 10-11
<1.00 x 10-16

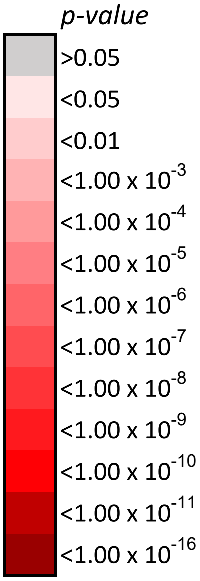
p-value
>0.05
<0.05
<0.01
<1.00 x 10-3
<1.00 x 10-4
<1.00 x 10-5
<1.00 x 10-6
<1.00 x 10-7
<1.00 x 10-8
<1.00 x 10-9
<1.00 x 10-10
<1.00 x 10-11
<1.00 x 10-16

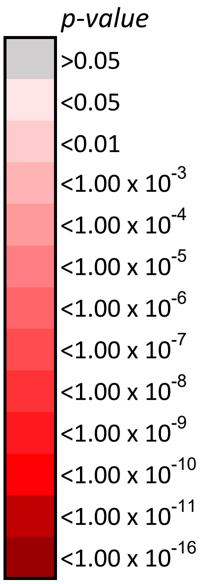

p-value
>0.05
<0.05
<0.01
<1.00 x 10^-3
<1.00 x 10^-4
<1.00 x 10^-5
<1.00 x 10^-6
<1.00 x 10^-7
<1.00 x 10^-8
<1.00 x 10^-9
<1.00 x 10^-10
<1.00 x 10^-11
<1.00 x 10^-16
