## Supplementary material for "An immune cell lipid atlas reveals the basis of susceptibility to ferroptosis": Mouse statistics

### One-way ANOVA results of global phos[illegible]

| Lipid class | PL acyl chains |
|---|---|
| Feature | 14:0 |
| One-way ANOVA *p-value* | $1.3 \times 10^{-11}$ |
| FDR adjuested *p-value* | $2.5 \times 10^{-11}$ |

| | |
|---|---|
| B Cell | # |
| CD4 T Cell | $1.0 \times 10^{-1}$ |
| CD8 T Cell | $2.0 \times 10^{-1}$ |
| NK Cell | $1.7 \times 10^{-4}$ |
| Ly6C High Monocyte | $2.4 \times 10^{-4}$ |
| Ly6C Low Monocyte | $2.2 \times 10^{-4}$ |
| Eosinophil | $1.8 \times 10^{-5}$ |
| Neutrophil | $1.9 \times 10^{-8}$ |
| | B Cell |

| Lipid class | PL acyl chains |
|---|---|
| Feature | 15:0 |
| One-way ANOVA *p-value* | $1.6 \times 10^{-40}$ |
| FDR adjuested *p-value* | $3.3 \times 10^{-39}$ |

| | |
|---|---|
| B Cell | # |
| CD4 T Cell | $7.2 \times 10^{-1}$ |
| CD8 T Cell | $8.1 \times 10^{-1}$ |
| NK Cell | $1.0 \times 10^{-1}$ |
| Ly6C High Monocyte | $2.9 \times 10^{-1}$ |
| Ly6C Low Monocyte | $1.0 \times 10^{-1}$ |
| Eosinophil | $6.6 \times 10^{-9}$ |
| Neutrophil | $9.5 \times 10^{-12}$ |
| | B Cell |

| Lipid class | PL acyl chains |
|---|---|
| Feature | 16:0 |
| One-way ANOVA *p-value* | $5.8 \times 10^{-8}$ |
| FDR adjuested *p-value* | $6.8 \times 10^{-8}$ |

| | |
|---|---|
| B Cell | # |
| CD4 T Cell | $1.0 \times 10^{-1}$ |
| CD8 T Cell | $1.6 \times 10^{-1}$ |
| NK Cell | $6.8 \times 10^{-3}$ |
| Ly6C High Monocyte | $1.1 \times 10^{-1}$ |
| Ly6C Low Monocyte | $2.7 \times 10^{-1}$ |
| Eosinophil | $6.8 \times 10^{-1}$ |

| | |
|---|---|
| Neutrophil | 1.6 x $10^{-1}$ |
| | B Cell |

| | |
|---|---|
| Lipid class | **PL acyl chains** |
| Feature | 16:1 |
| One-way ANOVA *p-value* | 8.7 x $10^{-10}$ |
| FDR adjuested *p-value* | 1.3 x $10^{-9}$ |

| | |
|---|---|
| B Cell | # |
| CD4 T Cell | 1.0 x $10^{-1}$ |
| CD8 T Cell | 4.4 x $10^{-1}$ |
| NK Cell | 4.5 x $10^{-2}$ |
| Ly6C High Monocyte | 6.6 x $10^{-1}$ |
| Ly6C Low Monocyte | 6.8 x $10^{-2}$ |
| Eosinophil | 8.8 x $10^{-1}$ |
| Neutrophil | 2.1 x $10^{-2}$ |
| | B Cell |

| | |
|---|---|
| Lipid class | **PL acyl chains** |
| Feature | 17:0 |
| One-way ANOVA *p-value* | 8.9 x $10^{-3}$ |
| FDR adjuested *p-value* | 8.9 x $10^{-3}$ |

| | |
|---|---|
| B Cell | # |
| CD4 T Cell | 9.3 x $10^{-1}$ |
| CD8 T Cell | 7.3 x $10^{-1}$ |
| NK Cell | 9.4 x $10^{-1}$ |
| Ly6C High Monocyte | 4.1 x $10^{-1}$ |
| Ly6C Low Monocyte | 1.3 x $10^{-2}$ |
| Eosinophil | 1.8 x $10^{-2}$ |
| Neutrophil | 6.6 x $10^{-1}$ |
| | B Cell |

Lipid class **PL acyl chains**

| Feature | 17:1 |
| --- | --- |
| One-way ANOVA *p-value* | $4.2 \times 10^{-8}$ |
| FDR adjuested *p-value* | $5.2 \times 10^{-8}$ |

| | B Cell |
| --- | --- |
| B Cell | # |
| CD4 T Cell | $3.4 \times 10^{-2}$ |
| CD8 T Cell | $1.2 \times 10^{-1}$ |
| NK Cell | $1.4 \times 10^{-1}$ |
| Ly6C High Monocyte | $8.3 \times 10^{-1}$ |
| Ly6C Low Monocyte | $1.0 \times 10^{-1}$ |
| Eosinophil | $1.9 \times 10^{-3}$ |
| Neutrophil | $1.6 \times 10^{-2}$ |

| Lipid class | **PL acyl chains** |
| --- | --- |
| Feature | 18:0 |
| One-way ANOVA *p-value* | $3.8 \times 10^{-9}$ |
| FDR adjuested *p-value* | $5.0 \times 10^{-9}$ |

| | B Cell |
| --- | --- |
| B Cell | # |
| CD4 T Cell | $1.0 \times 10^{-1}$ |
| CD8 T Cell | $2.7 \times 10^{-1}$ |
| NK Cell | $7.0 \times 10^{-2}$ |
| Ly6C High Monocyte | $2.3 \times 10^{-1}$ |
| Ly6C Low Monocyte | $1.0 \times 10^{-1}$ |
| Eosinophil | $3.6 \times 10^{-4}$ |
| Neutrophil | $1.9 \times 10^{-1}$ |

| Lipid class | **PL acyl chains** |
| --- | --- |
| Feature | 18:1 |
| One-way ANOVA *p-value* | $5.0 \times 10^{-26}$ |
| FDR adjuested *p-value* | $1.8 \times 10^{-25}$ |

| | B Cell |
| --- | --- |
| B Cell | # |
| CD4 T Cell | $6.8 \times 10^{-10}$ |
| CD8 T Cell | $1.9 \times 10^{-8}$ |
| NK Cell | $2.5 \times 10^{-7}$ |
| Ly6C High Monocyte | $9.5 \times 10^{-12}$ |
| Ly6C Low Monocyte | $9.5 \times 10^{-12}$ |
| Eosinophil | $9.5 \times 10^{-12}$ |
| Neutrophil | $9.5 \times 10^{-12}$ |

B Cell

| Lipid class | **PL acyl chains** |
|---|---|
| Feature | 18:2 |
| One-way ANOVA *p-value* | 1.9 x $10^{-9}$ |
| FDR adjuested *p-value* | 2.7 x $10^{-9}$ |

| | |
|---|---|
| B Cell | # |
| CD4 T Cell | 2.8 x $10^{-1}$ |
| CD8 T Cell | 5.9 x $10^{-2}$ |
| NK Cell | 7.8 x $10^{-1}$ |
| Ly6C High Monocyte | 1.2 x $10^{-1}$ |
| Ly6C Low Monocyte | 5.1 x $10^{-1}$ |
| Eosinophil | 1.3 x $10^{-2}$ |
| Neutrophil | 1.0 x $10^{-1}$ |

B Cell

| Lipid class | **PL acyl chains** |
|---|---|
| Feature | 18:3 |
| One-way ANOVA *p-value* | 8.0 x $10^{-28}$ |
| FDR adjuested *p-value* | 4.2 x $10^{-27}$ |

| | |
|---|---|
| B Cell | # |
| CD4 T Cell | 1.0 x $10^{-1}$ |
| CD8 T Cell | 1.0 x $10^{-1}$ |
| NK Cell | 1.0 x $10^{-1}$ |
| Ly6C High Monocyte | 1.5 x $10^{-1}$ |
| Ly6C Low Monocyte | 1.0 x $10^{-1}$ |
| Eosinophil | 1.0 x $10^{-1}$ |
| Neutrophil | 9.5 x $10^{-12}$ |

B Cell

| Lipid class | **PL acyl chains** |
|---|---|
| Feature | 19:0 |

| | |
|---|---|
| B Cell | # |

| | |
|---|---|
| One-way ANOVA *p-value* | 2.1 x 10⁻¹³ |
| FDR adjuested *p-value* | 4.4 x 10⁻¹³ |

| | B Cell |
|---|---|
| CD4 T Cell | 1.0 x 10⁻¹ |
| CD8 T Cell | 9.0 x 10⁻¹ |
| NK Cell | 1.0 x 10⁻¹ |
| Ly6C High Monocyte | 1.0 x 10⁻¹ |
| Ly6C Low Monocyte | 4.1 x 10⁻² |
| Eosinophil | 1.5 x 10⁻³ |
| Neutrophil | 5.4 x 10⁻¹¹ |

| | |
|---|---|
| Lipid class | **PL acyl chains** |
| Feature | 20:0 |
| One-way ANOVA *p-value* | 5.6 x 10⁻¹¹ |
| FDR adjuested *p-value* | 9.1 x 10⁻¹¹ |

| | B Cell |
|---|---|
| B Cell | # |
| CD4 T Cell | 1.0 x 10⁻¹ |
| CD8 T Cell | 8.2 x 10⁻¹ |
| NK Cell | 1.0 x 10⁻¹ |
| Ly6C High Monocyte | 1.0 x 10⁻¹ |
| Ly6C Low Monocyte | 1.0 x 10⁻¹ |
| Eosinophil | 2.2 x 10⁻² |
| Neutrophil | 3.2 x 10⁻⁸ |

| | |
|---|---|
| Lipid class | **PL acyl chains** |
| Feature | 20:1 |
| One-way ANOVA *p-value* | 5.1 x 10⁻¹⁹ |
| FDR adjuested *p-value* | 1.3 x 10⁻¹⁸ |

| | |
|---|---|
| B Cell | # |
| CD4 T Cell | 2.7 x 10⁻⁷ |
| CD8 T Cell | 4.5 x 10⁻⁸ |
| NK Cell | 1.1 x 10⁻⁶ |
| Ly6C High Monocyte | 8.5 x 10⁻¹ |
| Ly6C Low Monocyte | 6.4 x 10⁻⁵ |
| Eosinophil | 1.0 x 10⁻¹ |
| Neutrophil | 3.4 x 10⁻¹ |

B Cell

| Lipid class | **PL acyl chains** |
|---|---|
| Feature | 20:2 |
| One-way ANOVA *p-value* | $1.5 \times 10^{-7}$ |
| FDR adjuested *p-value* | $1.6 \times 10^{-7}$ |

| | |
|---|---|
| B Cell | # |
| CD4 T Cell | $1.0 \times 10^{-1}$ |
| CD8 T Cell | $5.7 \times 10^{-2}$ |
| NK Cell | $1.6 \times 10^{-1}$ |
| Ly6C High Monocyte | $1.3 \times 10^{-1}$ |
| Ly6C Low Monocyte | $2.1 \times 10^{-1}$ |
| Eosinophil | $1.0 \times 10^{-1}$ |
| Neutrophil | $1.4 \times 10^{-1}$ |

B Cell

| Lipid class | **PL acyl chains** |
|---|---|
| Feature | 20:3 |
| One-way ANOVA *p-value* | $7.9 \times 10^{-6}$ |
| FDR adjuested *p-value* | $8.3 \times 10^{-6}$ |

| | |
|---|---|
| B Cell | # |
| CD4 T Cell | $1.5 \times 10^{-1}$ |
| CD8 T Cell | $1.8 \times 10^{-1}$ |
| NK Cell | $2.8 \times 10^{-1}$ |
| Ly6C High Monocyte | $4.4 \times 10^{-3}$ |
| Ly6C Low Monocyte | $6.8 \times 10^{-7}$ |
| Eosinophil | $2.6 \times 10^{-1}$ |
| Neutrophil | $3.0 \times 10^{-1}$ |

B Cell

| Lipid class | **PL acyl chains** |
|---|---|
| Feature | 20:4 |

| | |
|---|---|
| B Cell | # |

| | |
|---|---|
| One-way ANOVA *p-value* | $1.8 \times 10^{-17}$ |
| FDR adjuested *p-value* | $4.1 \times 10^{-17}$ |

| | B Cell |
|---|---|
| CD4 T Cell | $1.0 \times 10^{-1}$ |
| CD8 T Cell | $1.0 \times 10^{-1}$ |
| NK Cell | $1.7 \times 10^{-1}$ |
| Ly6C High Monocyte | $2.6 \times 10^{-1}$ |
| Ly6C Low Monocyte | $1.0 \times 10^{-1}$ |
| Eosinophil | $2.4 \times 10^{-1}$ |
| Neutrophil | $7.6 \times 10^{-11}$ |

| | |
|---|---|
| Lipid class | **PL acyl chains** |
| Feature | 20:5 |
| One-way ANOVA *p-value* | $3.2 \times 10^{-38}$ |
| FDR adjuested *p-value* | $3.4 \times 10^{-37}$ |

| | B Cell |
|---|---|
| B Cell | # |
| CD4 T Cell | $1.5 \times 10^{-11}$ |
| CD8 T Cell | $2.6 \times 10^{-9}$ |
| NK Cell | $1.0 \times 10^{-1}$ |
| Ly6C High Monocyte | $1.0 \times 10^{-11}$ |
| Ly6C Low Monocyte | $9.4 \times 10^{-4}$ |
| Eosinophil | $9.5 \times 10^{-12}$ |
| Neutrophil | $5.3 \times 10^{-1}$ |

| | |
|---|---|
| Lipid class | **PL acyl chains** |
| Feature | 22:2 |
| One-way ANOVA *p-value* | $1.4 \times 10^{-11}$ |
| FDR adjuested *p-value* | $2.5 \times 10^{-11}$ |

| | |
|---|---|
| B Cell | # |
| CD4 T Cell | $8.7 \times 10^{-4}$ |
| CD8 T Cell | $2.1 \times 10^{-3}$ |
| NK Cell | $1.0 \times 10^{-1}$ |
| Ly6C High Monocyte | $8.6 \times 10^{-3}$ |
| Ly6C Low Monocyte | $1.6 \times 10^{-1}$ |
| Eosinophil | $1.0 \times 10^{-1}$ |
| Neutrophil | $5.4 \times 10^{-2}$ |

B Cell

| Lipid class | **PL acyl chains** |
|---|---|
| Feature | 22:4 |
| One-way ANOVA *p-value* | $6.6 \times 10^{-22}$ |
| FDR adjuested *p-value* | $2.0 \times 10^{-21}$ |

| | |
|---|---|
| B Cell | # |
| CD4 T Cell | $1.0 \times 10^{-1}$ |
| CD8 T Cell | $5.7 \times 10^{-1}$ |
| NK Cell | $8.7 \times 10^{-1}$ |
| Ly6C High Monocyte | $1.0 \times 10^{-1}$ |
| Ly6C Low Monocyte | $4.3 \times 10^{-10}$ |
| Eosinophil | $9.8 \times 10^{-12}$ |
| Neutrophil | $1.0 \times 10^{-1}$ |

B Cell

| Lipid class | **PL acyl chains** |
|---|---|
| Feature | 22:5 |
| One-way ANOVA *p-value* | $2.2 \times 10^{-35}$ |
| FDR adjuested *p-value* | $1.5 \times 10^{-34}$ |

| | |
|---|---|
| B Cell | # |
| CD4 T Cell | $3.2 \times 10^{-7}$ |
| CD8 T Cell | $9.7 \times 10^{-12}$ |
| NK Cell | $9.5 \times 10^{-12}$ |
| Ly6C High Monocyte | $1.0 \times 10^{-1}$ |
| Ly6C Low Monocyte | $3.4 \times 10^{-2}$ |
| Eosinophil | $6.3 \times 10^{-2}$ |
| Neutrophil | $1.3 \times 10^{-11}$ |

B Cell

| Lipid class | **PL acyl chains** |
|---|---|
| Feature | 22:6 |

| | |
|---|---|
| B Cell | # |

| | |
|---|---|
| One-way ANOVA *p-value* | $2.9 \times 10^{-26}$ |
| FDR adjuested *p-value* | $1.2 \times 10^{-25}$ |

| | B Cell |
|---|---|
| CD4 T Cell | $4.7 \times 10^{-1}$ |
| CD8 T Cell | $7.8 \times 10^{-1}$ |
| NK Cell | $8.4 \times 10^{-1}$ |
| Ly6C High Monocyte | $1.4 \times 10^{-11}$ |
| Ly6C Low Monocyte | $4.1 \times 10^{-8}$ |
| Eosinophil | $9.6 \times 10^{-12}$ |
| Neutrophil | $9.6 \times 10^{-12}$ |

**pholipid acyl chain composition in 8 different murine immune cells**

| CD4 T Cell | CD8 T Cell | NK Cell | Ly6C High Monocyte | Ly6C Low Monocyte | Eosinophil | Neutrophil |
|---|---|---|---|---|---|---|
| # | | | | | | |
| $1.5 \times 10^{-5}$ | # | | | | | |
| $1.5 \times 10^{-5}$ | $3.1 \times 10^{-1}$ | # | | | | |
| $1.9 \times 10^{-5}$ | $4.0 \times 10^{-1}$ | $1.0 \times 10^{-1}$ | # | | | |
| $2.0 \times 10^{-5}$ | $3.5 \times 10^{-1}$ | $1.0 \times 10^{-1}$ | $1.0 \times 10^{-1}$ | # | | |
| $1.3 \times 10^{-6}$ | $8.9 \times 10^{-2}$ | $1.0 \times 10^{-1}$ | $1.0 \times 10^{-1}$ | $1.0 \times 10^{-1}$ | # | |
| $9.1 \times 10^{-10}$ | $5.8 \times 10^{-4}$ | $3.1 \times 10^{-1}$ | $1.9 \times 10^{-1}$ | $2.8 \times 10^{-1}$ | $7.2 \times 10^{-1}$ | # |

| CD4 T Cell | CD8 T Cell | NK Cell | Ly6C High Monocyte | Ly6C Low Monocyte | Eosinophil | Neutrophil |
|---|---|---|---|---|---|---|
| # | | | | | | |
| $1.0 \times 10^{-1}$ | # | | | | | |
| $1.0 \times 10^{-1}$ | $1.0 \times 10^{-1}$ | # | | | | |
| $2.4 \times 10^{-3}$ | $5.3 \times 10^{-3}$ | $2.6 \times 10^{-2}$ | # | | | |
| $6.6 \times 10^{-1}$ | $7.6 \times 10^{-1}$ | $1.0 \times 10^{-1}$ | $2.8 \times 10^{-1}$ | # | | |
| $1.2 \times 10^{-11}$ | $2.3 \times 10^{-11}$ | $6.7 \times 10^{-11}$ | $1.7 \times 10^{-5}$ | $3.1 \times 10^{-9}$ | # | |
| $9.5 \times 10^{-12}$ | $9.5 \times 10^{-12}$ | $9.5 \times 10^{-12}$ | $9.5 \times 10^{-12}$ | $9.5 \times 10^{-12}$ | $9.5 \times 10^{-12}$ | # |

| | | | | | |
|---|---|---|---|---|---|
| # | | | | | |
| $1.4 \times 10^{-2}$ | # | | | | |
| $1.4 \times 10^{-2}$ | $1.0 \times 10^{-1}$ | # | | | |
| $2.1 \times 10^{-1}$ | $1.0 \times 10^{-1}$ | $9.4 \times 10^{-1}$ | # | | |
| $4.3 \times 10^{-1}$ | $1.0 \times 10^{-1}$ | $8.1 \times 10^{-1}$ | $1.0 \times 10^{-1}$ | # | |
| $8.5 \times 10^{-1}$ | $1.0 \times 10^{-1}$ | $3.8 \times 10^{-1}$ | $1.0 \times 10^{-1}$ | $1.0 \times 10^{-1}$ | # |

| CD4 T Cell | CD8 T Cell | NK Cell | Ly6C High Monocyte | Ly6C Low Monocyte | Eosinophil | Neutrophil |
|---|---|---|---|---|---|---|
| $5.2 \times 10^{-2}$ | $2.4 \times 10^{-5}$ | $1.2 \times 10^{-7}$ | $6.4 \times 10^{-6}$ | $4.7 \times 10^{-5}$ | $6.1 \times 10^{-4}$ | # |

| CD4 T Cell | CD8 T Cell | NK Cell | Ly6C High Monocyte | Ly6C Low Monocyte | Eosinophil | Neutrophil |
|---|---|---|---|---|---|---|
| # | | | | | | |
| $1.9 \times 10^{-2}$ | # | | | | | |
| $1.9 \times 10^{-2}$ | $1.0 \times 10^{-1}$ | # | | | | |
| $7.6 \times 10^{-1}$ | $4.9 \times 10^{-3}$ | $6.9 \times 10^{-5}$ | # | | | |
| $9.3 \times 10^{-2}$ | $7.1 \times 10^{-5}$ | $6.2 \times 10^{-7}$ | $8.6 \times 10^{-1}$ | # | | |
| $7.4 \times 10^{-1}$ | $1.0 \times 10^{-1}$ | $5.7 \times 10^{-1}$ | $3.9 \times 10^{-2}$ | $7.5 \times 10^{-4}$ | # | |
| $2.9 \times 10^{-2}$ | $1.1 \times 10^{-5}$ | $7.0 \times 10^{-8}$ | $6.1 \times 10^{-1}$ | $1.0 \times 10^{-1}$ | $1.3 \times 10^{-4}$ | # |

| CD4 T Cell | CD8 T Cell | NK Cell | Ly6C High Monocyte | Ly6C Low Monocyte | Eosinophil | Neutrophil |
|---|---|---|---|---|---|---|
| # | | | | | | |
| $1.0 \times 10^{-1}$ | # | | | | | |
| $1.0 \times 10^{-1}$ | $1.0 \times 10^{-1}$ | # | | | | |
| $1.0 \times 10^{-1}$ | $1.0 \times 10^{-1}$ | $1.0 \times 10^{-1}$ | # | | | |
| $2.2 \times 10^{-1}$ | $5.3 \times 10^{-1}$ | $2.1 \times 10^{-1}$ | $7.4 \times 10^{-1}$ | # | | |
| $2.8 \times 10^{-1}$ | $6.0 \times 10^{-1}$ | $2.6 \times 10^{-1}$ | $8.1 \times 10^{-1}$ | $1.0 \times 10^{-1}$ | # | |
| $1.0 \times 10^{-1}$ | $1.0 \times 10^{-1}$ | $1.0 \times 10^{-1}$ | $1.0 \times 10^{-1}$ | $4.7 \times 10^{-1}$ | $5.5 \times 10^{-1}$ | # |

| | CD4 T Cell | CD8 T Cell | NK Cell | Ly6C High Monocyte | Ly6C Low Monocyte | Eosinophil | Neutrophil |
|---|---|---|---|---|---|---|---|
| | # | | | | | | |
| | $1.0 \times 10^{-1}$ | # | | | | | |
| | $1.0 \times 10^{-1}$ | $1.0 \times 10^{-1}$ | # | | | | |
| | $1.3 \times 10^{-4}$ | $1.1 \times 10^{-3}$ | $1.2 \times 10^{-3}$ | # | | | |
| | $1.2 \times 10^{-2}$ | $5.0 \times 10^{-2}$ | $6.0 \times 10^{-2}$ | $9.2 \times 10^{-1}$ | # | | |
| | $1.0 \times 10^{-1}$ | $8.7 \times 10^{-1}$ | $7.7 \times 10^{-1}$ | $3.0 \times 10^{-6}$ | $4.7 \times 10^{-4}$ | # | |
| | $1.0 \times 10^{-1}$ | $1.0 \times 10^{-1}$ | $1.0 \times 10^{-1}$ | $4.0 \times 10^{-5}$ | $4.8 \times 10^{-3}$ | $1.0 \times 10^{-1}$ | # |

| | CD4 T Cell | CD8 T Cell | NK Cell | Ly6C High Monocyte | Ly6C Low Monocyte | Eosinophil | Neutrophil |
|---|---|---|---|---|---|---|---|
| | # | | | | | | |
| | $1.2 \times 10^{-1}$ | # | | | | | |
| | $1.2 \times 10^{-1}$ | $1.0 \times 10^{-1}$ | # | | | | |
| | $3.5 \times 10^{-1}$ | $1.0 \times 10^{-1}$ | $1.0 \times 10^{-1}$ | # | | | |
| | $1.0 \times 10^{-1}$ | $3.1 \times 10^{-1}$ | $8.0 \times 10^{-2}$ | $2.6 \times 10^{-1}$ | # | | |
| | $6.6 \times 10^{-4}$ | $3.3 \times 10^{-1}$ | $6.5 \times 10^{-1}$ | $2.5 \times 10^{-1}$ | $3.6 \times 10^{-4}$ | # | |
| | $7.7 \times 10^{-2}$ | $1.0 \times 10^{-4}$ | $5.8 \times 10^{-6}$ | $3.7 \times 10^{-5}$ | $1.2 \times 10^{-1}$ | $4.4 \times 10^{-9}$ | # |

| | CD4 T Cell | CD8 T Cell | NK Cell | Ly6C High Monocyte | Ly6C Low Monocyte | Eosinophil | Neutrophil |
|---|---|---|---|---|---|---|---|
| | # | | | | | | |
| | $8.0 \times 10^{-1}$ | # | | | | | |
| | $8.0 \times 10^{-1}$ | $1.0 \times 10^{-1}$ | # | | | | |
| | $2.0 \times 10^{-5}$ | $3.4 \times 10^{-6}$ | $4.3 \times 10^{-8}$ | # | | | |
| | $1.0 \times 10^{-3}$ | $1.9 \times 10^{-4}$ | $3.9 \times 10^{-6}$ | $1.0 \times 10^{-1}$ | # | | |
| | $9.6 \times 10^{-12}$ | $9.6 \times 10^{-12}$ | $9.6 \times 10^{-12}$ | $3.0 \times 10^{-5}$ | $1.3 \times 10^{-6}$ | # | |
| | $5.9 \times 10^{-4}$ | $1.0 \times 10^{-4}$ | $1.7 \times 10^{-6}$ | $1.0 \times 10^{-1}$ | $1.0 \times 10^{-1}$ | $8.2 \times 10^{-7}$ | # |

| | CD4 T Cell | CD8 T Cell | NK Cell | Ly6C High Monocyte | Ly6C Low Monocyte | Eosinophil | Neutrophil |
|---|---|---|---|---|---|---|---|
| CD4 T Cell | # | | | | | | |
| CD8 T Cell | $1.0 \times 10^{-1}$ | # | | | | | |
| NK Cell | $1.0 \times 10^{-1}$ | $7.3 \times 10^{-1}$ | # | | | | |
| Ly6C High Monocyte | $3.4 \times 10^{-5}$ | $2.7 \times 10^{-6}$ | $7.9 \times 10^{-4}$ | # | | | |
| Ly6C Low Monocyte | $8.8 \times 10^{-4}$ | $7.8 \times 10^{-5}$ | $1.3 \times 10^{-2}$ | $1.0 \times 10^{-1}$ | # | | |
| Eosinophil | $8.9 \times 10^{-1}$ | $1.0 \times 10^{-1}$ | $3.9 \times 10^{-1}$ | $1.8 \times 10^{-7}$ | $6.9 \times 10^{-6}$ | # | |
| Neutrophil | $2.1 \times 10^{-1}$ | $3.6 \times 10^{-2}$ | $7.2 \times 10^{-1}$ | $8.6 \times 10^{-2}$ | $4.5 \times 10^{-1}$ | $6.3 \times 10^{-3}$ | # |

| | CD4 T Cell | CD8 T Cell | NK Cell | Ly6C High Monocyte | Ly6C Low Monocyte | Eosinophil | Neutrophil |
|---|---|---|---|---|---|---|---|
| CD4 T Cell | # | | | | | | |
| CD8 T Cell | $1.0 \times 10^{-1}$ | # | | | | | |
| NK Cell | $1.0 \times 10^{-1}$ | $1.0 \times 10^{-1}$ | # | | | | |
| Ly6C High Monocyte | $2.4 \times 10^{-1}$ | $1.8 \times 10^{-1}$ | $3.6 \times 10^{-1}$ | # | | | |
| Ly6C Low Monocyte | $1.0 \times 10^{-1}$ | $1.0 \times 10^{-1}$ | $1.0 \times 10^{-1}$ | $6.9 \times 10^{-1}$ | # | | |
| Eosinophil | $1.0 \times 10^{-1}$ | $1.0 \times 10^{-1}$ | $1.0 \times 10^{-1}$ | $3.1 \times 10^{-1}$ | $1.0 \times 10^{-1}$ | # | |
| Neutrophil | $9.5 \times 10^{-12}$ | $9.5 \times 10^{-12}$ | $9.5 \times 10^{-12}$ | $9.5 \times 10^{-12}$ | $9.5 \times 10^{-12}$ | $9.5 \times 10^{-12}$ | # |

| CD4 T Cell | CD8 T Cell | NK Cell | Ly6C High Monocyte | Ly6C Low Monocyte | Eosinophil | Neutrophil |
|---|---|---|---|---|---|---|
| # | | | | | | |
| $9.4 \times 10^{-1}$ | # | | | | | |
| $9.4 \times 10^{-1}$ | $1.0 \times 10^{-1}$ | # | | | | |
| $1.0 \times 10^{-1}$ | $1.0 \times 10^{-1}$ | $1.0 \times 10^{-1}$ | # | | | |
| $2.6 \times 10^{-2}$ | $5.7 \times 10^{-1}$ | $3.7 \times 10^{-1}$ | $2.6 \times 10^{-1}$ | # | | |
| $7.7 \times 10^{-4}$ | $7.9 \times 10^{-2}$ | $3.1 \times 10^{-2}$ | $1.7 \times 10^{-2}$ | $1.0 \times 10^{-1}$ | # | |
| $2.0 \times 10^{-11}$ | $1.1 \times 10^{-8}$ | $1.4 \times 10^{-9}$ | $3.5 \times 10^{-10}$ | $1.2 \times 10^{-5}$ | $7.6 \times 10^{-4}$ | # |

| CD4 T Cell | CD8 T Cell | NK Cell | Ly6C High Monocyte | Ly6C Low Monocyte | Eosinophil | Neutrophil |
|---|---|---|---|---|---|---|
| # | | | | | | |
| $1.0 \times 10^{-1}$ | # | | | | | |
| $1.0 \times 10^{-1}$ | $8.1 \times 10^{-1}$ | # | | | | |
| $6.0 \times 10^{-1}$ | $3.1 \times 10^{-1}$ | $1.0 \times 10^{-1}$ | # | | | |
| $1.0 \times 10^{-1}$ | $1.0 \times 10^{-1}$ | $1.0 \times 10^{-1}$ | $7.5 \times 10^{-1}$ | # | | |
| $2.2 \times 10^{-1}$ | $5.5 \times 10^{-1}$ | $1.8 \times 10^{-2}$ | $1.1 \times 10^{-3}$ | $1.3 \times 10^{-1}$ | # | |
| $9.4 \times 10^{-7}$ | $1.5 \times 10^{-5}$ | $1.4 \times 10^{-8}$ | $2.4 \times 10^{-10}$ | $3.6 \times 10^{-7}$ | $9.5 \times 10^{-3}$ | # |

| | | | | | | |
|---|---|---|---|---|---|---|
| # | | | | | | |
| $1.0 \times 10^{-1}$ | # | | | | | |
| $1.0 \times 10^{-1}$ | $1.0 \times 10^{-1}$ | # | | | | |
| $1.7 \times 10^{-10}$ | $4.2 \times 10^{-11}$ | $7.4 \times 10^{-10}$ | # | | | |
| $8.3 \times 10^{-1}$ | $4.5 \times 10^{-1}$ | $1.0 \times 10^{-1}$ | $6.7 \times 10^{-8}$ | # | | |
| $2.6 \times 10^{-7}$ | $4.2 \times 10^{-8}$ | $1.1 \times 10^{-6}$ | $7.2 \times 10^{-1}$ | $7.1 \times 10^{-5}$ | # | |
| $1.5 \times 10^{-11}$ | $1.1 \times 10^{-11}$ | $3.3 \times 10^{-11}$ | $1.0 \times 10^{-1}$ | $2.1 \times 10^{-9}$ | $2.1 \times 10^{-1}$ | # |

CD4 T Cell | CD8 T Cell | NK Cell | Ly6C High Monocyte | Ly6C Low Monocyte | Eosinophil | Neutrophil

| CD4 T Cell | CD8 T Cell | NK Cell | Ly6C High Monocyte | Ly6C Low Monocyte | Eosinophil | Neutrophil |
|---|---|---|---|---|---|---|
| # | | | | | | |
| $5.9 \times 10^{-1}$ | # | | | | | |
| $5.9 \times 10^{-1}$ | $1.0 \times 10^{-1}$ | # | | | | |
| $5.4 \times 10^{-1}$ | $1.0 \times 10^{-1}$ | $1.0 \times 10^{-1}$ | # | | | |
| $6.8 \times 10^{-1}$ | $1.0 \times 10^{-1}$ | $1.0 \times 10^{-1}$ | $1.0 \times 10^{-1}$ | # | | |
| $1.0 \times 10^{-1}$ | $3.0 \times 10^{-2}$ | $9.4 \times 10^{-2}$ | $7.5 \times 10^{-2}$ | $1.3 \times 10^{-1}$ | # | |
| $1.1 \times 10^{-2}$ | $3.4 \times 10^{-6}$ | $1.4 \times 10^{-5}$ | $7.1 \times 10^{-6}$ | $2.3 \times 10^{-5}$ | $1.7 \times 10^{-1}$ | # |

| CD4 T Cell | CD8 T Cell | NK Cell | Ly6C High Monocyte | Ly6C Low Monocyte | Eosinophil | Neutrophil |
|---|---|---|---|---|---|---|
| # | | | | | | |
| $1.0 \times 10^{-1}$ | # | | | | | |
| $1.0 \times 10^{-1}$ | $1.0 \times 10^{-1}$ | # | | | | |
| $9.0 \times 10^{-1}$ | $9.1 \times 10^{-1}$ | $7.4 \times 10^{-1}$ | # | | | |
| $7.3 \times 10^{-3}$ | $1.0 \times 10^{-2}$ | $2.6 \times 10^{-3}$ | $1.7 \times 10^{-1}$ | # | | |
| $1.0 \times 10^{-1}$ | $1.0 \times 10^{-1}$ | $1.0 \times 10^{-1}$ | $7.8 \times 10^{-1}$ | $3.1 \times 10^{-3}$ | # | |
| $1.0 \times 10^{-1}$ | $1.0 \times 10^{-1}$ | $1.0 \times 10^{-1}$ | $6.6 \times 10^{-1}$ | $1.4 \times 10^{-3}$ | $1.0 \times 10^{-1}$ | # |

| CD4 T Cell | CD8 T Cell | NK Cell | Ly6C High Monocyte | Ly6C Low Monocyte | Eosinophil | Neutrophil |
|---|---|---|---|---|---|---|
| # | | | | | | |
| 1.5 x $10^{-1}$ | # | | | | | |
| 1.5 x $10^{-1}$ | 3.4 x $10^{-1}$ | # | | | | |
| 2.3 x $10^{-1}$ | 4.7 x $10^{-1}$ | 1.0 x $10^{-1}$ | # | | | |
| 1.0 x $10^{-1}$ | 1.0 x $10^{-1}$ | 5.1 x $10^{-1}$ | 6.6 x $10^{-1}$ | # | | |
| 2.0 x $10^{-1}$ | 1.1 x $10^{-1}$ | 5.7 x $10^{-5}$ | 9.8 x $10^{-5}$ | 3.7 x $10^{-2}$ | # | |
| 3.0 x $10^{-11}$ | 2.4 x $10^{-11}$ | 9.6 x $10^{-12}$ | 9.6 x $10^{-12}$ | 1.1 x $10^{-11}$ | 7.1 x $10^{-7}$ | # |

| CD4 T Cell | CD8 T Cell | NK Cell | Ly6C High Monocyte | Ly6C Low Monocyte | Eosinophil | Neutrophil |
|---|---|---|---|---|---|---|
| # | | | | | | |
| 8.4 x $10^{-11}$ | # | | | | | |
| 8.4 x $10^{-11}$ | 4.7 x $10^{-8}$ | # | | | | |
| 9.5 x $10^{-12}$ | 9.5 x $10^{-12}$ | 9.6 x $10^{-12}$ | # | | | |
| 9.5 x $10^{-12}$ | 9.5 x $10^{-12}$ | 1.7 x $10^{-5}$ | 2.6 x $10^{-5}$ | # | | |
| 9.5 x $10^{-12}$ | 9.5 x $10^{-12}$ | 9.5 x $10^{-12}$ | 2.7 x $10^{-10}$ | 9.5 x $10^{-12}$ | # | |
| 9.6 x $10^{-12}$ | 9.8 x $10^{-12}$ | 7.1 x $10^{-2}$ | 2.6 x $10^{-10}$ | 1.5 x $10^{-1}$ | 9.5 x $10^{-12}$ | # |

| | | | | | | |
|---|---|---|---|---|---|---|
| # | | | | | | |
| 2.0 x $10^{-3}$ | # | | | | | |
| 2.0 x $10^{-3}$ | 4.8 x $10^{-3}$ | # | | | | |
| 1.0 x $10^{-1}$ | 1.0 x $10^{-1}$ | 1.9 x $10^{-2}$ | # | | | |
| 5.9 x $10^{-1}$ | 7.1 x $10^{-1}$ | 2.8 x $10^{-1}$ | 1.0 x $10^{-1}$ | # | | |
| 7.0 x $10^{-4}$ | 1.8 x $10^{-3}$ | 1.0 x $10^{-1}$ | 7.5 x $10^{-3}$ | 1.5 x $10^{-1}$ | # | |
| 1.3 x $10^{-9}$ | 6.3 x $10^{-9}$ | 1.4 x $10^{-2}$ | 1.7 x $10^{-8}$ | 2.6 x $10^{-6}$ | 3.3 x $10^{-2}$ | # |

CD4 T Cell | CD8 T Cell | NK Cell | Ly6C High Monocyte | Ly6C Low Monocyte | Eosinophil | Neutrophil

| CD4 T Cell | CD8 T Cell | NK Cell | Ly6C High Monocyte | Ly6C Low Monocyte | Eosinophil | Neutrophil |
|---|---|---|---|---|---|---|
| # | | | | | | |
| $2.6 \times 10^{-1}$ | # | | | | | |
| $2.6 \times 10^{-1}$ | $1.0 \times 10^{-1}$ | # | | | | |
| $5.5 \times 10^{-1}$ | $1.0 \times 10^{-1}$ | $1.0 \times 10^{-1}$ | # | | | |
| $1.2 \times 10^{-11}$ | $1.0 \times 10^{-6}$ | $4.7 \times 10^{-8}$ | $2.4 \times 10^{-9}$ | # | | |
| $9.6 \times 10^{-12}$ | $4.9 \times 10^{-10}$ | $2.6 \times 10^{-11}$ | $1.0 \times 10^{-11}$ | $5.1 \times 10^{-1}$ | # | |
| $5.4 \times 10^{-1}$ | $2.3 \times 10^{-4}$ | $1.1 \times 10^{-3}$ | $4.8 \times 10^{-3}$ | $9.6 \times 10^{-12}$ | $9.5 \times 10^{-12}$ | # |

| CD4 T Cell | CD8 T Cell | NK Cell | Ly6C High Monocyte | Ly6C Low Monocyte | Eosinophil | Neutrophil |
|---|---|---|---|---|---|---|
| # | | | | | | |
| $1.1 \times 10^{-10}$ | # | | | | | |
| $1.1 \times 10^{-10}$ | $3.0 \times 10^{-3}$ | # | | | | |
| $1.9 \times 10^{-9}$ | $9.6 \times 10^{-12}$ | $9.5 \times 10^{-12}$ | # | | | |
| $2.6 \times 10^{-2}$ | $7.9 \times 10^{-9}$ | $9.6 \times 10^{-12}$ | $1.2 \times 10^{-3}$ | # | | |
| $1.0 \times 10^{-11}$ | $9.5 \times 10^{-12}$ | $9.5 \times 10^{-12}$ | $2.9 \times 10^{-1}$ | $3.6 \times 10^{-7}$ | # | |
| $9.5 \times 10^{-12}$ | $9.5 \times 10^{-12}$ | $9.5 \times 10^{-12}$ | $2.2 \times 10^{-11}$ | $9.5 \times 10^{-12}$ | $3.8 \times 10^{-7}$ | # |

| CD4 T Cell | CD8 T Cell | NK Cell | Ly6C High Monocyte | Ly6C Low Monocyte | Eosinophil | Neutrophil |
|---|---|---|---|---|---|---|
| # | | | | | | |
| $1.5 \times 10^{-2}$ | # | | | | | |
| $1.5 \times 10^{-2}$ | $1.0 \times 10^{-1}$ | # | | | | |
| $7.0 \times 10^{-9}$ | $9.6 \times 10^{-12}$ | $9.6 \times 10^{-12}$ | # | | | |
| $6.4 \times 10^{-5}$ | $7.6 \times 10^{-11}$ | $5.4 \times 10^{-11}$ | $3.8 \times 10^{-1}$ | # | | |
| $4.5 \times 10^{-11}$ | $9.6 \times 10^{-12}$ | $9.5 \times 10^{-12}$ | $7.9 \times 10^{-1}$ | $1.3 \times 10^{-2}$ | # | |
| $1.1 \times 10^{-11}$ | $9.5 \times 10^{-12}$ | $9.5 \times 10^{-12}$ | $3.5 \times 10^{-1}$ | $1.2 \times 10^{-3}$ | $1.0 \times 10^{-1}$ | # |

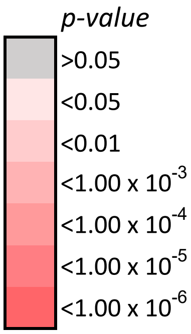
p-value
>0.05
<0.05
<0.01
<1.00 x 10-3
<1.00 x 10-4
<1.00 x 10-5
<1.00 x 10-6

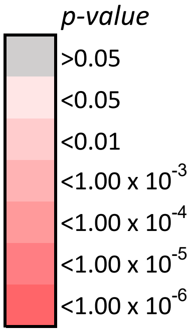
p-value
>0.05
<0.05
<0.01
<1.00 x 10-3
<1.00 x 10-4
<1.00 x 10-5
<1.00 x 10-6

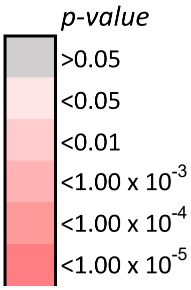
p-value
>0.05
<0.05
<0.01
<1.00 x 10-3
<1.00 x 10-4
<1.00 x 10-5

<$1.00 \times 10^{-6}$

*p-value*

>0.05

<0.05

<0.01

<$1.00 \times 10^{-3}$

<$1.00 \times 10^{-4}$

<$1.00 \times 10^{-5}$

<$1.00 \times 10^{-6}$

*p-value*

>0.05

<0.05

<0.01

<$1.00 \times 10^{-3}$

<$1.00 \times 10^{-4}$

<$1.00 \times 10^{-5}$

<$1.00 \times 10^{-6}$

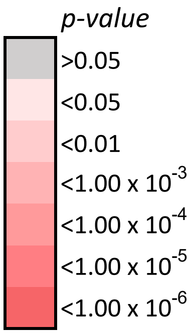
p-value
>0.05
<0.05
<0.01
<1.00 x 10-3
<1.00 x 10-4
<1.00 x 10-5
<1.00 x 10-6

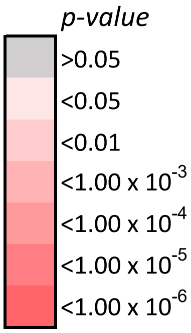
p-value
>0.05
<0.05
<0.01
<1.00 x 10-3
<1.00 x 10-4
<1.00 x 10-5
<1.00 x 10-6

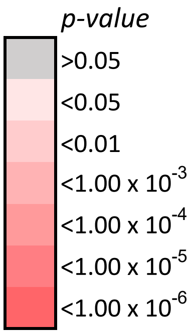
p-value
>0.05
<0.05
<0.01
<1.00 x 10-3
<1.00 x 10-4
<1.00 x 10-5
<1.00 x 10-6

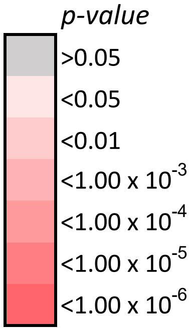
*p-value*
>0.05
<0.05
<0.01
<1.00 x 10-3
<1.00 x 10-4
<1.00 x 10-5
<1.00 x 10-6

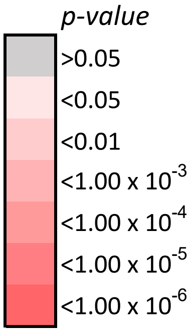
*p-value*
>0.05
<0.05
<0.01
<1.00 x 10-3
<1.00 x 10-4
<1.00 x 10-5
<1.00 x 10-6

*p-value*

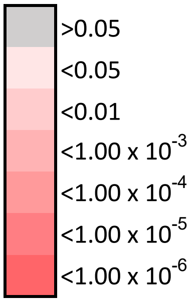
>0.05
<0.05
<0.01
$<1.00 \times 10^{-3}$
$<1.00 \times 10^{-4}$
$<1.00 \times 10^{-5}$
$<1.00 \times 10^{-6}$

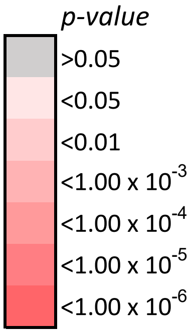
*p-value*
>0.05
<0.05
<0.01
$<1.00 \times 10^{-3}$
$<1.00 \times 10^{-4}$
$<1.00 \times 10^{-5}$
$<1.00 \times 10^{-6}$

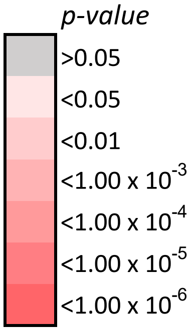
*p-value*
>0.05
<0.05
<0.01
$<1.00 \times 10^{-3}$
$<1.00 \times 10^{-4}$
$<1.00 \times 10^{-5}$
$<1.00 \times 10^{-6}$

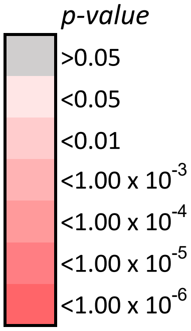
p-value
>0.05
<0.05
<0.01
<1.00 x 10-3
<1.00 x 10-4
<1.00 x 10-5
<1.00 x 10-6

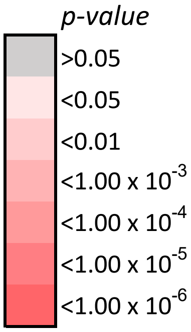
p-value
>0.05
<0.05
<0.01
<1.00 x 10-3
<1.00 x 10-4
<1.00 x 10-5
<1.00 x 10-6

*p-value*

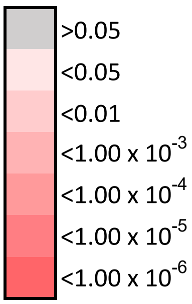
>0.05
<0.05
<0.01
<1.00 x 10-3
<1.00 x 10-4
<1.00 x 10-5
<1.00 x 10-6

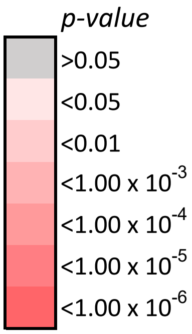
p-value
>0.05
<0.05
<0.01
<1.00 x 10-3
<1.00 x 10-4
<1.00 x 10-5
<1.00 x 10-6

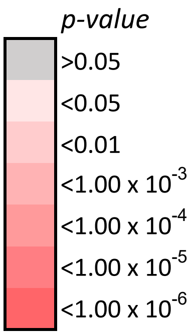
p-value
>0.05
<0.05
<0.01
<1.00 x 10-3
<1.00 x 10-4
<1.00 x 10-5
<1.00 x 10-6

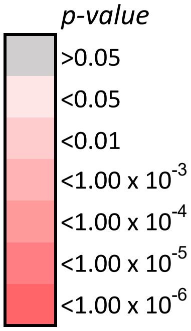
*p-value*
>0.05
<0.05
<0.01
<1.00 x $10^{-3}$
<1.00 x $10^{-4}$
<1.00 x $10^{-5}$
<1.00 x $10^{-6}$

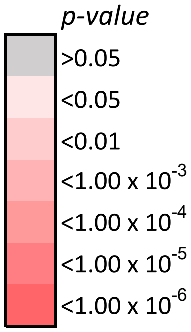
*p-value*
>0.05
<0.05
<0.01
<1.00 x $10^{-3}$
<1.00 x $10^{-4}$
<1.00 x $10^{-5}$
<1.00 x $10^{-6}$

*p-value*

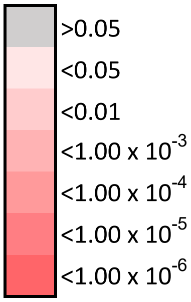
>0.05
<0.05
<0.01
<1.00 x 10^-3
<1.00 x 10^-4
<1.00 x 10^-5
<1.00 x 10^-6
